# Targeted Transcriptional Repression by Induced Proximity

**DOI:** 10.1101/2025.10.22.680877

**Authors:** Christian E. Stieger, Xinru Chen, Dustin Dovala, Fabian Wu, Nicolas Pizzato, Jeffrey McKenna, Cory Johannessen, Barna D. Fodor, Markus Schirle, Daniel K. Nomura

## Abstract

Most cancer-driving proteins remain “undruggable” due to the absence of ligandable pockets and their reliance on intrinsically disordered or protein–DNA/protein–protein interactions. Transcription factors, which orchestrate oncogenic gene expression programs, are particularly challenging: they turn over rapidly, evade durable pharmacological inhibition, and resist even emerging targeted protein degradation strategies. Here, we describe a new induced-proximity therapeutic modality, **T**ranscriptional **R**egulation via **A**ctive **C**ontrol of **E**pigenetic **R**eprogramming (TRACER), that enforces locus-specific transcriptional silencing by recruiting endogenous corepressor complexes to transcription factor binding sites. We developed small-molecule TRACERs that tether methyl-CpG binding domain protein 2 (MBD2) and the Nucleosome Remodeling and Deacetylase (NuRD) complex to transcription factor–directed ligands. An estrogen receptor (ER) TRACER potently suppressed ER transcriptional activity in breast cancer cells, downregulated canonical ER target genes, and required MBD2 and histone deacetylase (HDAC1/2) for activity, confirming on-target epigenetic repression. Extending this approach to prostate cancer, an androgen receptor (AR) TRACER transcriptionally repressed both full-length AR and the drug-resistant truncation variant AR-V7, achieving >90% inhibition of AR transcriptional activity in androgen-independent prostate cancer cells with locus-specific gene repression. We further show that TRACERs can be modularly reprogrammed to recruit alternative repressors, including PRC2. Collectively, these findings establish TRACERs as a generalizable modality to pharmacologically silence undruggable transcription factors through targeted epigenetic reprogramming, offering a powerful new strategy for treating cancers refractory to existing therapies.

## Introduction

Despite the discovery of so many potential therapeutic targets for cancer, a critical bottleneck in realizing new cancer cures that exploit our expanding knowledge of cancer is the daunting realization that most proteins, over 90 %, are still considered “undruggable.”^1^ This is because most proteins do not possess classical binding pockets often found in enzyme active sites, but rather function through protein-protein or protein-nucleic acid interactions, or are highly intrinsically disordered or unstructured. Even with advances in innovative technologies for mapping ligandable sites and approaches for enabling ligand discovery against these undruggable targets, we still lack the complementary arsenal of therapeutic modalities for functionally, durably, and safely exploiting these therapeutic targets to combat cancer, while mitigating toxicity in patients.

One of the most challenging and undruggable, yet therapeutically critical, classes of proteins known to drive many human cancers is transcription factors ^2,3^. Transcription factors, such as MYC, CTNNB1, androgen receptors (AR and truncation variant AR-V7), estrogen receptors (ER), YAP and TAZ, STAT3, and many others, are known to drive cancer pathogenesis through engaging their respective promoters at genomic loci to activate the transcription of genes involved in cell proliferation, survival, and metastasis. While some of these transcription factors have been pharmacologically targeted and even degraded with newer therapeutic modalities such as targeted protein degradation (TPD) (e.g. ER and AR)^4^, many of these transcription factors have been more undruggable and difficult to target directly. Furthermore, transcription factors present an additional challenge in that many of these proteins have very rapid turnover rates, particularly in cancer cells, where they are rapidly degraded and resynthesized, making them challenging to target with classical inhibitors or even innovative degraders in a durable manner. Efforts to epigenetically target these transcription factors by broadly targeting chromatin remodelling through the modulation of histone modifications (e.g., HDAC1/2, CBP/EP300, SMARCA2/4, and BRD4 inhibitors and degraders^5–7^) have thus far not shown an ideal therapeutic window.

An ideal therapeutic strategy would be to specifically and directly target specific transcription factors and epigenetically silence genetic loci associated with those transcription factors, rather than broadly targeting multiple transcriptional loci. While epigenome editing using dCas9/gRNA fusions with DNA methyltransferases has enabled locus-specific epigenetic silencing, these large protein-nucleic acid complexes are not practical for systemic and comprehensive delivery to most cancer types ^8^. Recently, the Gray and Crabtree groups have reported on a small-molecule based targeted transcriptional activation or rewiring approaches with transcriptional/epigenetic chemical inducers of proximity (TCIPs) wherein they developed heterobifunctional molecules to recruit BRD4, CDK9, or lysine acetyltransferases to BCL6 transcriptional loci to activate BCL6 genetic loci and repress MYC ^9–11^. In this study, we have developed a novel induced proximity-based therapeutic modality for targeted transcriptional repression, termed Transcriptional Regulation via Active Control of Epigenetic Reprogramming (TRACER). TRACERs consist of a transcriptional corepressor complex recruiter linked to a transcription factor targeting ligand to epigenetically and locus-specifically repress transcription. We demonstrate proof-of-concept of the TRACER platform through the recruitment of methyl CpG binding domain protein 2 (MBD2) and the nucleosome remodelling and deacetylase (NuRD) corepressor complex to estrogen receptor (ER) and androgen receptor (AR) genetic loci in breast and prostate cancer cells, respectively.

## Results

### Characterization of an MBD2 inhibitor as an MBD2 recruiter for TRACER applications

MBD2 functions as a core subunit of the NuRD complex, where it binds methylated DNA and recruits other NuRD components—including histone deacetylases (HDAC1/2), CHD3/4, RBBP4/7, and MTA family proteins--to coordinate chromatin remodelling and histone deacetylation, ultimately leading to transcriptional repression ^12^. MBD2 may also bring in additional repressors, including PRMT5 and SIN3A-associated corepressor complexes, although these interactions are more context-specific and less universally conserved than NuRD ^13^. To enable the TRACER strategy, we hypothesized that we could utilize MBD2 as a general recruiter of corepressor complexes to facilitate targeted transcriptional repression. We identified a previously reported MBD2 ligand, KCC-07, that is able to block the interaction between MBD2 and hypermethylated DNA both *in vitro* and *in* situ as a promising scaffold to develop TRACERs **(Figure 1a)** ^14–16^. We recapitulated the functional MBD2 binding assay with a fluorescent methylated CpG-containing oligonucleotide by fluorescence polarization and the reported inhibitory activity of the MBD2 ligand. The structurally related compound CS-1-69, as well as linker-modified variants, still maintained MBD2 inhibitory activity **(Figure S1a, Figure 1b-1c)**. We also prepared a photoaffinity probe to demonstrate MBD2 binding with the probe and displacement of binding with excess KCC-07 **(Figure 1d-1f)**.

**Figure 1.**
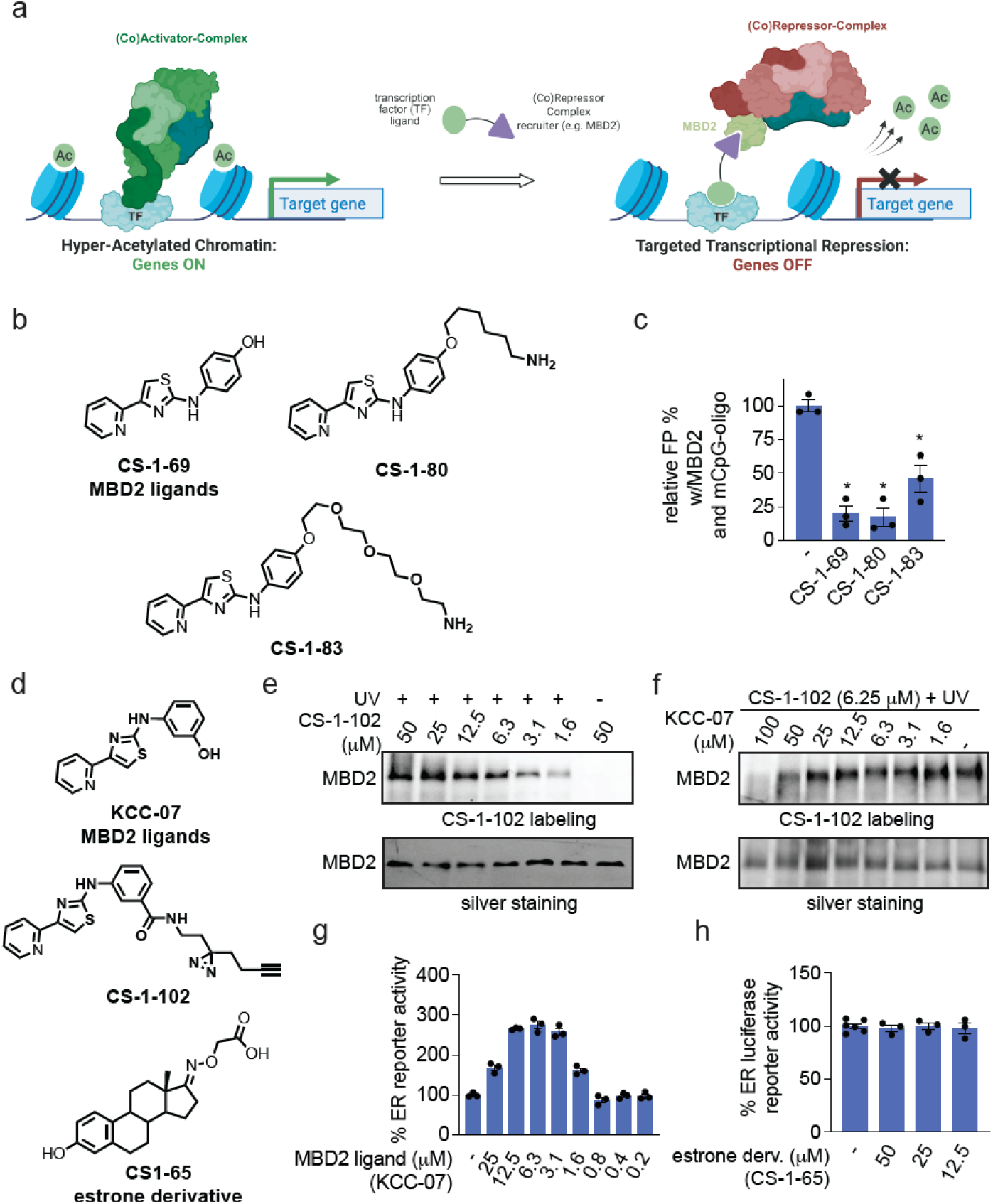
Targeted Transcriptional Repression with TRACERs using an MBD2 recruiter. **(a)** Targeted transcriptional repression with TRACERs that consist of an epigenetic recruiter (e.g. against MBD2) linked to a transcription factor targeting ligand to recruit a corepressor complex to specific transcriptional loci to repress transcription. **(b)** Structures of MBD2 inhibitors and linkered analogs. **(c)** Characterization of MBD2 inhibitors. Pure MBD2 protein (150 nM) was incubated with a fluorescent methylated CpG oligonucleotide (20 nM) in the presence of ligand (50 μM) for 30 min, and MBD2 interaction with the mCpG-oligo was determined by fluorescence polarization. **(d)** Structures of additional MBD2 ligand KCC-07 and photoaffinity probe CS-1-102, and ER agonist derivative CS1-65. **(e)** Photoaffinity labeling of pure MBD2 protein. Pure MBD2 protein in **(e)** was treated with CS-1-102 for 30 min followed by photocrosslinking and subsequently subjected to azide-alkyne cycloaddition (CuAAC)-mediated appendage of a rhodamine-functionalized azide, after which protein labeling was read out by SDS/PAGE and in-gel fluorescence and loading was assessed by silver staining. **(f)** Competition of CS-1-102 labeling by KCC-07. The experiment was performed as described in **(e)** except that the MBD2 protein was pre-incubated with DMSO vehicle or KCC-07 for 30 min. **(g**,**h)** ER luciferase reporter activity in T47D cells treated with KCC-07 or CS-1-65 for 24 h. Data shown in **(e-h)** are from n=3 biologically independent replicates per group. Gels in **(e**,**f)** are representative. Data in **(g**,**h)** are individual replicate values and average ± sem.

To show initial proof-of-concept of the TRACER platform, we chose estrogen receptor (ER) as a target to pursue in breast cancer cells using an ER agonist, estrone. Given the precedence that functionalized estrogens can retain the ability to bind and even activate ER, we sought to use the oxime-linked estrone derivative CS-1-65 as an ER-ligand ^10,17^. Analogously to two naturally occurring estrogens, estradiol and estrone, the linker-functionalized estrone-derivative CS-1-96 was able to induce ER-mediated transcription, indicating good agonistic activity (**Figure S1b**). Furthermore, we demonstrated that the MBD2 inhibitor KCC-07 alone also stimulated ER transcriptional activity at higher concentrations **(Figure 1g-1h)**.

### Development and characterization of an ER TRACER

We subsequently synthesized heterobifunctional TRACERs against ER containing a variety of different linkers and showed that many of these molecules robustly and potently inhibited ER transcriptional activity in T47D breast cancer cells with overall minimal impairments in cell viability **(Figure S2a-S2b, Figure 2a-2d, Figure S3a-S3e)**. Among these compounds, CS-1-103, CS-1-163, and CS-1-86 showed the best potency, with 50% effective concentrations of 0.36, 0.89, and 1.3 μM, respectively **(Figures 2a-2d, S3a-S3e)**. The inhibition of ER transcriptional activity by the more potent compound CS-1-103, observed after 24 h of treatment, was even more pronounced after 48 and 72 h of treatment **(Figure 2e-2f)**. Transcriptional inhibitory activity was also still observed after a 12-hour washout of CS-1-103 following a 24-hour initial treatment, demonstrating the durability of the effect **(Figure 2g)**. To demonstrate the necessity of the MBD2 recruiter in the observed activity, we showed complete attenuation of CS-1-103 and CS-1-86-mediated ER inhibitory activity with KCC-07 or CS-1-69 cotreatment **(Figure 3a, Figure S4a)**. To demonstrate on-target activity through recruitment of HDACs and the NuRD complex, we also showed significant attenuation of CS-1-103 and CS-1-86-mediated ER inhibition by HDAC1/2-selective inhibitor Mocetinostat of CI-944 **(Figure 3b; Figure S4b)**. Most importantly, we demonstrated significant rescue of CS-1-103 and CS-1-86-mediated ER inhibitory activity upon MBD2 knockdown, demonstrating on-target activity **(Figure 3c-3d; Figure S4c)**. While these data were promising, our readout was using an artificial luciferase reporter system, and thus, we sought to confirm this ER transcriptional inhibitory activity with native ER gene targets. We showed that CS-1-103 treatment significantly downregulated ER target genes, including ER (ESR1) itself and GREB1, both under basal conditions with serum containing native steroid levels as well as under charcoal stripped conditions depleting native steroids and adding back exogenous estrogen E2 **(Figure 4a-4b)**. We also observed that ER protein levels were downregulated by CS-1-86 treatment, but we observed that the ER transcriptional activity was inhibited earlier than ER protein level reductions in a time-course study, indicating that the loss of ER protein levels was likely due to transcriptional inhibition and ER transcriptional downregulation rather than ER degradation **(Figure S4d-S4f)**. To further confirm that the CS-1-103-mediated transcriptional inhibitory activity was not occurring through proteasomal degradation-mediated pathways, we further demonstrated that the CS-1-103-mediated ER transcriptional inhibitory activity was not attenuated with pre-treatment of cells with a proteasome inhibitor, bortezomib **(Figure S4g)**. In accordance with our observations in the luciferase reporter system, we further demonstrated significant attenuation of ER target gene downregulation upon MBD2 knockdown while tamoxifen-mediated inhibition of ER target genes, was not attenuated by MBD2 knockdown **(Figure 4c-4d, Figure S4h)**. Quantitative proteomics revealed a significant reduction in ER target protein abundance following 24 h of treatment **(Figure 4e)**. Transcriptomic profiling by RNAseq showed robust repression of various ER target genes and revealed high selectivity for genes associated with estrogen-response **(Figure 4f, S5a)**. Moreover, the comparison of the RNAseq data with publicly available transcription factor ChIPseq experiments revealed a strong enrichment for ESR1 and ESR2 datasets **(Figure 4g, S5b)**.

**Figure 2.**
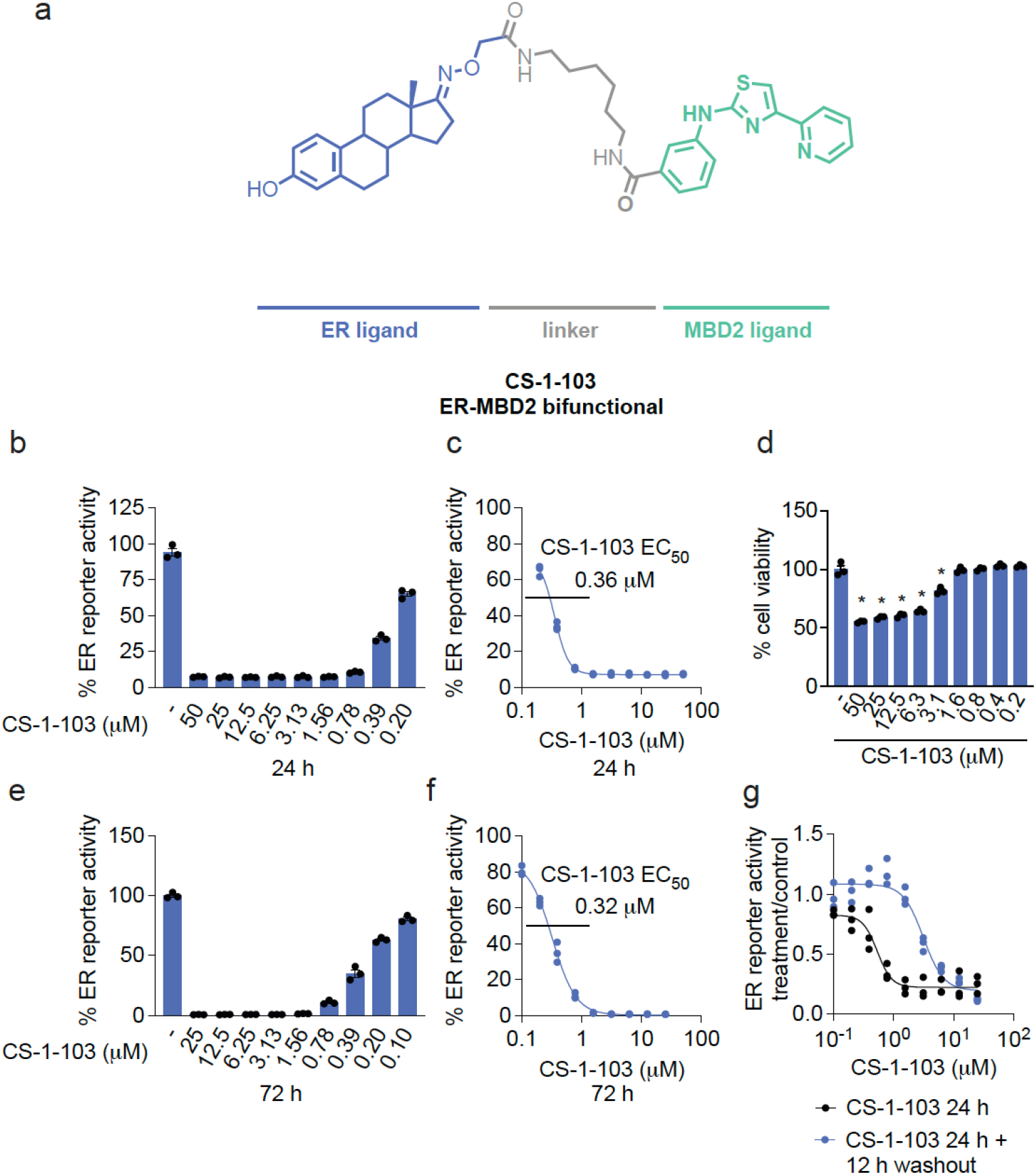
ER TRACER CS-1-103. **(a)** Structure of ER TRACER CS-1-103. **(b**,**c)** Dose-responsive inhibition of ER luciferase transcriptional reporter activity in T47D cells upon treatment with DMSO vehicle or TRACER CS-1-103 for 24 h, showing an EC_50_ of 360 nM. **(d)** 24 h cell viability by Cell TiterGlo in T47D cells. **(e**,**f)** Inhibition of T47D ER luciferase reporter activity after treatment with DMSO vehicle or CS-1-103 treatment for 72 h, showing an EC_50_ of 320 nM. **(g)** Dose-response of ER luciferase transcriptional reporter inhibition in T47D cells from treatment of cells with DMSO vehicle or CS-1-103 for 24 h or 24 h and then 12 hours of compound washout. Shown in **(b-g)** are individual replicate values and average ± sem.

**Figure 3.**
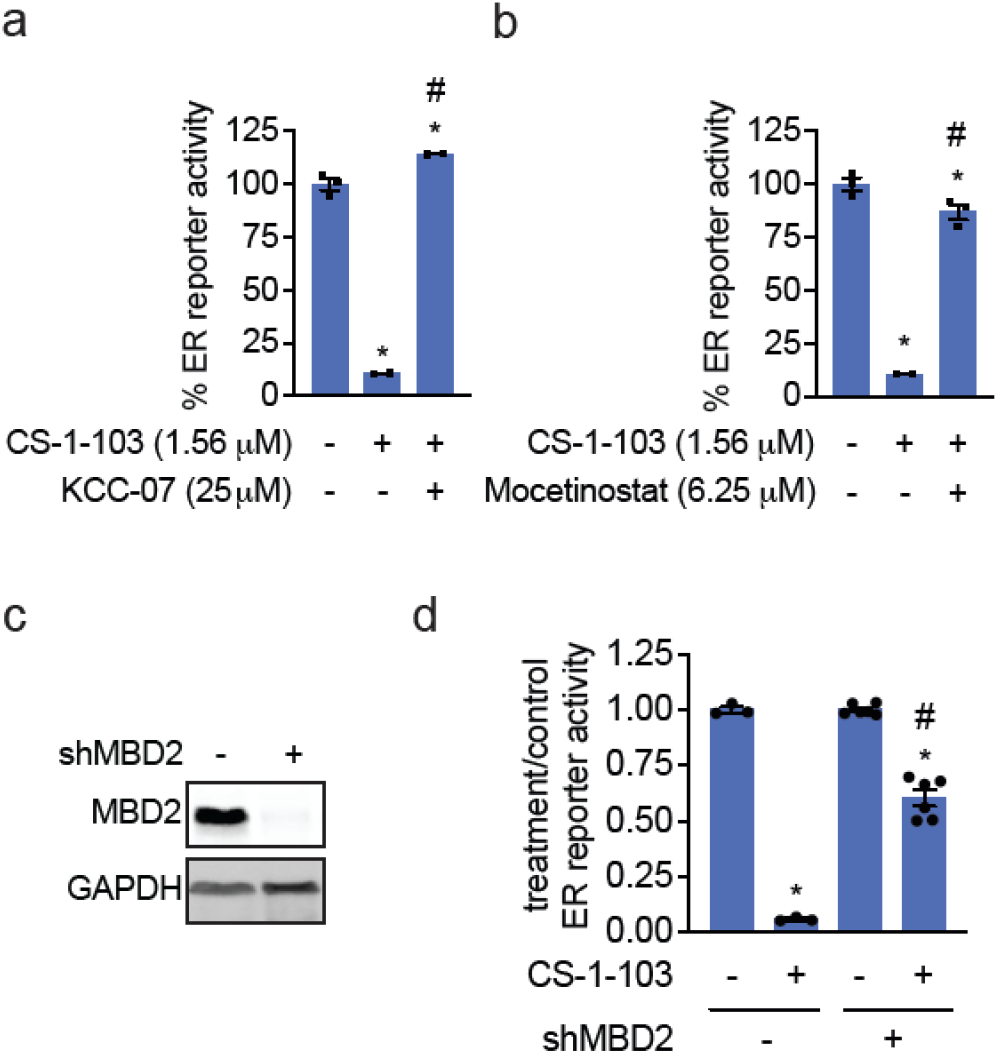
Confirming the mechanism of ER TRACER. **(a**,**b)** Attenuation of CS-1-103-mediated ER luciferase reporter inhibition in T47D cells with KCC-07 or HDAC inhibitor Mocetinostat. ER luciferase reporter T47D cells were co-treated with DMSO vehicle, KCC-07 **(a)**, or pre-treated with Mocetinostat **(b)** for 1h prior to treatment of cells with DMSO vehicle or CS-1-103 for 24 h, after which ER luciferase transcriptional activity was read out. **(c)** Stable short hairpin RNA (shRNA) knockdown of MBD2 in T47D cells assessed by SDS/PAGE and Western blotting alongside loading control GAPDH. **(d)** Attenuation of CS-1-103-mediated inhibition of T47D ER luciferase reporter activity upon MBD2 knockdown. T47D shControl versus shMBD2 cells were treated with DMSO vehicle or CS-1-103 (1.56 μM) for 24 h, after which ER luciferase transcriptional activity was read out. Data in **(a-d)** are from n=3-6 biologically independent replicates per group. Data in **(a**,**b**,**d)** show individual replicate values and average ± sem. Blot in **(c)** is representative. Significance is expressed as *p<0.05 compared to vehicle-treated control and #p<0.05 compared to CS-1-103-treated groups in **(a**,**b)** and CS-1-103-treated shControl cells in **(d)**.

**Figure 4.**
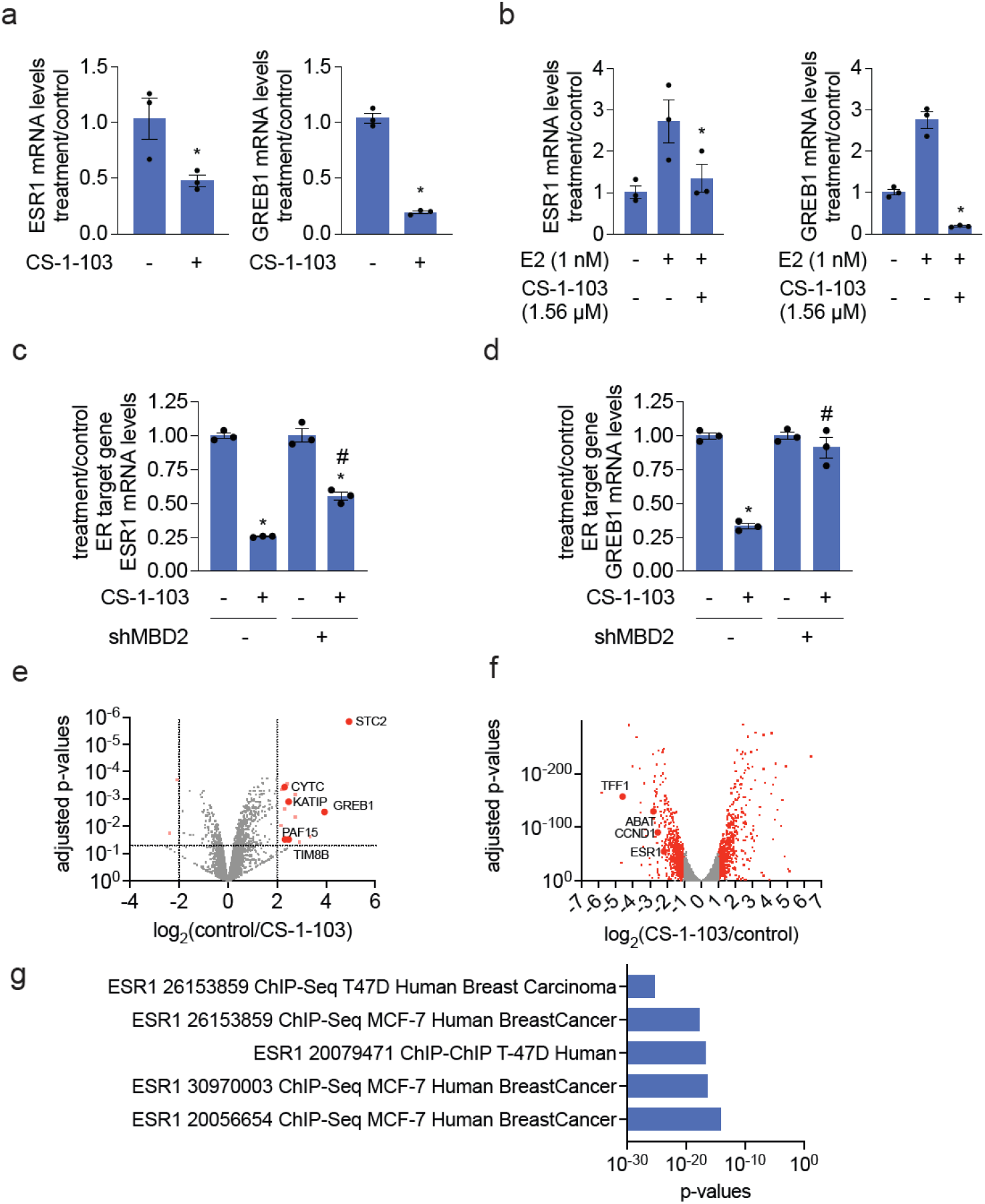
Characterization of ER TRACER. **(a)** qPCR of ER target genes with CS-1-103 treatment. T47D cells grown in normal serum were treated with DMSO vehicle or CS-1-103 (1.56 μM) for 24 h, after which ESR1 and GREB1 mRNA levels were assessed by qPCR. **(b)** qPCR of ER target genes with CS-1-103 treatment. T47D cells were grown under charcoal-stripped serum for 72 h, followed by co-treatment with E2 (1 nM) and DMSO vehicle or CS-1-103 (1.56 μM) for 24 h, and ESR1 and GREB1 mRNA levels were assessed by qPCR. **(c**,**d)** Attenuation of CS-1-103-mediated downregulation of ER target genes upon MBD2 knockdown. T47D shControl and shMBD2 cells were treated with DMSO vehicle or CS-1-103 (1.56 μM) for 24 h, and ESR1 **(c)** and GREB1 **(d)** mRNA levels were assessed by qPCR. **(e)** Proteomic analysis of T47D cells treated with DMSO vehicle or CS-1-103 (12.5 μM) for 24 h. **(f)** RNAseq transcriptomic data of T47D cells co-treated with 1 nM E2 and DMSO vehicle or CS-1-103 (1.56 μM) for 24 h. **(g)** Comparison of significantly down-regulated genes to the ChEA 2022 transcription factor targets. Data in **(a-g)** are n=3 biologically independent replicates per group. Data shown in **(a-d)** show individual replicate values and average ± sem. Significance is expressed as *p<0.05 compared to vehicle-treated controls and #p<0.05 compared to CS-1-103-treated shControl CS-1-103-treated groups.

To define the chromatin and transcriptional consequences of CS-1-103 treatment, we performed Assay for Transposase-Accessible Chromatin with sequencing (ATAC-seq^18^) in T47D cells and integrated these data with RNA-seq and ER-binding profiles. Motif analysis revealed that CS-1-103-responsive downregulated ATAC-seq peaks were strongly enriched for canonical ER response elements, as well as motifs for cooperating transcription factors including NFY, FOXA, and TEAD family members **(Figure 5a-5c; Table S3)**. Further analysis of the promoter regions that were associated with changes in either DNA-accessibility or RNA-expression revealed that these were not exclusive to ER-bound elements and resulted in changes inconsistent with repression. These could potentially reflect indirect downstream effects as well as differences in additional regulatory mechanisms modulating response to compound treatment (**Figure 5b)**. Thus, we next focused on only the subset of these promoters that had overlapping ER peaks and asked if the promoters with changes consistent with repression (ATAC-seq / and / or RNA-expression downregulated) had any sequence features discriminating them form the other categories (**Figure 5c)**. The ERβ motif (ERB) was enriched in the ATAC-seq and / or RNA-expression down categories and was depleted in the opposite categories. The opposite was observed for transcription factor binding motifs such as MITF and LHX6. These results suggest that CS-1-103 dependent transcriptional repression at ER-bound loci may be context dependent.

**Figure 5.**
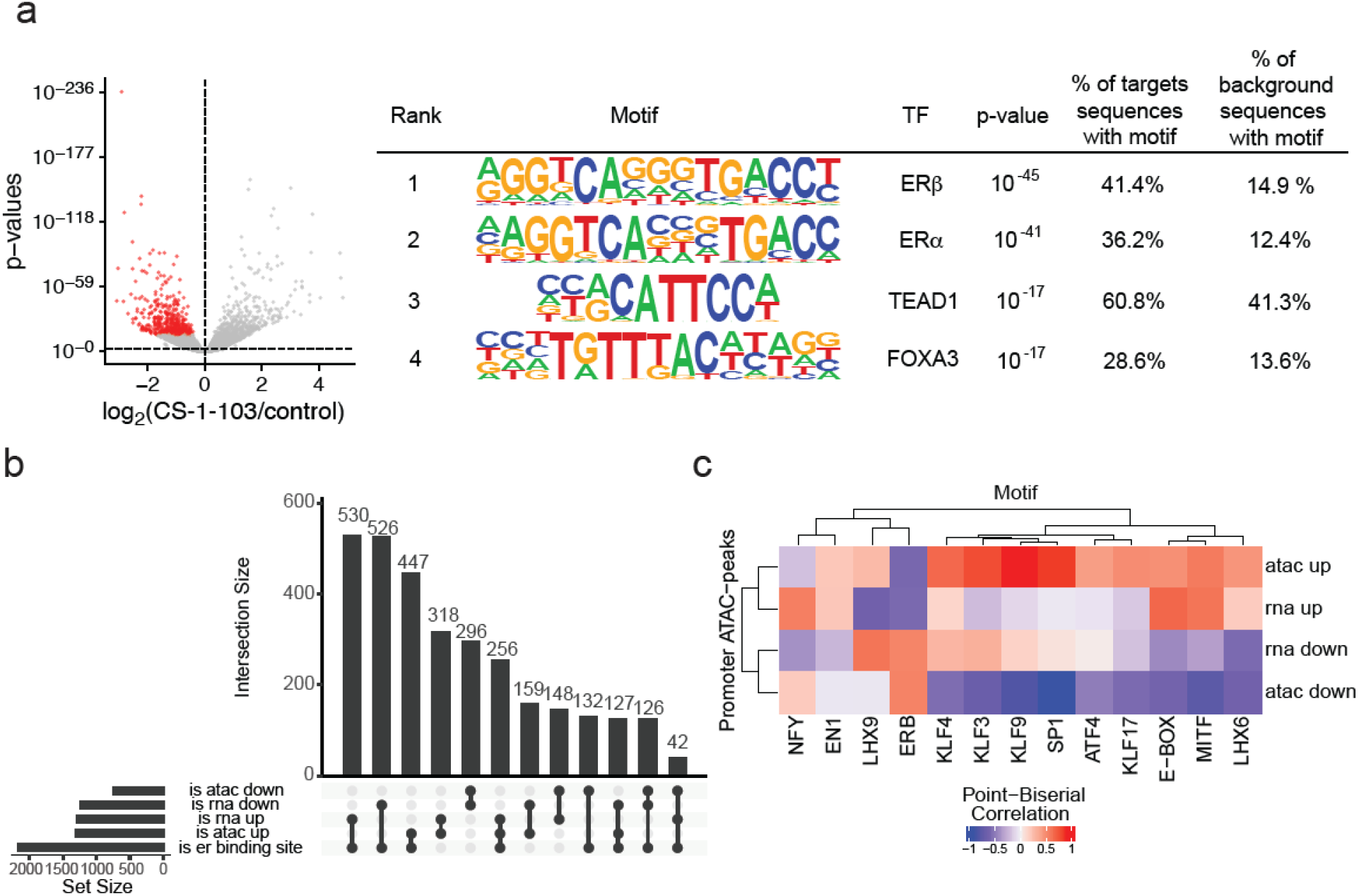
Integrative chromatin accessibility and transcriptional profiling of CS-1-103 TRACER activity in T47D cells. **(a)** Motif enrichment analysis of CS-1-103–responsive regions (500 downregulated ATAC-seq peaks with the highest p-values, see left panel), confirming top enrichment of canonical estrogen receptor binding motifs, along with TEAD and FOXA motifs (right panel). ATAC-seq data are from n=3 biologically independent replicates per group. **(b)** UpSet plot depicting the number of promoters with a change in accessibility (measured by ATAC-seq) or / and RNA expression with or without estrogen receptor peaks (measured by CUT&RUN). **(c)** Point-biserial correlation heatmap relating transcription factor motif enrichment to estrogen receptor target promoters from **(b)**. Motif enrichment is represented as the log_2_(odds ratio) from HOMER analysis. Peak categories are defined by significant changes in chromatin accessibility (ATAC up/down) and gene expression (RNA up/down). After CS-1-103 treatment the ERβ (ERB) motif is enriched in the promoters with downregulated ATAC-seq peaks and RNA-expression, while it is depleted in promoters with upregulated ATAC-seq and RNA-expression, respectively. ATAC-seq data can be found in **Table S3**.

### Development and Characterization of an AR TRACER

While ER was a promising proof-of-concept for our TRACER platform that may be able to tackle certain ER mutants, existing ER-targeting drugs, such as SERDs, Fulvestrant, and recently approved ER PROTACs, are likely to successfully tackle a large fraction of ER+ breast tumor patients. We sought to show TRACER proof-of-concept against a currently unmet oncology need with androgen-independent prostate cancers that are resistant to AR antagonists and current clinical AR PROTACs due to the expression of truncation variants, such as constitutively activated AR-V7, that are devoid of the ligand-binding domain, alongside full-length AR. While AR-V7 has been challenging to target directly due to its high intrinsic disorder, both AR and AR-V7 still bind to the same genetic loci. As such, epigenetic suppression of AR loci with a TRACER would also repress AR-V7 loci. To develop AR TRACERs, we linked a selective androgen receptor modulator (SARM) ligand RU59063 to our MBD2 recruiter using linkers of various lengths and composition to generate five AR/AR-V7 TRACERS—CS-1-162, CS-1-168, CS-1-169, CS-1-175, and CS-1-176 **(Figure S6a)**. Expectedly, RU59063 inhibited AR transcriptional activity by about only 40 % in 22Rv1 cells expressing both AR and AR-V7, since RU59063 binds to the ligand-binding domain of AR **(Figure 6a-6b)**. Upon testing the five AR and AR-V7 TRACERs in 22Rv1 AR transcriptional reporter assays, we found several compounds, including CS-1-162, CS-1-168, CS-1-175, and CS-1-176, that robustly inhibited total AR transcriptional activity by >90 % in a dose-responsive manner with EC_50_ values of 1.3, 4.7, 1.3, and 2.5 μM, respectively, while CS-1-169 only showed modest inhibition comparable to RU59063 **(Figure S6b)**. However, CS-1-162, CS-1-168, and CS-1-176 showed substantial impairments in cell viability, although the cytotoxic effects showed less potency than the transcriptional inhibitory effects **(Figure S6b)**. In contrast, CS-1-175 showed >90 % inhibition of total AR luciferase reporter transcriptional activity in 22Rv1 cells with minimal cytostatic effects **(Figure 6d-6e; Figure S6b)**. In addition to the reporter system, CS-1-175 also significantly downregulated endogenous AR target genes **(Figure 6f)**. This inhibition of AR transcriptional activity was attenuated by pre-incubation with an HDAC inhibitor, and the downregulation of AR target genes can be completely attenuated upon MBD2 knockdown, indicating on-target activity **(Figure 6g-6j)**. Transcriptomic profiling of CS-1-175 by RNAseq showed significant downregulation of many AR target genes **(Figure 7a; Figure S7; Table S4)**. Analysing effects upon genes that have been previously shown to be regulated by AR-V7^19^, CS-1-175 universally and significantly affected almost every AR-V7 target gene within this dataset **(Figure 7b)**. Pathway enrichment analysis showed a broader transcriptional signature than our ER TRACER, that included significant downregulation of androgen response genes, but also included MYC target genes, fatty acid and cholesterol metabolism, and DNA repair genes **(Figure 7c)**. Overall, we demonstrated that we could achieve targeted transcriptional repression against ER and AR/AR-V7 by developing TRACER heterobifunctional compounds that link an MBD2 recruiter to ER and AR ligand-binding domain modulators to durably suppress ER and AR transcriptional activity.

**Figure 6.**
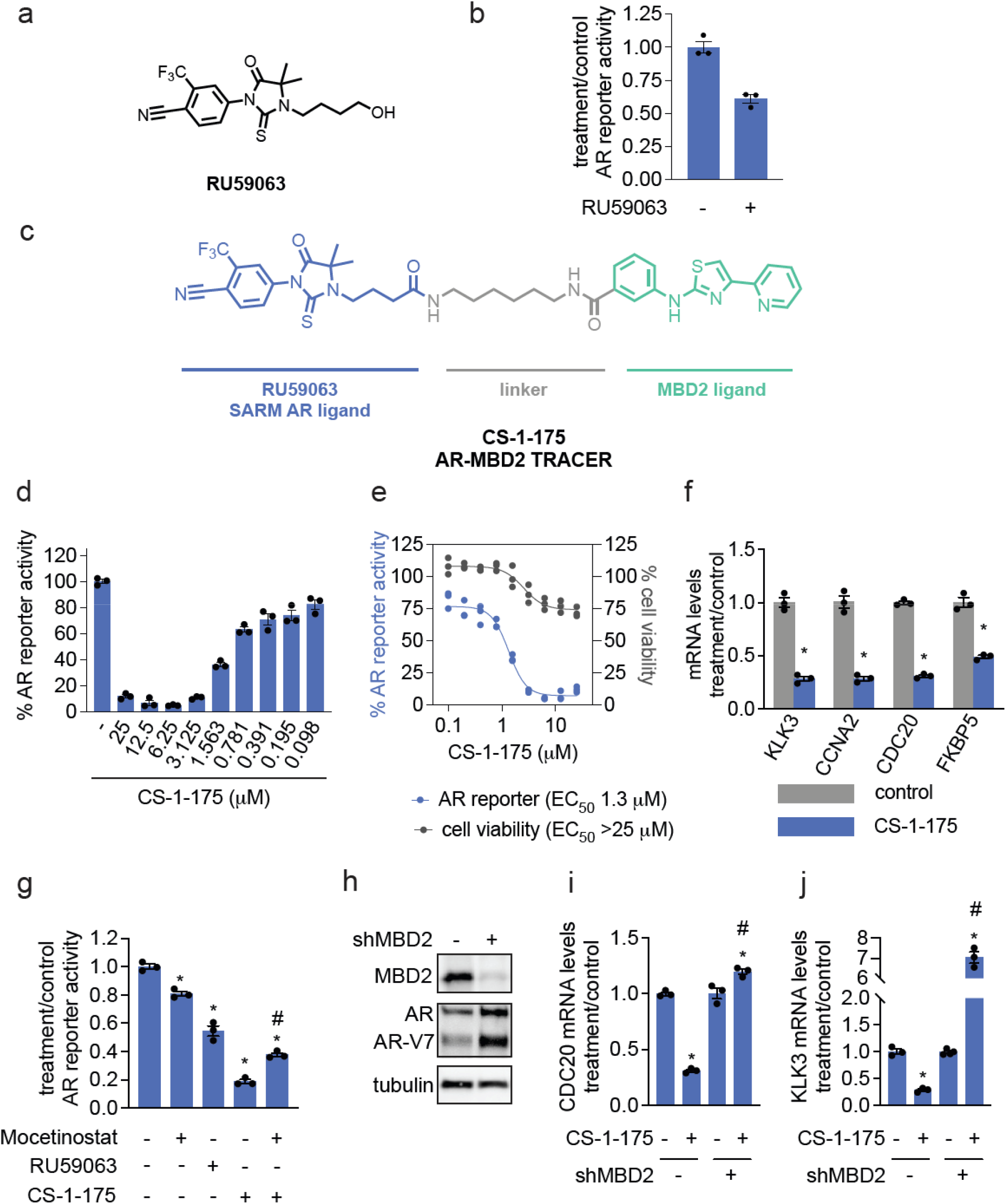
AR TRACER. **(a)** Structure of AR-targeting SARM RU59063. **(b)** AR luciferase reporter activity from RU59063 treatment in 22Rv1 androgen-independent prostate cancer cells. 22Rv1 cells were treated with DMSO vehicle or RU59063 (6.25 μM) for 24 h, after which AR transcriptional reporter activity was assessed. **(c)** Structure of AR TRACER CS-1-175. **(d**,**e)** Dose-responsive inhibition of AR luciferase reporter activity with CS-1-175 treatment in 22Rv1 cells compared to DMSO vehicle-treated controls **(d)**, with an EC_50_ of 1300 nM from 24 h treatment **(e)**. Also shown is cell viability assessed by Cell TiterGlo **(e). (f)** Downregulation of AR target genes with CS-1-175 treatment (6.25 μM) in 22Rv1 cells for 24 h compared to DMSO vehicle-treated controls. **(g)** CS-1-175-mediated inhibition of AR luciferase reporter activity in 22Rv1 cells is significantly attenuated upon pre-treatment with HDAC inhibitor Mocetinostat. 22Rv1 cells were pre-treated with DMSO, RU59063 (6.25 μM), or Mocetinostat (0.38 μM) for 1 h prior to treatment of cells with DMSO vehicle or CS-1-175 for 24 h. AR transcriptional reporter activity was subsequently assessed. **(h)** MBD2 knockdown in 22Rv1 cells. MBD2, AR, AR-V7, and loading control tubulin levels in Control versus shMBD2 cells. **(i**,**j)** qPCR of AR target genes. Control and shMBD2 22Rv1 cells were treated with DMSO vehicle or CS-1-175 (6.25 μM) for 24 h, and CDC20 **(i)** and KLK3 mRNA **(j)** levels were assessed by qPCR. Data in **(b, d-j)** are from n=3 biologically independent replicates per group. Blot shown in **(h)** is representative. Data shown in **(b, d-g, i**,**j)** are individual replicate values and average ± sem. Significance is expressed as *p<0.05 compared to vehicle-treated controls in **(f**,**g**,**i**,**j)** and #p<0.05 compared to CS-1-175-treated CS-1-175-treated or Control CS-1-175-treated groups in **(g**,**i**,**j)**.

**Figure 7.**
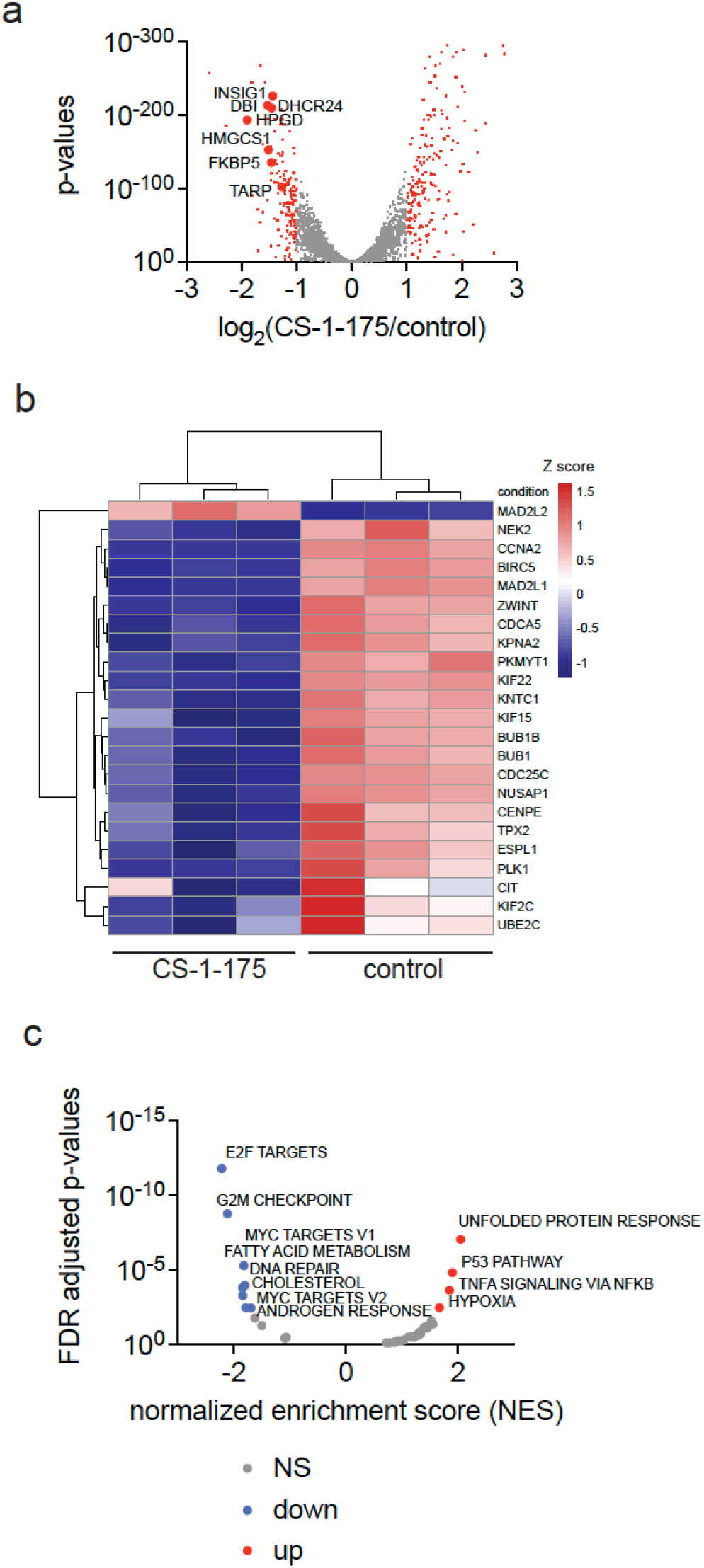
Transcriptomic analysis of AR TRACER CS-1-175. **(a**,**b**,**c)** 22Rv1 cells were treated with DMSO vehicle or CS-1-175 (12.5 μM) for 24 h, after which mRNA was extracted and subjected to RNAseq. Significantly altered genes were analyzed **(a)**, Heatmap analysis of previously identified AR-V7 target genes **(b)**, and hallmark gene set enrichment analysis was performed to identify compound-mediated alterations in biological pathways **(b)**. Data are from n=3 biologically independent replicates per group and data can be found in **Table S4**.

### Exploiting Other Epigenetic Targets for TRACER Approaches

We next sought to determine whether we could use existing inhibitors against other epigenetic repression complexes to expand the TRACER platform. We first assessed whether the Polycomb Repressor Complex (PRC) could be exploited using the EED targeting PRC2 inhibitor EED226.^20^ We linked an EED226 to estrone to generate CS-1-129 and CS-1-132 and were pleased to show that these molecules also suppressed ER transcriptional luciferase reporter activity in T47D cells, while EED226 treatment itself did not show any effect **(Figure S8a-8c)**. We also showed ER transcriptional repression with ER TRACER, CS-1-123 that is based on EZH2-methyltransferase inhibitor GSK126 **(Figure S9a-S9b)**.^21^ Even though these molecules were not as potent as our MBD2-based TRACERs, we demonstrated that the TRACER approach could also be expanded to encompass additional epigenetic repression machinery.

## Discussion

The TRACER platform described here introduces a fundamentally new paradigm for pharmacologically targeting transcription factors—an enduring frontier in drug discovery. By co-opting endogenous transcriptional corepressor complexes and tethering them to disease-relevant transcription factor binding sites, TRACERs overcome many of the liabilities that have hindered traditional small molecules, degraders, and even emerging modalities in this space. The demonstration that locus-specific repression can be achieved with small molecules in cancer cells establishes proof-of-concept for a modality that is modular, generalizable, and uniquely suited to tackle proteins long considered undruggable.

Several aspects of this study underscore the strength of this approach. First, TRACERs achieve durable and selective suppression of transcriptional programs, in contrast to the transient and often incomplete inhibition observed with antagonists or degraders of transcription factors with rapid turnover. Second, our results validate the on-target activity of TRACERs through multiple orthogonal approaches, including rescue with MBD2 knockdown, attenuation by HDAC inhibitors, and transcriptomic profiling demonstrating selective repression of bona fide target genes. Third, TRACERs can address clinically important resistance mechanisms. For example, while androgen receptor truncation variants such as AR-V7 evade ligand-directed antagonists and degraders, AR-TRACERs effectively silence both AR and AR-V7 transcriptional activity at their shared loci. Finally, the adaptability of this platform is underscored by our extension to alternative repressor complexes such as PRC2, highlighting its potential modularity.

At the same time, several caveats merit consideration. Our proof-of-concept studies have thus far been confined to in vitro models, and the pharmacokinetic, pharmacodynamic, and safety properties of TRACERs in vivo remain to be established. Because TRACERs rely on hijacking endogenous corepressor complexes, differential expression or activity of these complexes across tissues may affect efficacy and selectivity.

Furthermore, while our transcriptomic analyses suggest locus-selective repression, comprehensive genome-wide profiling will be needed to fully define off-target effects and to ensure that broad epigenetic remodeling does not undermine the therapeutic window. The chemical optimization of recruiter ligands, linker architectures, and transcription factor–binding ligands will also be critical to maximize potency, selectivity, and drug-like properties.

Looking ahead, TRACERs open several exciting avenues. The modular nature of this platform suggests it can be extended to a wide range of transcription factors beyond ER and AR, many of which remain untreatable yet central to cancer and other diseases. Systematic exploration of alternative repressor complexes—such as NCoR/SMRT, SIN3A, or additional Polycomb family members—could expand the toolkit for tuning repression strength, durability, and context-specificity. From a translational standpoint, TRACERs may be particularly impactful in malignancies driven by transcription factors for which no targeted therapies exist, such as MYC, CTNNB1, and YAP/TAZ. Beyond oncology, the ability to selectively reprogram gene expression programs raises the possibility of using TRACERs in inflammatory disorders, autoimmune disease, and even rare genetic conditions where pathological gene activation underlies disease.

In summary, TRACERs establish a new modality for silencing transcriptional drivers of disease by coupling the precision of small molecules with the power of epigenetic reprogramming. By addressing long-standing challenges in drugging transcription factors, this work lays the foundation for a versatile and expandable therapeutic strategy with broad implications across human disease.

## Acknowledgment

We thank the members of the Nomura Research Group and Novartis BioMedical Research for critically reading the manuscript. This work was also supported by Novartis Biomedical Research, the National Science Foundation Molecular Foundations for Biotechnology (MFB) grant (2127788), the UC Berkeley Molecular Therapeutics Initiative (MTI), the Mark Foundation for Cancer Research ASPIRE Award, the National Institutes of Health (R35CA263814, R01CA240981), and the Bakar Award. CS is supported by a Walter-Benjamin Postdoctoral Fellowship by the German Research Foundation (DFG). We also thank Hasan, Lund, and the UC Berkeley NMR facility in the College of Chemistry (CoC-NMR) for spectroscopic assistance. Instruments in the College of Chemistry NMR facility are partly supported by NIH S10OD024998.

## Author Contributions

CS, DKN conceived of and co-directed the project. CS and XC performed experiments for the paper. FW, NP, and BF provided critical insights and analysed data. DD, JM, MS, CJ provided critical insights into project experiments. CS, DKN wrote the paper.

