## Supporting Information for "Targeted Transcriptional Repression by Induced Proximity"

[1] Departments of Chemistry and Molecular and Cell Biology. University of California, Berkeley, Berkeley, CA 94720 USA

[2] Innovative Genomics Institute, Berkeley, CA 94720 USA

[3] Molecular Therapeutics Initiative, Berkeley, CA 94720 USA

[4] Novartis-Berkeley Translational Chemical Biology Institute, Berkeley, CA 94720 USA

[5] Novartis BioMedical Research, Emeryville, CA USA; Cambridge, MA USA; Basel, Switzerland

**Supporting Table Legends**

**Table S1. Proteomic profiling of CS-1-103.** Proteomic analysis of T47D cells treated with DMSO vehicle or CS-1-103 (12.5 μM) for 24 h. Data are from n=3 biologically independent replicates per group.

**Table S2. RNA sequencing of CS-1-103.** RNAseq transcriptomic data of T47D cells co-treated with 1 nM E2 and DMSO vehicle or CS-1-103 (1.56 μM) for 24 h. Data are from n=3 biologically independent replicates per group.

**Table S3. ATTACseq of CS-1-103.** ATTAC-seq data from T47D cells treated with DMSO vehicle or CS-1-103 treatment (1.56 μM) with 1 nM E2 over 24 h treatment time.

**Table S4. RNA sequencing of CS-1-175.** RNAseq transcriptomic data of 22Rv1 treated with DMSO vehicle or CS-1-175 (12.5 μM) for 24 h, after which mRNA was extracted and subjected to RNAseq. Data are from n=3 biologically independent replicates per group.

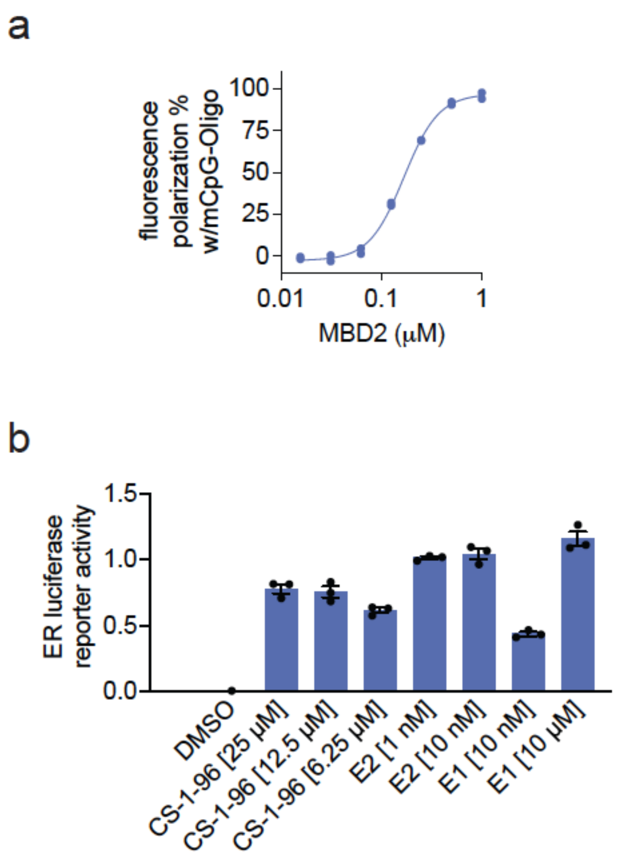


**Figure S1. Establishing MBD2-mCpG-Oligo fluorescence polarization assay. (a)** Pure MBD2 protein was incubated with a fluorescent methylated CpG oligonucleotide (20 nM) binding was determined by fluorescence polarization. Shown are data from n=2 biologically independent replicates per group. **(b)** ER luciferase transcriptional reporter assay in T47D cells grown in charcoal-stripped media where ER agonists were added back and luciferase reporter activity was assessed. Shown in **(a,b)** are individual replicate values from n=2-3 biologically independent replicates per group.

a

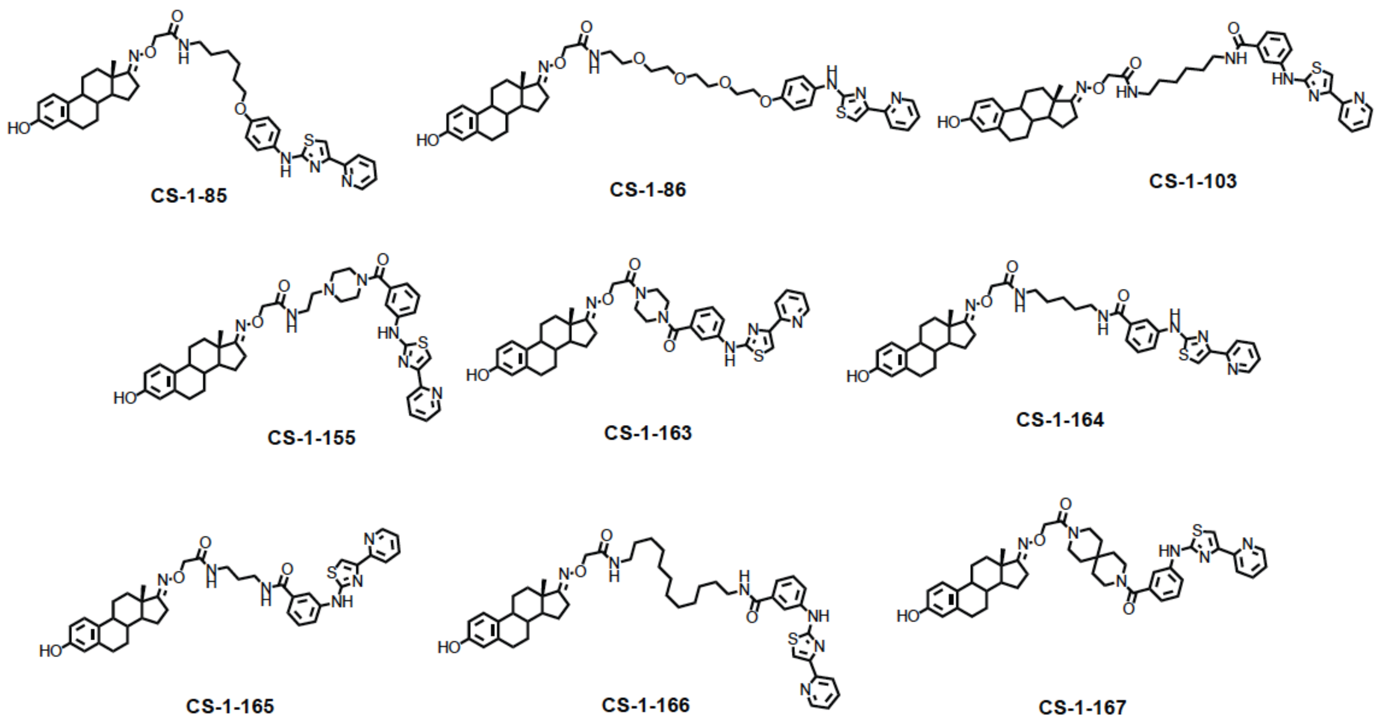


b

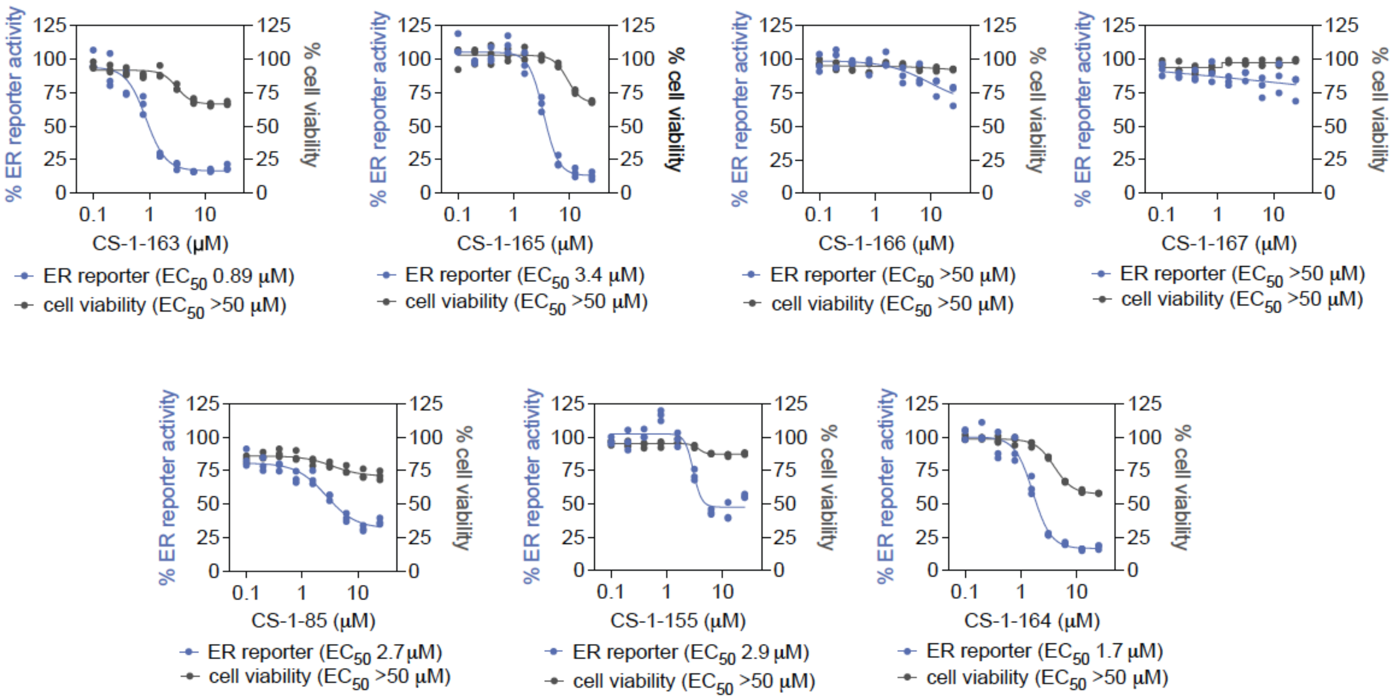


**Figure S2. Characterization of ER TRACERs. (a)** Structure of ER TRACER CS-1-86. **(b)** Dose-responsive inhibition of ER luciferase transcriptional reporter activity and cell viability (assessed by Cell TiterGlo) in T47D cells upon treatment with DMSO vehicle or TRACER for 24 h, showing EC50 values. Shown in **(b)** are individual replicate values from n=3 biologically independent replicates per group.

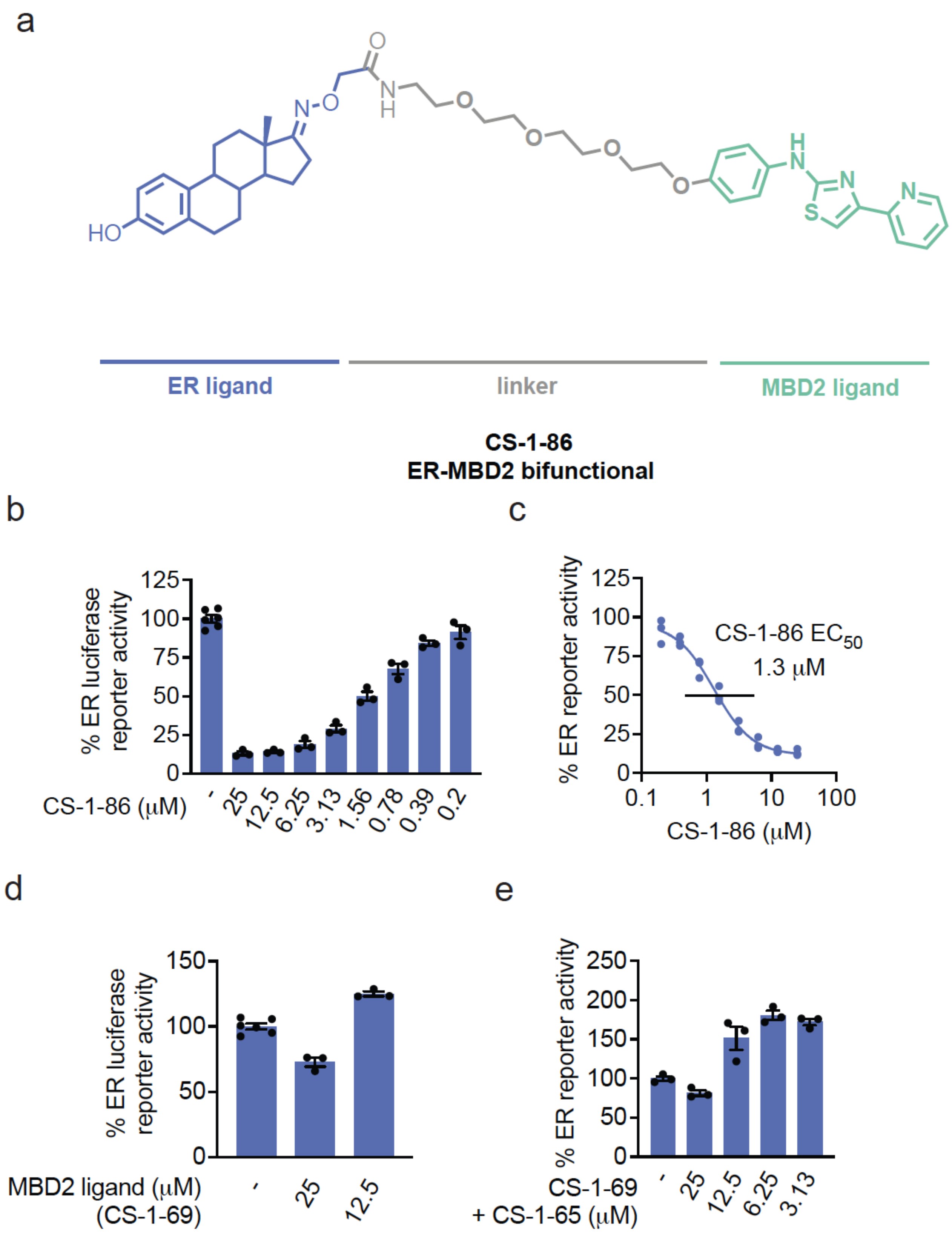


**Figure S3. Characterization of CS-1-86. (a)** Structure of ER TRACER CS-1-86. **(b,c)** Dose-responsive inhibition of ER luciferase transcriptional reporter activity in T47D cells upon treatment with DMSO vehicle or TRACER CS-1-86 for 24 h, showing an $EC_{50}$ of 1300 nM. **(d,e)** T47D ER luciferase reporter activity after treatment with DMSO vehicle or CS-1-69 or CS-1-69 and CS-1-65 treatment for 24 h. Data in **(b-e)** are from n=3 biologically independent replicates per group. Shown in **(b-e)** are individual replicate values and average ± sem for **(b,d,e)**.

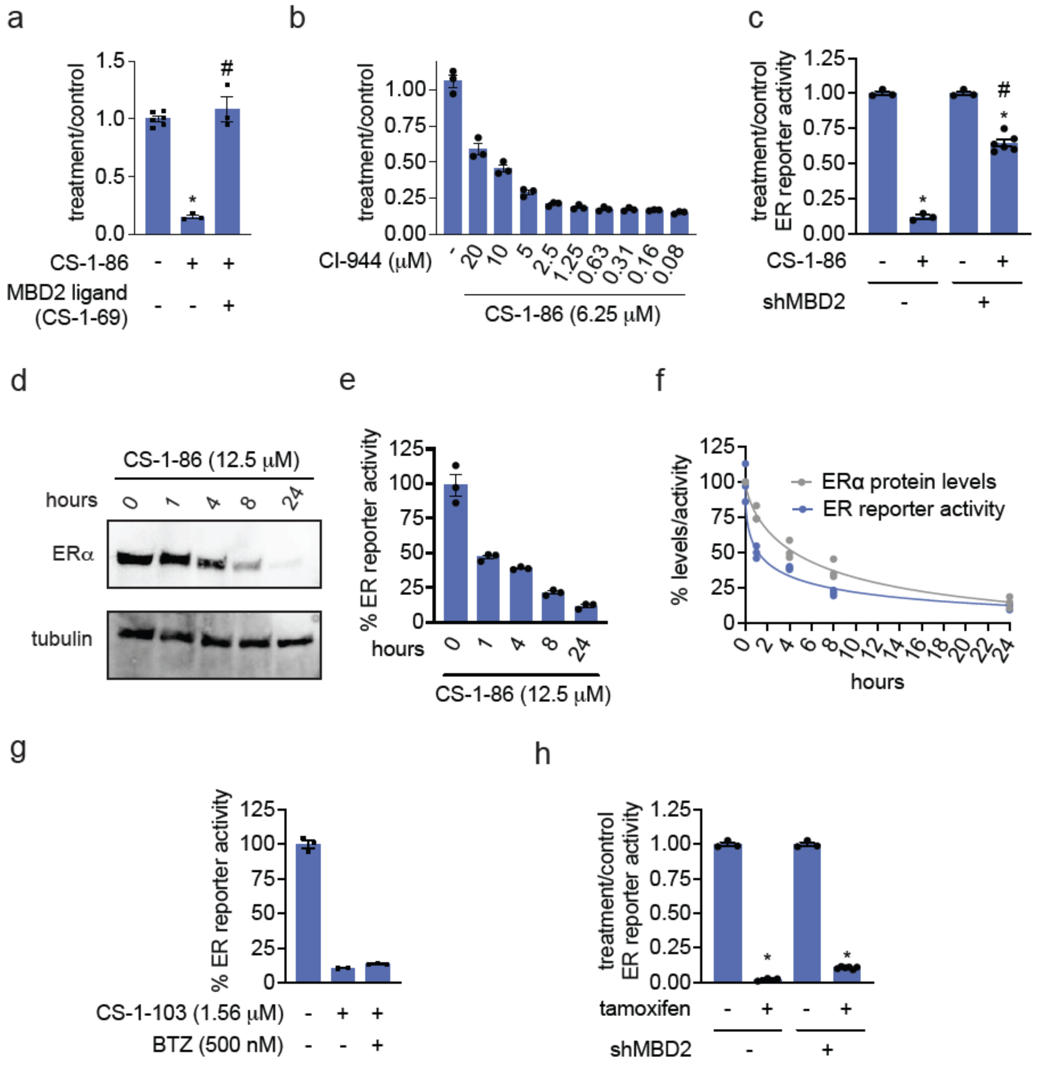


**Figure S4. Further characterization of CS-1-86. (a,b)** Attenuation of CS-1-86-mediated ER luciferase reporter inhibition in T47D cells with CS-1-69 or HDAC inhibitor CI-944. ER luciferase reporter T47D cells were co-treated with DMSO vehicle, CS-1-69 (50 μM) **(a)**, or pre-treated with CI-944 **(b)** for 1h prior to treatment of cells with DMSO vehicle or CS-1-86 (12.5 μM or 6.25, respectively) for 24 h, after which ER luciferase transcriptional activity was read out. **(c)** Attenuation of CS-1-86-mediated inhibition of T47D ER luciferase reporter activity upon MBD2 knockdown. T47D shControl versus shMBD2 cells were treated with DMSO vehicle or CS-1-86 (12.5 μM) for 24 h, after which ER luciferase transcriptional activity was read out. **(d),e** ER protein levels from CS-1-86 treatment. T47D cells were treated with CS-1-86 (12.5 μM), and ER and loading control tubulin levels were assessed by SDS/PAGE and Western blotting **(d)** and quantified in **(e). (f)** ER transcriptional reporter activity and ER protein levels from CS-1-86 treatment in a time-course study. T47D cells were treated with CS-1-86 (12.5 μM) for the designated time-points, and ER luciferase reporter activity and ER protein levels were assessed and plotted. **(g)** CS-1-103-mediated ER transcriptional reporter activity is not attenuated by proteasome inhibitor pre-treatment. ER luciferase reporter T47D cells were pre-treated with DMSO vehicle or bortezomib (500 nM) for 1 h prior to treatment of cells with DMSO vehicle or CS-1-103 (1.56 μM) for 24 h, after which ER luciferase reporter activity was read out. **(h)** Tamoxifen-mediated inhibition of ER transcriptional reporter activity is not MBD2-dependent. T47D Control and shMBD2 cells were treated with DMSO vehicle or tamoxifen (10 μM) for 24 h, after which ER luciferase reporter activity was read out. Data in **(a-d)** are from n=3-6 biologically independent replicates per group. Data in **(a-h)** are from n=3 biologically independent replicates per group. Blot in **(d)** is representative. Bar graphs and plots in **(a-h)** show individual replicate values and bar graphs in **(a-e, c,g,h)** show average ± sem. Significance expressed as *p<0.05 compared to vehicle-treated controls in **(a,c,h)** and #p<0.05 compared to CS-1-86 treatment alone in **(a)** or CS-1-86 treatment in Control cells in **(c).**

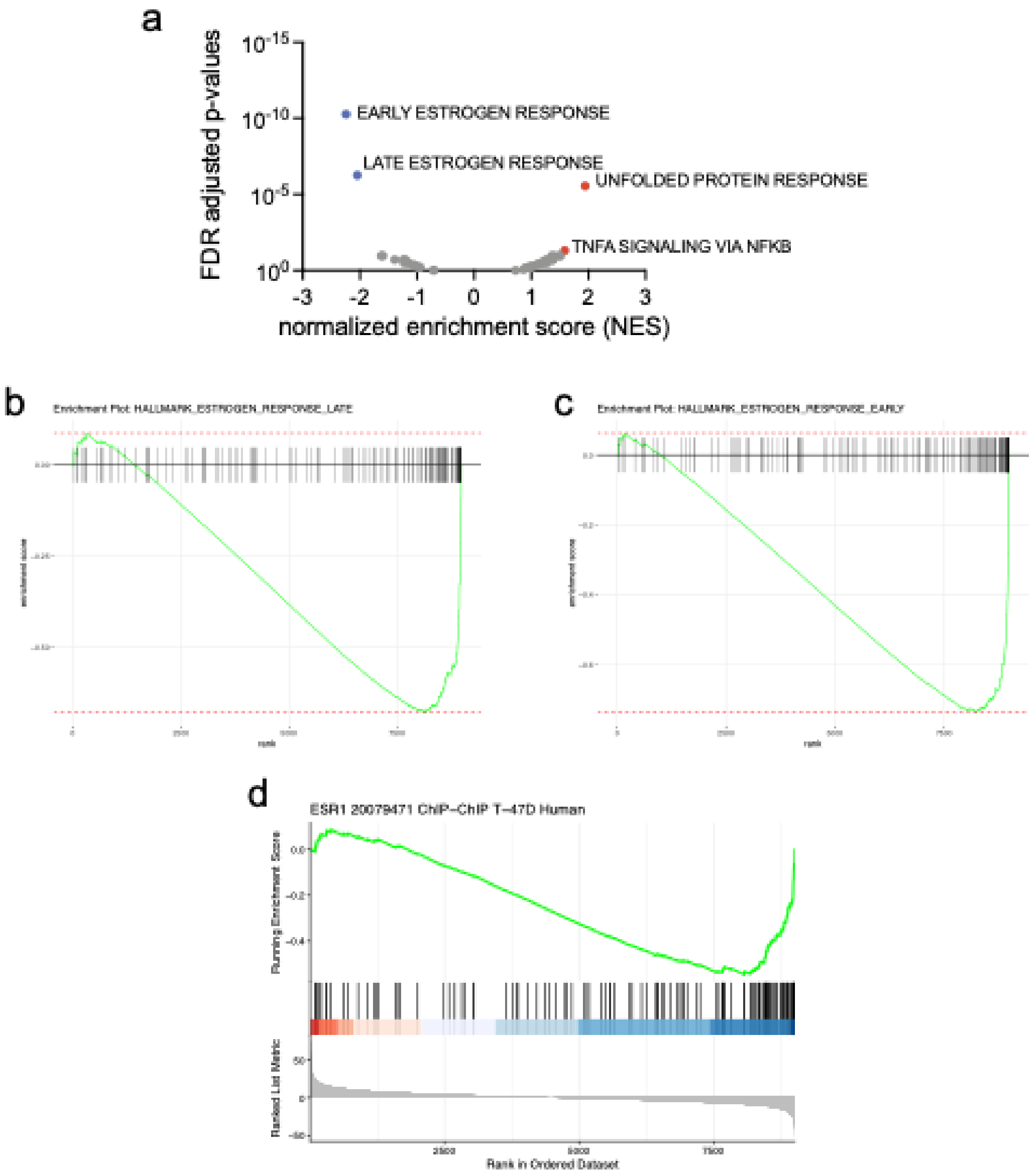


**Figure S5. Gene enrichment analysis of ER TRACER RNAseq data. (a)** fGSEA hallmark gene enrichment analysis of ER TRACER. **(b)** Enrichment plot of preset hallmark gene sets of cells treated with ER-TRACER compared to control. Top ranked downregulated gene sets are shown. **(c)** Enrichment plot of ER-TRACER regulated genes on a representative ChEA 2022 ESR1 dataset.

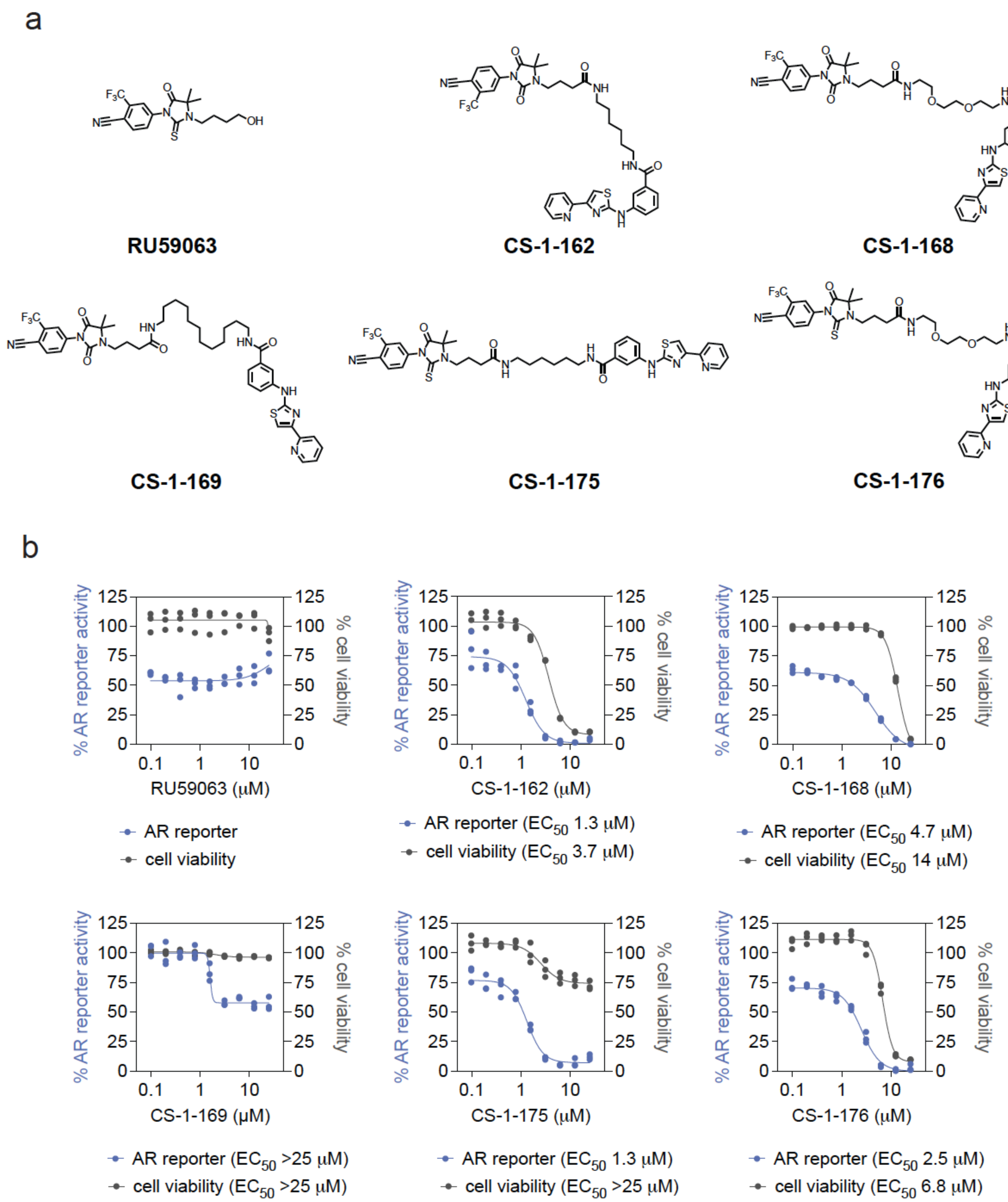


**Figure S6. Characterization of AR TRACERs. (a)** Structure of SARM RU59063 and MBD2-based AR TRACERs with varying linkers. **(b)** AR luciferase transcriptional reporter activity and cell viability in 22Rv1 cells. AR luciferase reporter 22Rv1 cells were treated with RU59063, CS-1-162, CS-1-168, CS-1-169, CS-1-175, or CS-1-176 for 24 h after which AR luciferase reporter activity was read out and cell viability was also assessed by Cell TiterGlo. Shown below each plot are $EC_{50}$ values for both AR transcriptional reporter activity and cell viability. Data in **(b)** are from n=3 biologically independent replicates per group and shown are individual replicate values.

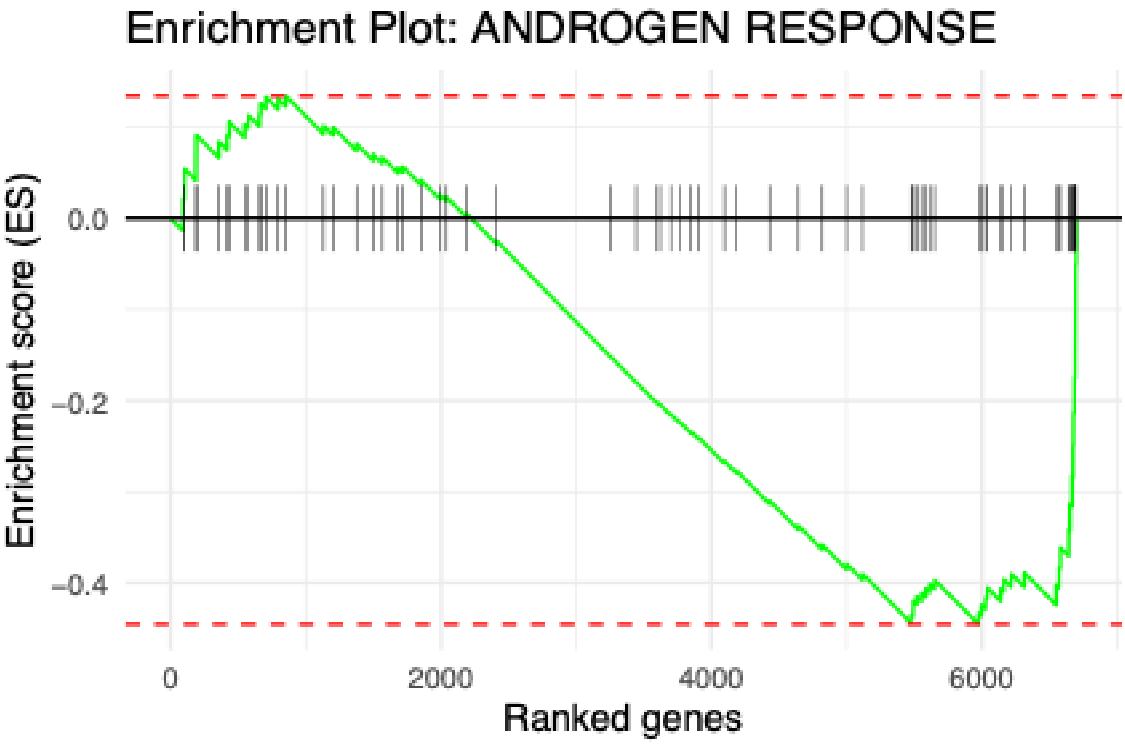


**Figure S7. Transcriptomic analysis of AR TRACER CS-1-175.** 22Rv1 cells were treated with DMSO vehicle or CS-1-175 (12.5 μM) for 24 h, after which mRNA was extracted and subjected to RNAseq. Shown is an enrichment plot of hallmark androgen response genes from these data.

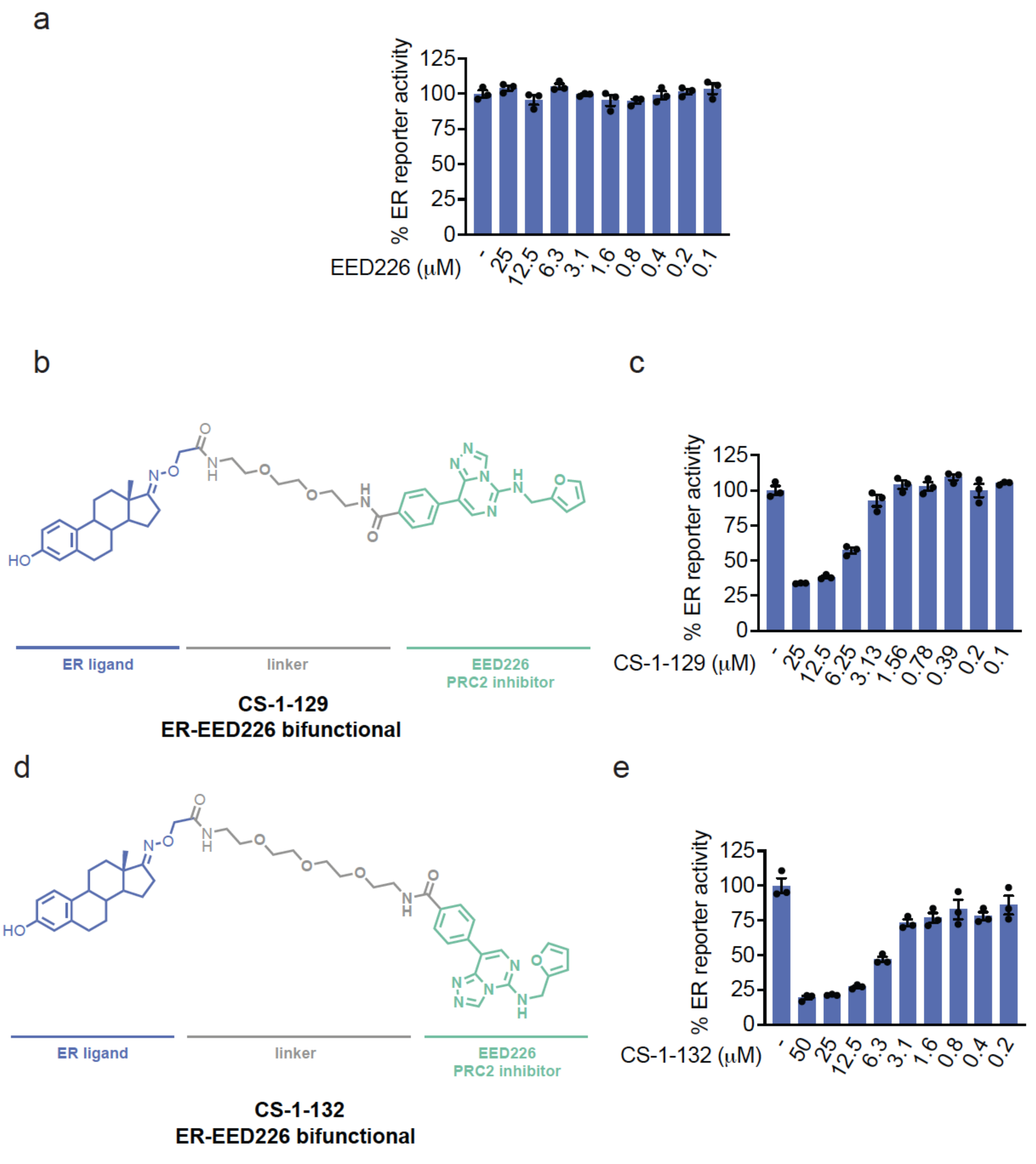


**Figure S8. TRACERs using additional handles for epigenetic corepressor complexes. (a-e)** Effects of EED226 and EED226-based ER TRACERs CS-1-129 and CS-1-132 on ER transcriptional reporter activity. ER luciferase reporter T47D cells were treated with DMSO vehicle, EED226, CS-1-129, or CS-1-132 for 24 h, after which ER luciferase reporter activity was read out. Structures of CS-1-129 and CS-1-132 are shown in **(b,d)**. Data in **(a,c,e)** are from n=3 biologically independent replicates per group. Data show individual replicate values and average ± sem.

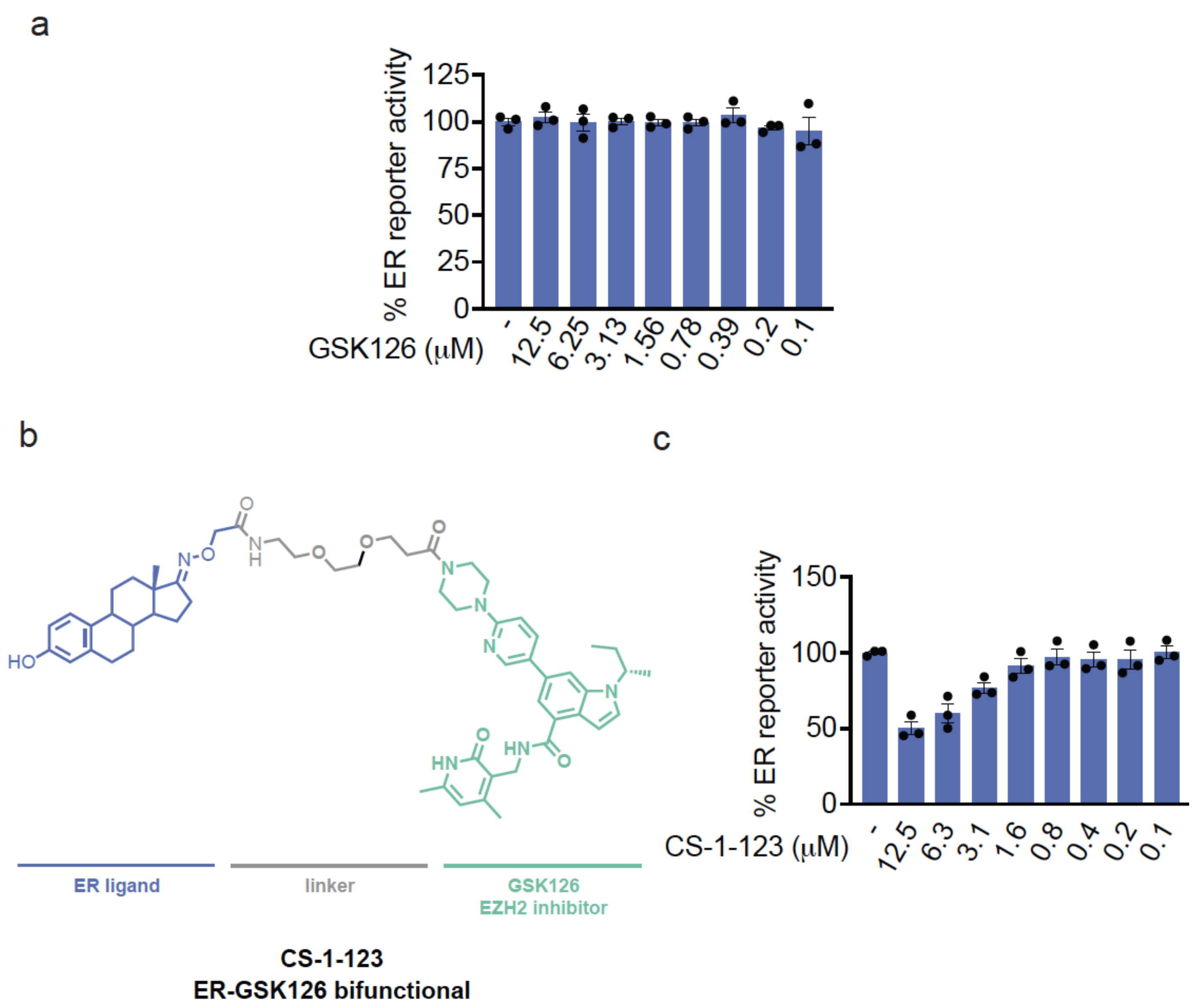


**Figure S9. TRACERs using additional handles for epigenetic corepressor complexes. (a-c)** Effects of GSK126 and GSK126-based ER TRACER CS-1-123 on ER transcriptional reporter activity. ER luciferase reporter T47D cells were treated with DMSO vehicle, GSK126, or CS-1-123 for 24 h, after which ER luciferase reporter activity was read out. The structure of CS-1-123 is shown in **(b)**. Data in **(a,c)** are from n=3 biologically independent replicates per group. Data show individual replicate values and average ± sem.

### Supporting Methods

#### Fluorescence Polarization Assay

Fluorescein-labeled mCpG-DNA-oligos (20 nM final concentration), recombinant MBD2, and ligands (50 mM stock solution in DMSO) were diluted in buffer (DPBS pH 7.4, 1 mM $MgCl_2$). The final DMSO concentration used in the assay was always <10 %. A dilution series of MBD2 was prepared in black 384-microwell plates (Corning) in a final sample volume of 20 µL in triplicate. mCpG-DNA-oligos (20 nM) and recombinant MBD2 (150 nM) were incubated followed by the addition of the indicated compound concentration and fluorescence anisotropy was measured after 30 min incubation at room temperature, using a Tecan plate reader (filter set $\lambda_{ex}$: 485 ± 20 nm, $\lambda_{em}$: 535 ± 25 nm).

#### Cell Culture

T47D wild-type (obtained from UC Berkeley Cell Culture Facility) and Estrogen Receptor Luciferase Reporter T47D Stable Cell Line (obtained from Signosis, SL-0002) was cultured in RPMI-1640 medium containing 10% (v/v) fetal bovine serum (FBS) and 1x glutamine. For hormone competition experiments, cells were grown in phenol-red-free RPMI supplemented with 10% charcoal-treated FBS for at least 72 h prior to treatment.
22Rv1wild-type and Androgen Receptor Luciferase Reporter 22Rv1 cells (both from UC Berkeley Cell Culture Facility) were cultured in RPMI-1640 medium containing 10% (v/v) fetal bovine serum (FBS) and 1× GlutaMAX.
HEK293T (UC Berkeley Cell Culture Facility) were cultured in DMEM containing 10% (v/v) fetal bovine serum (FBS).
All cell lines were maintained at 37°C and 5% $CO_2$.

#### Luciferase Reporter Assay

Cells were seeded in white 96-well plates at 20,000 cells/well in 100 µL complete medium and allowed to adhere overnight. 10x stocks of the respective small molecule in complete medium were added to the wells to a final DMSO concentration of 1%. Cells were incubated for the indicated time before luciferase signal was measured using the Bright-Glo® Luciferase Assay System (Promega, E2620). In parallel, cell viability was assessed using CellTiter-Glo® 2.0 assay (Promega, G9242). Values for luciferase activity and viability were normalized to the DMSO and blank controls in Prism 10 (GraphPad), and curves were fit using a nonlinear regression. Additionally, the luciferase signal was normalized to viability.

#### MBD2 Lentiviral Knockdown Studies

Lentiviral stable MBD2 knockdown studies in T47D and 22rv1 cells were performed, as previously described. 2 µg of the following plasmids, shRNA construct of MBD2 (carrying a puromycin resistance gene, obtained from Vectorbuilder), psPAX2 (carrying GAG, REV, and pol genes), and pMD2G (carrying the VSVG pseudotyping gene), were dissolved in 1.2 mL of Gibco Opti-MEM Reduced Serum Medium (catalog no. 31985–062). In parallel, lipofectamine 2000 (Invitrogen, 11668019) was diluted in 1.2 mL of Gibco Opti-MEM Reduced Serum Medium. After 5 min, after which the two tubes were mixed and incubated for another 30 min at room temperature and subsequently the mixture was added to HEK293T cells at 30-40% confluence (cultured in DMEM + 10% heat inactivated FBS). The following day the medium was replaced to fresh DMEM + 10% heat inactivated FBS, and the cells were incubated for 48-72 h. On the day of infection, the virus containing medium was collected from the HEK293T cells filtered through a sterile 0.45 µm filter and combined with an equal volume of the target cell line media containing polybrene (Sigma-Aldrich, TR-1003-G). The lentiviral mixture was then added to the target cells and incubated for 24 h. Following a medium exchange to target cell line medium and 24 h incubation, infected cells were selected using puromycin (2.5 µg/ml) for 96 h followed by recovery in complete medium.

**Western Blotting**

Cells were lysed using RIPA buffer supplemented with Benzonase® Nuclease (1:1000; Millipore, 70746) for 15 min at 37°C and protein concentration was determined using Pierce™ BCA Protein Assay Kit (Thermo Fisher) according to the manufacturers protocol. The protein amount was normalized between the samples and proteins were separated on precast 4–20% Criterion TGX gels (Bio-Rad) followed by a transfer to a nitrocellulose membrane using the Trans-Blot Turbo transfer system (Bio-Rad). After the transfer, the membrane was blocked with Tris-buffered saline containing Tween 20 (TBST) containing 5% bovine serum albumin (BSA) for 1 h at RT. After blocking, target proteins were probed with primary antibodies in TBST with 5% BSA (primary antibody dilutions were performed according to the manufacturer). Incubation with primary antibodies was performed overnight at 4°C. After washing the membrane trice with TBST (5 min each) the membrane was incubated with IR680 (anti-mouse) or IR800 (anti-rabbit) conjugated secondary antibodies (1:10,000 dilution) for 1 h at RT. After 3 washes with TBST the membrane was imaged on a ChemiDoc Imaging System (Bio-Rad).
The following antibodies were used in this study: alpha-tubulin (mouse, Cell Signaling Technology, 3873S), GAPDH (mouse, proteintech, 60004-1-Ig), MBD2 (rabbit, Invitrogen, MA5-57472), Estrogen Receptor alpha (rabbit, Cell Signaling Technology, 8644S) Androgen Receptor (rabbit, Cell Signaling Technology, 5153S), IRDye 680RD goat anti-Mouse (LICOR 926–60870), and IRDye 800CW goat anti-rabbit (LICOR 926–32211).

**RT-qPCR**

20,000 cells were plated in 96-well tissue culture-treated plates (Corning 3513) and returned to the incubator to adhere overnight. The following day, compounds or DMSO control were added to the cells in biological triplicate, and plates were returned to the incubator for 24 h. The following day, medium was removed, and cells were lysed using Luna® Cell Ready Lysis Module (New England Biolabs, E3032S) according to the manufacturers protocol. RNA concentrations were determined using a NanoDrop Spectrophotometer (Thermo Fisher), and approx. 0.5 µg of RNA was used with Luna® Universal One-Step RT-qPCR Kit. RT-qPCR reactions were prepared in technical triplicate in TempPlate® 96-well PCR plates (USA Scientific) with 20 µL final volume. Thermal cycling was performed on a the CFX Connect Real-Time PCR Detection System

(BioRad) by using the manufacturers recommendation. Relative fold-change was determined using the 2^-dd$C_t$ method.

The following primers were used in this study: GAPDH (Hs.PT.39a.22214836), beta-Actin (Hs.PT.39a.22214847), ESR1 (Hs.PT.58.14846478), GREB1 (Hs.PT.58.26216464), MBD2 (Hs.PT.58.2050541), KLK3 (Hs.PT.58.20358116), FKBP5 (Hs.PT.58.39051416), CCNA2 (Hs.PT.56a.4535284), CDC20 (Hs.PT.58.20297042)

**Quantitative Tandem Mass Tagging (TMT)-Based Proteomic Profiling**

Cells were treated when they reached 70-80% confluency, with either DMSO vehicle or 12.5 µM of the respective TRACER for 24 hours. Total cell lysates were prepared using RIPA buffer supplemented with Benzonase® Nuclease (1:1000; Millipore, 70746) for 15 min at 37°C and protein concentration was determined using Pierce™ BCA Protein Assay Kit (Thermo Fisher) according to the manufacturers protocol. 100 µg of protein per replicate was reduced using 10 µM TCEP (37°C, 30 min) and alkylated using 20 µM iodoacetamide (room temperature, 30 min). Subsequently, protein samples were precipitated onto mixed hydrophilic/hydrophobic Sera-Mag™ Carboxylate-Modified Magnetic Beads (Cytiva) by the addition of EtOH to 80% (v/v). Beads were wasghed twice with 80% EtOH followed by resuspending the beads in 25 µl 8 M Urea in Tris-buffer pH 8.5. The sample was diluted to 2 M Urea and trypsin was added for overnight digestion (1:100, 37°C). After digestion, peptides were precipitated onto the magnetic beads by the addition of acetonitrile to >95% (v/v). Beads were washed twice with acetonitrile and digested peptides were eluted using $H_2O$ with 2% DMSO. 30 µg peptide per sample were labeled with TMTsixplex™ (Thermo Fisher Scientific, 90061), in accordance with the manufacturer’s protocol. TMT samples were then consolidated and fractionated using high pH reversed-phase peptide fractionation kits (Thermo Fisher Scientific, 84868) according to the manufacturer’s protocol. Fractions were vacuum concentrated, then reconstituted in 0.1% (v/v) formic acid and centrifuged at 20,000 g (5 min) in preparation for LC-MS/MS analysis.

Mass spectrometry analysis was performed on an Orbitrap Eclipse Tribrid Mass Spectrometer with a High Field Asymmetric Waveform Ion Mobility (FAIMS Pro) Interface (Thermo Fisher Scientific) with an UltiMate 3000 Nano Flow Rapid Separation LCnano System (Thermo Fisher Scientific). Offline fractionated samples (5 µL aliquot of 25 µL sample) were injected via an autosampler (Thermo Fisher Scientific) onto a 5 µL sample loop, which was subsequently eluted onto an Acclaim PepMap 100 C18 HPLC column (75 µm × 50 cm, NanoViper). The peptides were separated at a flow rate of 0.3 µL/min using the following gradient: 2% buffer B (100% acetonitrile with 0.1% formic acid) in buffer A (95:5 water/acetonitrile, 0.1% formic acid) for 5 min, followed by a gradient from 2−40% buffer B from 5−159 min, 40−95% buffer B from 159−160 min, held at 95% B from 160−179 min, 95% to 2% buffer B from 179−180 min, and then 2% buffer B from 180−200 min. The voltage applied to the nano-LC electrospray ionization source was 2.1 kV. Data were acquired through an MS1 master scan (Orbitrap analysis, resolution 120,000, 400−1800 m/z, RF lens 30%, heated capillary temperature 250 °C) with dynamic exclusion (repeat count 1, duration 60 sec). Data-dependent data acquisition comprised a full MS1 scan, followed by sequential MS2 scans based on 2 sec cycle times. FAIMS compensation voltages (CVs) of −35, −45, and −55 were applied. MS2 analysis consisted of a quadrupole isolation window of 0.7 m/z of the precursor ion followed by a higher energy collision dissociation (HCD) energy of 38% with an orbitrap resolution of 50,000.

Raw-files were analyzed using the Chaparral Platform and SagePro™. Trypsin cleavage specificity (cleavage at K, R, except if followed by P) allowed for up to 2 missed cleavages. Carbamidomethylation of cysteine residues (+57.02146) and TMT modification of peptide *N*-termini and lysine residues were set as static modification and methionine oxidation (+15.9949) was set as variable modification. MS1 tolerance was set to 10 ppm and MS2 tolerance to 300 ppm. Reporter-ion quantification was performed based on MS3 scans.

**Cellular Treatments for RNA-seq, CUT&RUN, and ATAC-seq Experiments**

T47D wild-type cells were grown in phenol-red-free RPMI supplemented with 10% charcoal-treated FBS (cFBS) for at 72h. The day before the experiment, cells were seeded into a 6-well tissue culture plate and cultured for 24 h. Afterwards, cells were treated with 1 nM E2 and DMSO vehicle or CS-1-103 (1.56 μM) for 24 h. 22Rv1 cells were cultured in 6-well tissue culture plates until 70-80% confluent and treated with DMSO vehicle or 12.5 μM CS-1-175 for 24 h in complete medium.

**RNAseq Analysis**

After treatment, RNA was isolated using the Monarch® Total RNA Miniprep Kit (NEB, T2010S). Total RNA quality, as well as poly-dT enriched mRNA quality, were assessed on an Agilent 2100 Bioanalyzer. Libraries were prepared using the KAPA mRNA Hyper Prep kit (Roche KK8581). Truncated universal stub adapters were ligated to cDNA fragments, which were then extended using 10 cycles of PCR using unique dual indexing primers into full length Illumina libraries. Library quality was checked on an AATI (now Agilent) Fragment Analyzer and transferred to the Vincent J. Coates Genomics Sequencing Laboratory (GSL), another QB3-Berkeley Core Research Facility at UC Berkeley.

FASTQ raw files were aligned and quantified using Kallisto[1] and filtered for >10 transcripts per million (tpm) for all replicates of at least one condition followed by differential gene expression analysis using DESeq2[2] and a threshold of $|\log2(\text{FoldChange})| > 1$ and an adjusted $p$-value $< 0.05$ was applied to identify differentially expressed genes. fGSEA analysis[3] was performed to identify significantly regulated MSigDB Hallmark gene sets[4]. Significantly downregulated genes were additionally compared to the ChEA 2022 TF-target dataset using Enrichr[5,6].

**CUT&RUN**

ER-binding sites in T47D were mapped using a modified CUT&RUN protocol described previously[7] with the anti-ESR1 (Epicypher, #13-2011) antibody. Sequenced CUT&RUN reads were processed as described by Skene et al.[8]

**ATAC-seq analysis**

Reads were counted over merged IDR-conservative ATAC-seq peaks in T47D across treatments, and peaks passing CPM > 5 in at least one sample group were retained. Differential expression was conducted using DEseq2. For visualization, a volcano plot was generated from filtered peaks, highlighting the top 500 downregulated peaks by p-value. Motif analysis on the top 500 downregulated peaks was performed using HOMER (size=given) with HOMER-generated background.

DEseq2 (1.46.0) was used for differential analysis. Rsubread (2.20.0) was used to count reads in peaks. HOMER (5.1) was used for motif analyses.

**Integrative data analysis**

Differential analysis results from ATAC-seq were annotated for overlap with high-confidence ER CUT&RUN sites (IDR-conservative – p-value ≥ 10-10). Differential expression from RNA-seq was integrated by joining promoter-annotated ATAC peaks to genes, producing a promoter-centric dataset. Binary indicators were defined as follows: ER binding site = 1 for overlaps; ATAC up or down = 1 if p-value ≤ 10-2 with sign determining the direction; RNA up or down = 1 if p-value ≤ 10-5 with sign determining the direction. Promoter entries with at least two active indicators (row sum ≥ 2) were retained and visualized with an UpSet plot to summarize intersections among ER binding and ATAC/RNA changes. For motif analysis, promoters were grouped by their combined indicator categories; HOMER was run with size="given" using a background of promoters lacking ER binding and nonsignificant in both ATAC (p-value > 5*10-2) and RNA (p-value > 5*10-2). Known-motif enrichments from ER-containing categories were combined, filtered to motifs meeting rank ≤ 5 and p-value ≤ 10−10, reshaped to a motif-by-category matrix (log odds ratios), and correlated with the binary category design to yield a point-biserial correlation matrix.

ComplexHeatmap (2.22.0) was used to plot the correlation heatmap. UpSetR (1.4.0) was used to plot the Upset plot. ggplot2 (4.0.0), cowplot (1.2.0) and ggtext (0.1.2) were used to plot the volcano plot. HOMER (5.1) was used for motif analyses.

### Synthetic Methods and Characterization

#### General Considerations

All chemical reactions were carried out under air with non-dry solvents, unless otherwise noted. Reagents were purchased at the highest commercial quality and used without further purification, unless otherwise stated.

Room temperature is defined as between 19-22 °C.

Reactions were stirred magnetically and monitored by thin layer chromatography (TLC) using TLC plates precoated with silica gel 60 F254 on aluminium (Merck KGaA). Detection was by UV (254 nm and 365 nm) or chemical stain (KMnO4, ninhydrin, iodine).

Solvents were removed in vacuo using a Buchi R-300 Rotavapor (equipped with an I-300 Pro Interface, B-300 Base Heating Bath, Welch 2037B-01 DryFast pump, and VWR AD15R-40-V11B Circulating Bath).

Automated flash chromatography was performed on a Biotage® Selekt instrument, equipped with a UV detector. Chromatograms were recorded at 254 and 280 nm.

High-resolution mass spectra (HRMS) were obtained on a Q Exactive Plus mass spectrometer (Thermo Fisher Scientific).

$^{1}$H and $^{13}$C Nuclear Magnetic Resonance (NMR) spectra were recorded on BRUKER NEO or a JEOUL spectrometer operating at 500 or 400 MHz for $^{1}$H and at 126 or 101 MHz for $^{13}$C NMR, respectively. Measurements were carried out at ambient temperature. Chemical shifts (δ) are reported in ppm with the residual solvent signal as internal standard (chloroform at 7.26 and 77.2 ppm for $^{1}$H NMR and $^{13}$C NMR, respectively). The multiplicity of each signal is indicated as s = singlet, d = doublet, t = triplet, q = quartet, quin = quintet, m = multiplet (i.e. complex peak obtained due to overlap). Coupling constants (J) are reported in Hertz (Hz). $^{13}$C NMR spectra were recorded with broadband $^{1}$H decoupling. Product peaks overlapping with the solvent signal were determined using two-dimensional $^{1}$H-$^{13}$C-HSQC NMR.

UPLC-UV/MS traces were recorded on either a Waters H-class or an Agilent 1290 Infinity II instrument. The Waters instrument was equipped with a quaternary solvent manager, a Waters autosampler, a Waters TUV detector and a Waters Acquity QDa detector with an Acquity UPLC BEH C18 1.7 µm, 2.1 x 50 mm RP column (Waters Corp., USA). Both instruments use A: 0.1 % TFA in H2O; B: 0.1% TFA in MeCN and
all methods have a flow rate of 0.6 mL/min. If not specified, the following gradient was used: 5% B 0.0 - 0.5 min, 5-95% B 0.5 - 3.0 min, 95% B 3.0 - 3.9 min, 5% B 3.9 - 5.0 min.

**General Procedure A** (Amide coupling)

A mixture of the corresponding carboxylic acid and HATU (1 equiv.) was dissolved in N,N-dimethylformamide (DMF) (0.1 M) followed by the addition of TEA (5 equiv.) was added and the reaction mixture stir for 2 minutes. The corresponding amine (1 equiv.) was dissolved in DMF and added to the reaction mixture. After 2 h at room temperature the reaction was purified by silica gel flash chromatography using a ternary gradient (Hexanes->Etylacetate->Methanol) on a Biotage® Selekt.

**General Procedure B** (Boc/tBu deprotection)

The Boc/tBu protected compound was dissolved in a minimal amount of MeOH and 4 M HCl in dioxane was added dropwise. After 1 h at r.t. the solvent was removed under a stream of nitrogen to obtain the desired compound as an HCl-salt. Deprotected compounds were used without further purification.

2-((((13*S*,*E*)-3-hydroxy-13-methyl-6,7,8,9,11,12,13,14,15,16-decahydro-17*H*-cyclopenta[*a*]phenanthren-17-ylidene)amino)oxy)acetic acid (**CS-1-65**):

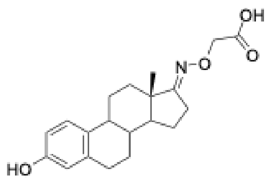

Estrone (500.0 mg, 1.85 mmol) and O-(carboxylmethyl)hydroxylamine (610 mg, 5.5 mmol) were dissolved in 16 ml dry pyridine and stirred overnight at room temperature. Afterwards the mixture was poured into 60 ml 10% HCl and extracted with EtOAc (3x 50 ml). The combined organic layers were washed with brine and dried over $MgSO_4$, filtered and evaporation before purification of the residue by flash column chromatography (DCM/MeOH 0-20%) to yield 605 mg of the desired product as a white powder (89% yield). Spectral data matched reported values from the literature.

**$^1$H NMR** (400 MHz, METHANOL-*D*$_4$) δ 7.57 (d, *J* = 2.9 Hz, 1H), 7.06 (d, *J* = 8.3 Hz, 1H), 6.55 (dd, *J* = 8.4, 2.7 Hz, 1H), 6.49 (s, 1H), 4.50 (t, *J* = 4.5 Hz, 2H), 2.81 – 2.74 (m, 1H), 2.56 (t, *J* = 6.1 Hz, 2H), 2.45 – 2.25 (m, 1H), 2.18 (d, *J* = 9.7 Hz, 1H), 2.06 – 1.83 (m, 3H), 1.71 – 1.25 (m, 7H), 0.91 (d, *J* = 2.9 Hz, 3H).
**$^{13}$C NMR** (101 MHz, METHANOL-*D*$_4$) δ 173.06, 156.57, 154.52, 137.67, 131.15, 126.15, 115.08, 112.68, 69.72, 52.89, 44.53, 43.97, 38.31, 33.93, 29.45, 27.24, 26.18, 25.98, 22.82, 16.87.

4-((4-(pyridin-2-yl)thiazol-2-yl)amino)phenol (**CS-1-69**):

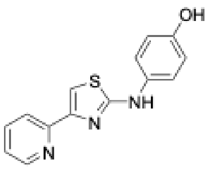

The target compound was synthesized via the Hantzsch thiazole synthesis. 2-bromo-1-(pyridin-2-yl)ethan-1-one (1 mmol) and 1-(4-hydroxyphenyl)thiourea (1 mmol) were dissolved in 20 ml EtOH (20 mL) and stirred at reflux for 1 h. The mixture was cooled to r.t., diluted with water (50 mL), and the pH was adjusted to approx. 8. After 2 h, the formed precipitate was filtered, washed with water and purified by flash chromatography (Hex/EtOAc 0−50%) to give the thiazole in 95% yield. (255 mg)

**$^1$H NMR** (400 MHz, $CDCl_3$/METHANOL-*D*$_4$) δ 8.47 (d, *J* = 2.7 Hz, 1H), 7.97 (d, *J* = 8.0 Hz, 1H), 7.79 (td, *J* = 7.8, 1.8 Hz, 1H), 7.34 (d, *J* = 8.7 Hz, 2H), 7.25 (s, 1H), 7.22 (d, *J* = 6.0 Hz, 1H), 6.81 (d, *J* = 8.9 Hz, 2H).
**$^{13}$C NMR** (101 MHz, $CDCl_3$/METHANOL-*D*$_4$) δ 167.74, 153.59, 153.06, 150.58, 149.16, 138.11, 133.82, 123.09, 121.95, 121.78 (2C), 116.30 (2C), 106.17.
**HRMS** for $C_{14}H_{11}N_3OS$ $[M+H]^+$ calc.: 270.0696 Da; found: 270.0691 Da

benzyl(6-(2-((((13*S*,*E*)-3-hydroxy-13-methyl-6,7,8,9,11,12,13,14,15,16-decahydro-17*H*-cyclopenta[*a*] phenanthren-17-ylidene)amino)oxy)acetamido)hexyl)carbamate (**CS-1-96**):

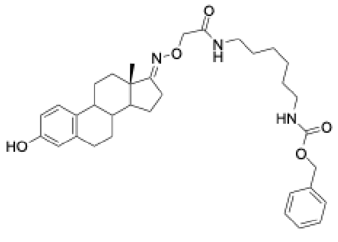

Was prepared according to **General Procedure A** starting from **CS-1-65** (10 mg) and benzyl (6-aminohexyl)carbamate. (3 mg, 18% yield)

**$^1$H NMR** (400 MHz, DMSO-$D_6$) δ 9.00 (s, 1H), 7.95 (s, 1H), 7.39 (t, *J* = 5.8 Hz, 1H), 7.37 – 7.24 (m, 6H), 7.20 (t, *J* = 5.6 Hz, 1H), 7.04 (d, *J* = 8.5 Hz, 1H), 6.50 (dd, *J* = 8.5, 3.0 Hz, 1H), 6.44 (s, 1H), 4.97 (s, 2H), 4.30 (s, 2H), 3.09 (q, *J* = 6.8 Hz, 2H), 2.96 (q, *J* = 6.6 Hz, 2H), 2.28 (d, *J* = 10.3 Hz, 1H), 2.14 (s, 1H), 1.85 (s, 3H), 1.53 – 1.42 (m, 2H), 1.37 (s, 11H), 1.23 (s, 4H), 0.87 (s, 3H).
**$^{13}$C NMR** (101 MHz, DMSO-$D_6$) δ 171.92, 169.41, 162.84, 156.60, 155.53, 137.84, 137.58, 130.55, 128.85, 128.24, 126.55, 115.47, 113.31, 72.78, 65.58, 52.87, 44.59, 44.04, 38.78, 38.52, 38.33, 36.31, 34.48, 31.31, 29.95, 29.75, 29.58, 26.49, 26.35, 23.06, 17.65.
**HRMS** for $C_{34}H_{45}N_3O_5$ $[M+H]^+$ calc.: 576.3432 Da; found: 576.3434 Da

*N*-(2-(3-(but-3-yn-1-yl)-3*H*-diazirin-3-yl)ethyl)-3-((4-(pyridin-2-yl)thiazol-2-yl)amino) benzamide (**CS-1-102**):

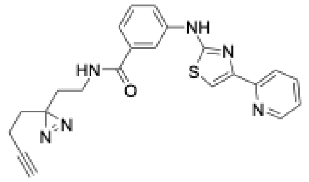

Was prepared according to **General Procedure A** starting from 3-((4-(pyridin-2-yl)thiazol-2-yl)amino)benzoic acid (8 mg) and 2-(3-(but-3-yn-1-yl)-3*H*-diazirin-3-yl)ethan-1-amine. (7 mg, 61% yield)

**$^1$H NMR** (400 MHz, DMSO-$D_6$) δ 10.42 (s, 1H), 8.55 (d, *J* = 4.6 Hz, 1H), 8.39 (t, *J* = 5.6 Hz, 1H), 8.13 (t, *J* = 2.0 Hz, 1H), 7.99 (d, *J* = 7.9 Hz, 1H), 7.95 – 7.80 (m, 2H), 7.54 (s, 1H), 7.48 – 7.33 (m, 2H), 7.33 – 7.23 (m, 1H), 3.14 (q, *J* = 6.7 Hz, 2H), 2.79 (t, *J* = 2.6 Hz, 1H), 2.65 (s, 1H), 1.98 (td, *J* = 7.4, 2.6 Hz, 2H), 1.62 (dt, *J* = 12.7, 7.2 Hz, 4H).
**$^{13}$C NMR** (101 MHz, DMSO-$D_6$) δ 166.99, 163.75, 152.62, 150.88, 149.96, 141.73, 137.76, 136.17, 129.51, 123.22, 120.92, 120.12, 119.85, 116.62, 107.68, 83.70, 72.32, 34.97, 32.53, 31.85, 27.86, 13.25.
**HRMS** for $C_{22}H_{21}N_6OS$ $[M+H]^+$ calc.: 417.1492 Da; found: 417.1456 Da

2-((((13*S*,*E*)-3-hydroxy-13-methyl-6,7,8,9,11,12,13,14,15,16-decahydro-17*H*-cyclopenta [*a*]phenanthren-17-ylidene)amino)oxy)-*N*-(6-(4-((4-(pyridin-2-yl)thiazol-2-yl)amino) phenoxy)hexyl)acetamide (**CS-1-85**):

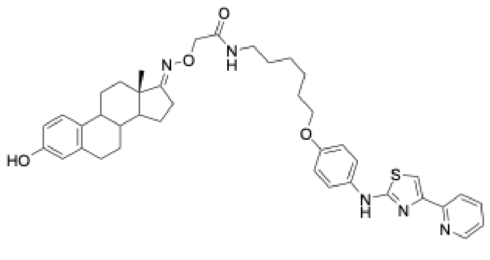

Was prepared according to **General Procedure A** starting from **CS-1-65** and **CS-1-71**. (6.3 mg, 63% yield)

**$^1$H NMR** (500 MHz, DMSO) δ 10.10 (s, 1H), 8.57 (dt, *J* = 4.7, 1.4 Hz, 1H), 8.28 (s, 1H), 8.05 – 7.95 (m, 1H), 7.88 (td, *J* = 7.7, 1.8 Hz, 1H), 7.72 – 7.55 (m, 2H), 7.47 (s, 1H), 7.42 (t, *J* = 6.0 Hz, 1H), 7.31 (ddd, *J* = 7.5, 4.8, 1.2 Hz, 1H), 7.02 (d, *J* = 8.5 Hz, 1H), 6.94 – 6.87 (m, 2H), 6.49 (dd, *J* = 8.4, 2.7 Hz, 1H), 6.43 (d, *J* = 2.6 Hz, 1H), 4.31 (s, 2H), 3.92 (t, *J* = 6.5 Hz, 2H), 3.13 (dq, *J* = 13.4, 6.6 Hz, 2H), 2.80 – 2.64

(m, 2H), 2.55 (s, 2H), 2.28 (dd, *J* = 13.9, 3.6 Hz, 1H), 2.14 (t, *J* = 10.3 Hz, 1H), 1.86 (ddt, *J* = 17.4, 14.1, 3.5 Hz, 2H), 1.69 (p, *J* = 6.7 Hz, 2H), 1.56 – 1.22 (m, 14H), 0.87 (s, 3H).
**HRMS** for $C_{40}H_{47}N_5O_4S$ $[M+H]^+$ calc.: 694.3422 Da; found: 694.3416 Da
**Retention time Gradient A**: 2.675 min

2-((((13*S*,*E*)-3-hydroxy-13-methyl-6,7,8,9,11,12,13,14,15,16-decahydro-17*H*-cyclopenta [*a*]phenanthren-17-ylidene)amino)oxy)-*N*-(2-(2-(2-(2-(4-((4-(pyridin-2-yl)thiazol-2-yl)amino)phenoxy)ethoxy)ethoxy)ethoxy) ethyl)acetamide (**CS-1-86**):

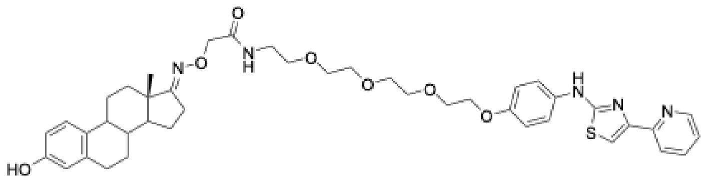

Was prepared according to **General Procedure A** starting from **CS-1-65** and **CS-1-76**. (10 mg, 51% yield)

**[1]H NMR** (400 MHz, DMSO-$D_6$) δ 10.06 (s, 1H), 8.95 (s, 1H), 8.53 (d, *J* = 4.6 Hz, 1H), 7.93 (d, *J* = 7.9 Hz, 1H), 7.84 (td, *J* = 7.7, 1.7 Hz, 1H), 7.59 (d, *J* = 9.0 Hz, 2H), 7.43 (s, 1H), 7.35 (t, *J* = 5.6 Hz, 1H), 7.31 – 7.22 (m, 1H), 6.99 (d, *J* = 8.5 Hz, 1H), 6.90 (d, *J* = 9.0 Hz, 2H), 6.46 (dd, *J* = 8.4, 2.4 Hz, 1H), 6.39 (d, *J* = 2.4 Hz, 1H), 4.29 (s, 2H), 4.01 (d, *J* = 4.8 Hz, 2H), 3.69 (dd, *J* = 5.3, 3.7 Hz, 2H), 3.59 – 3.44 (m, 8H), 3.39 (t, *J* = 6.0 Hz, 2H), 3.29 – 3.18 (m, 2H), 3.13 (d, *J* = 5.2 Hz, 1H), 2.74 – 2.64 (m, 2H), 2.32 – 2.19 (m, 1H), 2.10 (t, *J* = 8.1 Hz, 1H), 1.89 – 1.76 (m, 3H), 1.51 – 1.41 (m, 1H), 1.40 – 1.20 (m, 6H), 0.83 (s, 3H).
**[13]C NMR** (101 MHz, $CDCl_3$) δ 170.65, 169.16, 155.58, 154.08, 153.97, 149.39, 143.32, 138.16, 137.12, 136.96, 133.83, 126.47 (2C), 122.63, 121.76 (2C), 120.99, 118.67, 115.65 (2C), 113.23, 106.03, 72.69, 70.88, 70.68, 70.60, 70.36, 70.11, 69.88, 67.91, 53.00, 44.73, 44.65, 44.03, 38.76, 38.24, 36.58, 34.16, 29.55, 27.31, 26.19, 23.09.
**HRMS** for $C_{42}H_{51}N_5O_7S$ $[M+H]^+$ calc.: 769.3509 Da; found: 770.3580 Da

*N*-(6-(2-((((13*S*,*E*)-3-hydroxy-13-methyl-6,7,8,9,11,12,13,14,15,16-decahydro-17*H*-cyclopenta [*a*]phenanthren-17-ylidene)amino)oxy)acetamido)hexyl)-3-((4-(pyridin-2-yl)thiazol-2-yl)amino)benzamide (**CS-1-103):**

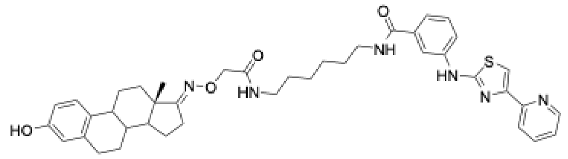

Was prepared according to **General Procedure A** starting from **CS-1-65** and **CS-1-99**. (3.4 mg, 47% yield)

**[1]H NMR** (400 MHz, DMSO-$D_6$) δ 10.43 (s, 1H), 8.55 (d, *J* = 4.9 Hz, 1H), 8.36 (d, *J* = 5.0 Hz, 2H), 8.15 (s, 1H), 7.99 (d, *J* = 7.9 Hz, 1H), 7.94 – 7.78 (m, 2H), 7.53 (s, 1H), 7.37 (dd, *J* = 14.1, 5.7 Hz, 3H), 7.28 (dd, *J* = 7.5, 4.8 Hz, 1H), 6.96 (d, *J* = 8.4 Hz, 1H), 6.44 (dd, *J* = 8.5, 2.6 Hz, 1H), 6.39 (d, *J* = 2.7 Hz, 1H), 4.27 (s, 2H), 3.21 (q, *J* = 6.6 Hz, 2H), 3.07 (q, *J* = 6.5 Hz, 2H), 2.75 – 2.59 (m, 2H), 2.50 (m, 2H), 2.23 (d, *J* = 12.9 Hz, 1H), 2.08 (s, 2H), 1.87 – 1.73 (m, 3H), 1.48 (dt, *J* = 10.4, 5.3 Hz, 3H), 1.43 – 1.21 (m, 11H), 0.82 (s, 3H).
**[13]C NMR** (101 MHz, DMSO-$D_6$) δ 171.93, 169.46, 166.87, 163.75, 155.52, 152.64, 149.94, 142.20, 141.67, 137.73, 137.55, 136.42, 132.45, 130.61, 126.53, 123.21, 120.91, 120.11, 119.66, 116.64, 115.46, 113.31,

107.62, 76.04, 52.84, 44.58, 44.02, 38.51, 38.31, 34.46, 29.74 (3C), 29.56, 27.74, 27.37, 26.74, 26.51, 26.34 (2C), 17.64.
**HRMS** for $C_{41}H_{49}N_6O_4S$ $[M+H]^+$ calc.: 721.3531 Da; found: 721.3525 Da

2-((((13*S*,*E*)-3-hydroxy-13-methyl-6,7,8,9,11,12,13,14,15,16-decahydro-17*H*-cyclopenta [*a*]phenanthren-17-ylidene)amino)oxy)-1-(4-(3-((4-(pyridin-2-yl)thiazol-2-yl)amino) benzoyl)piperazin-1-yl)ethan-1-one (**CS-1-163**):

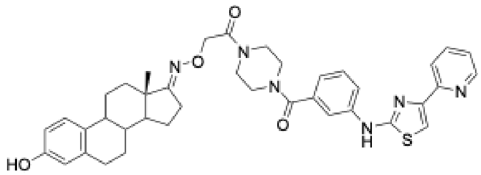

Was prepared according to **General Procedure A** starting from **CS-1-65** and **CS-1-158**. (5.6 mg, 81% yield)

**$^1$H NMR** (500 MHz, DMSO) δ 10.51 (s, 1H), 9.02 (s, 1H), 8.59 (dd, *J* = 5.0, 1.6 Hz, 1H), 7.99 (d, *J* = 7.8 Hz, 1H), 7.92 (td, *J* = 7.7, 1.8 Hz, 1H), 7.82 (d, *J* = 8.0 Hz, 2H), 7.62 (s, 1H), 7.45 (t, *J* = 7.9 Hz, 1H), 7.34 (dd, *J* = 7.4, 4.8 Hz, 1H), 7.03 (d, *J* = 7.9 Hz, 2H), 6.51 (dd, *J* = 8.4, 2.6 Hz, 1H), 6.44 (d, *J* = 2.6 Hz, 1H), 4.65 (s, 2H), 3.64 – 3.46 (m, 4H), 2.80 – 2.67 (m, 2H), 2.44 (s, 1H), 2.27 (d, *J* = 13.0 Hz, 1H), 2.14 (s, 1H), 1.94 – 1.73 (m, 4H), 1.55 – 1.20 (m, 6H), 0.85 (s, 3H). (Proton signal of two piperazine $CH_2$-groups overlaps with $H_2O$ peak)
**HRMS** for $C_{39}H_{42}N_6O_4S$ $[M+H]^+$ calc.: 691.3061 Da; found: 691.3054 Da
**Retention time Gradient A**: 2.403 min

*N*-(5-(2-((((13*S*,*E*)-3-hydroxy-13-methyl-6,7,8,9,11,12,13,14,15,16-decahydro-17*H*-cyclopenta [*a*]phenanthren-17-ylidene)amino)oxy)acetamido)pentyl)-3-((4-(pyridin-2-yl)thiazol-2-yl)amino)benzamide (**CS-1-164**):

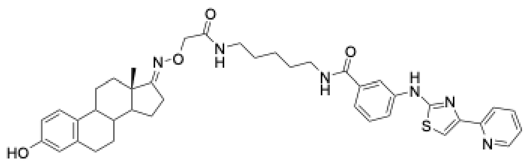

Was prepared according to **General Procedure A** starting from **CS-1-65** and **CS-1-157**. (5.2 mg, 73% yield)

**$^1$H NMR** (500 MHz, DMSO) δ 10.47 (s, 1H), 8.66 – 8.56 (m, 1H), 8.41 (t, *J* = 5.6 Hz, 1H), 8.31 (s, 1H), 8.19 (d, *J* = 2.4 Hz, 1H), 8.03 (d, *J* = 7.8 Hz, 1H), 7.98 – 7.85 (m, 2H), 7.57 (s, 1H), 7.43 (dd, *J* = 13.7, 6.1 Hz, 3H), 7.32 (dd, *J* = 7.6, 4.5 Hz, 1H), 7.01 (d, *J* = 8.6 Hz, 1H), 6.51 – 6.41 (m, 2H), 4.31 (s, 2H), 3.26 (q, *J* = 6.8 Hz, 3H), 3.13 (q, *J* = 6.7 Hz, 2H), 2.76 – 2.57 (m, 3H), 2.34 – 2.22 (m, 1H), 2.13 (s, 1H), 1.93 – 1.77 (m, 4H), 1.60 – 1.24 (m, 12H), 0.86 (s, 3H).
**HRMS** for $C_{40}H_{46}N_6O_4S$ $[M+H]^+$ calc.: 707.3374 Da; found: 707.3363 Da
**Retention time Gradient A**: 2.408 min

*N*-(3-(2-((((13*S*,*E*)-3-hydroxy-13-methyl-6,7,8,9,11,12,13,14,15,16-decahydro-17*H*-cyclopenta [*a*]phenanthren-17-ylidene)amino)oxy)acetamido)propyl)-3-((4-(pyridin-2-yl)thiazol-2-yl)amino)benzamide (**CS-1-165**):

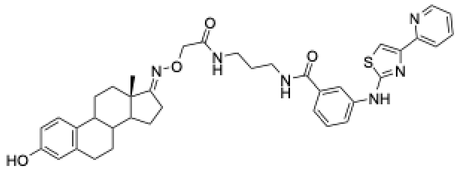

Was prepared according to **General Procedure A** starting from **CS-1-65** and **CS-1-160**. (5.3 mg, 78% yield)

**$^{1}$H NMR** (500 MHz, DMSO) δ 10.47 (s, 1H), 8.59 (dd, *J* = 4.9, 1.7 Hz, 1H), 8.48 (t, *J* = 5.8 Hz, 1H), 8.28 (d, *J* = 12.4 Hz, 2H), 8.03 (d, *J* = 7.9 Hz, 1H), 7.97 – 7.82 (m, 2H), 7.68 (t, *J* = 6.2 Hz, 1H), 7.58 (s, 1H), 7.41 (d, *J* = 7.4 Hz, 2H), 7.32 (dd, *J* = 7.6, 4.8 Hz, 1H), 6.97 (d, *J* = 8.5 Hz, 1H), 6.48 (dd, *J* = 8.4, 2.7 Hz, 1H), 6.41 (d, *J* = 2.6 Hz, 1H), 4.36 (s, 2H), 3.35 – 3.29 (m, 2H), 3.21 (q, *J* = 6.4 Hz, 2H), 2.76 – 2.57 (m, 3H), 2.23 (dq, *J* = 13.2, 3.7 Hz, 1H), 2.08 (d, *J* = 11.8 Hz, 1H), 1.94 – 1.74 (m, 3H), 1.67 (p, *J* = 6.8 Hz, 2H), 1.50 (td, *J* = 13.2, 4.0 Hz, 1H), 1.43 – 1.18 (m, 5H), 0.85 (s, 3H).
**HRMS** for $C_{38}H_{42}N_6O_4S$ $[M+H]^+$ calc.: 679.3061 Da; found: 679.3057 Da
**Retention time Gradient A**: 2.329 min

*N*-(12-(2-((((13*S*,*E*)-3-hydroxy-13-methyl-6,7,8,9,11,12,13,14,15,16-decahydro-17*H*-cyclopenta [*a*]phenanthren-17-ylidene)amino)oxy)acetamido)dodecyl)-3-((4-(pyridin-2-yl)thiazol-2-yl)amino)benzamide (**CS-1-166**):

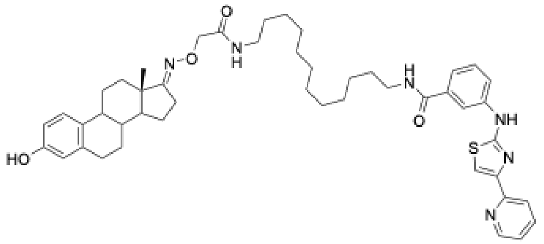

Was prepared according to **General Procedure A** starting from **CS-1-65** and **CS-1-159**. (4.3 mg, 51% yield)

**$^{1}$H NMR** (400 MHz, DMSO-*D*$_6$) δ 10.42 (s, 1H), 8.55 (d, *J* = 4.8 Hz, 1H), 8.44 – 8.31 (m, 1H), 8.15 (s, 1H), 7.99 (d, *J* = 7.9 Hz, 1H), 7.93 – 7.79 (m, 2H), 7.53 (s, 1H), 7.45 – 7.23 (m, 4H), 6.99 (d, *J* = 8.5 Hz, 1H), 6.47 (dd, *J* = 8.4, 2.6 Hz, 1H), 6.40 (d, *J* = 2.6 Hz, 1H), 4.26 (s, 2H), 3.20 (t, *J* = 6.6 Hz, 2H), 3.06 (ddq, *J* = 19.6, 13.1, 6.6 Hz, 2H), 2.70 (d, *J* = 13.7 Hz, 2H), 2.51 (d, *J* = 8.6 Hz, 1H), 2.24 (d, *J* = 13.0 Hz, 1H), 2.09 (d, *J* = 9.9 Hz, 1H), 1.89 – 1.73 (m, 3H), 1.52 – 1.43 (m, 3H), 1.37 – 1.15 (m, 24H), 0.83 (s, 3H).
**HRMS** for $C_{44}H_{51}F_3N_8O_4S$ $[M+H]^+$ calc.: 845.3779 Da; found: 845.3772 Da
**Retention time Gradient A**: 2.962 min

2-((((13*S*,*E*)-3-hydroxy-13-methyl-6,7,8,9,11,12,13,14,15,16-decahydro-17*H*-cyclopenta[*a*]phenanthren-17-ylidene)amino)oxy)-1-(9-(3-((4-(pyridin-2-yl)thiazol-2-yl)amino)benzoyl)-3,9-diazaspiro[5.5]undecan-3-yl)ethan-1-one (**CS-1-167**):

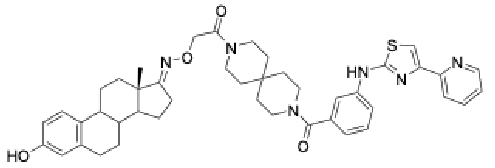

Was prepared according to **General Procedure A** starting from **CS-1-65** and **CS-1-161**. (2.9 mg, 62% yield)

**$^{1}$H NMR** (400 MHz, DMSO-*D*$_6$) δ 10.42 (s, 1H), 8.95 (s, 1H), 8.56 (d, *J* = 4.7 Hz, 1H), 7.92 (d, *J* = 7.8 Hz, 1H), 7.86 (t, *J* = 7.9 Hz, 1H), 7.77 (s, 1H), 7.72 (d, *J* = 8.1 Hz, 1H), 7.55 (s, 1H), 7.37 (t, *J* = 7.8 Hz, 1H), 7.29 (dd, *J* = 7.3, 4.8 Hz, 1H),

6.99 (d, $J$ = 8.5 Hz, 1H), 6.93 (d, $J$ = 7.5 Hz, 1H), 6.50 – 6.43 (m, 1H), 6.39 (d, $J$ = 2.6 Hz, 1H), 4.56 (s, 2H), 3.59 (s, 2H), 3.40 – 3.33 (m, 5H), 2.84 – 2.63 (m, 2H), 2.50 (s, 1H), 2.43 – 2.35 (m, 1H), 2.31 – 2.20 (m, 1H), 2.10 (s, 1H), 1.89 – 1.71 (m, 3H), 1.59 – 1.17 (m, 18H), 0.82 (s, 3H).

**HRMS** for $C_{44}H_{50}N_6O_4S$ $[M+H]^+$ calc.: 759.3687 Da; found: 759.3679 Da

**Retention time Gradient A**: 2.530 min

*N*-(6-(4-(3-(4-cyano-3-(trifluoromethyl)phenyl)-5,5-dimethyl-4-oxo-2-thioxoimidazolidin-1-yl)butanamido)hexyl)-3-((4-(pyridin-2-yl)thiazol-2-yl)amino)benzamide (**CS-1-175):**

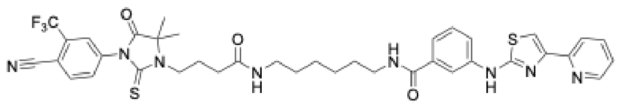

Was prepared according to **General Procedure A** starting from **CS-1-170** and **CS-1-99**. (8.6 mg, 56% yield)

**$^{1}$H NMR** (400 MHz, DMSO-*D*$_6$) δ 10.41 (s, 1H), 8.55 (d, *J* = 2.7 Hz, 1H), 8.38 (t, *J* = 5.7 Hz, 1H), 8.30 (d, *J* = 8.5 Hz, 1H), 8.19 (s, 1H), 8.14 (s, 1H), 7.98 (dd, *J* = 11.1, 7.9 Hz, 2H), 7.91 – 7.82 (m, 2H), 7.79 (t, *J* = 5.7 Hz, 1H), 7.54 (s, 1H), 7.38 (d, *J* = 7.1 Hz, 2H), 7.29 (dd, *J* = 7.8, 4.6 Hz, 1H), 3.67 – 3.58 (m, 2H), 3.22 (d, *J* = 6.4 Hz, 2H), 3.01 (q, *J* = 6.5 Hz, 2H), 2.13 (t, *J* = 7.4 Hz, 2H), 1.97 – 1.89 (m, 2H), 1.49 (s, 6H), 1.43 – 1.17 (m, 8H).
**$^{13}$C NMR** (101 MHz, DMSO-*D*$_6$) δ 178.73, 175.83, 171.63, 166.88, 163.76, 152.59, 150.83, 149.91, 141.68, 138.66, 137.79, 136.54, 136.44, 134.52, 131.50, 129.44, 128.59, 123.23, 122.76, 120.92, 120.14, 119.69, 116.65, 115.59, 108.91, 107.69, 65.72, 43.54, 38.96, 33.08, 29.68 (2C), 26.75, 26.73, 24.43 (2C), 22.71.
**HRMS** for $C_{38}H_{39}F_3N_8O_3S_2$ [M+H]$^+$ calc.: 777.2611 Da; found: 777.2606 Da

*N*-(2-(2-(2-(4-(3-(4-cyano-3-(trifluoromethyl)phenyl)-5,5-dimethyl-4-oxo-2-thioxoimidazolidin-1-yl)butanamido)ethoxy)ethoxy)ethyl)-3-((4-(pyridin-2-yl)thiazol-2-yl)amino)benzamide (**CS-1-176**):

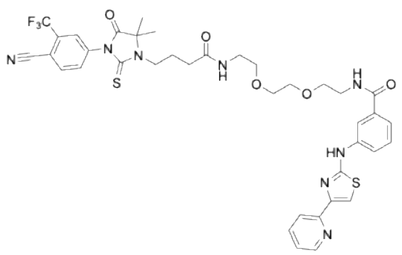

Was prepared according to **General Procedure A** starting from **CS-1-170** and **CS-1-138**. (9.8 mg, 61% yield)

**$^{1}$H NMR** (400 MHz, DMSO-*D*$_6$) δ 10.42 (s, 1H), 8.61 – 8.52 (m, 1H), 8.43 (t, *J* = 5.7 Hz, 1H), 8.30 (d, *J* = 8.2 Hz, 1H), 8.23 – 8.17 (m, 1H), 8.15 (t, *J* = 1.8 Hz, 1H), 7.98 (ddd, *J* = 12.3, 8.1, 1.5 Hz, 2H), 7.88 (dtd, *J* = 16.0, 6.9, 2.6 Hz, 2H), 7.54 (s, 1H), 7.46 – 7.36 (m, 2H), 7.35 – 7.25 (m, 1H), 3.68 – 3.57 (m, 2H), 3.56 – 3.44 (m, 6H), 3.39 (dt, *J* = 12.0, 5.8 Hz, 4H), 3.17 (q, *J* = 5.8 Hz, 2H), 2.14 (t, *J* = 7.5 Hz, 2H), 1.99 – 1.88 (m, 3H), 1.49 (s, 6H).
**$^{13}$C NMR** (101 MHz, DMSO-*D*$_6$) δ 178.73, 175.84, 171.96, 167.02, 163.74, 152.61, 150.86, 149.93, 141.73, 138.66, 137.79, 136.54, 136.04, 134.52, 131.50, 129.49, 128.59, 123.22, 122.76, 120.92, 120.11, 119.85, 116.71, 115.60, 108.91, 107.68, 70.09 (2C), 69.69, 69.46, 65.72, 43.51, 39.06, 32.97, 24.33, 22.70 (2C).
**HRMS** for $C_{38}H_{39}F_3N_8O_5S_2$ [M+H]$^+$ calc.: 809.2510 Da; found: 809.2509 Da

*N*-(12-(4-(3-(4-cyano-3-(trifluoromethyl)phenyl)-5,5-dimethyl-2,4-dioxoimidazolidin-1-yl)butanamido) dodecyl)-3-((4-(pyridin-2-yl)thiazol-2-yl)amino)benzamide (**CS-1-169**):

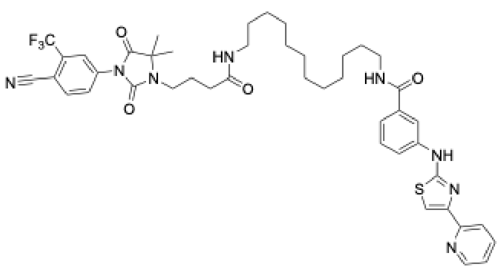

Was prepared according to **General Procedure A** starting from **CS-1-154**. (6.7 mg, 81% yield)

**$^1$H NMR** (400 MHz, DMSO-*D*$_6$) δ 10.41 (s, 1H), 8.55 (ddd, *J* = 4.8, 1.8, 0.9 Hz, 1H), 8.37 (t, *J* = 5.6 Hz, 1H), 8.26 (d, *J* = 8.4 Hz, 1H), 8.15 (d, *J* = 2.2 Hz, 2H), 7.99 (dt, *J* = 8.2, 1.8 Hz, 2H), 7.89 – 7.81 (m, 2H), 7.72 (t, *J* = 5.6 Hz, 1H), 7.53 (s, 1H), 7.46 – 7.33 (m, 2H), 7.33 – 7.23 (m, 1H), 3.30 – 3.15 (m, 4H), 2.97 (q, *J* = 6.6 Hz, 2H), 2.10 (t, *J* = 7.5 Hz, 2H), 1.90 – 1.72 (m, 2H), 1.60 – 1.45 (m, 2H), 1.42 (d, *J* = 4.5 Hz, 8H), 1.28 (d, *J* = 27.5 Hz, 6H), 1.18 (s, 10H).

**$^{13}$C NMR** (101 MHz, DMSO-*D*$_6$) δ 175.21, 171.81, 166.90, 163.74, 162.86, 153.06, 152.62, 150.86, 149.94, 141.67, 137.72, 137.39, 136.56, 136.45, 130.46, 129.42, 124.57, 123.19, 120.91, 120.13, 119.66, 116.62, 115.75, 107.64, 107.17, 62.16, 38.98, 33.11, 29.66 (2C), 29.53 (4C), 29.32, 29.29, 27.00, 26.95, 25.68, 23.06 (2C). two carbons are overlapping with the solvent peak.

**HRMS** for $C_{44}H_{51}F_3N_8O_4S$ $[M+H]^+$ calc.: 845.3779 Da; found: 845.3772 Da

*N*-(2-(2-(2-(4-(3-(4-cyano-3-(trifluoromethyl)phenyl)-5,5-dimethyl-2,4-dioxoimidazolidin-1-yl)butanamido) ethoxy)ethoxy)ethyl)-3-((4-(pyridin-2-yl)thiazol-2-yl)amino)benzamide (**CS-1-168**):

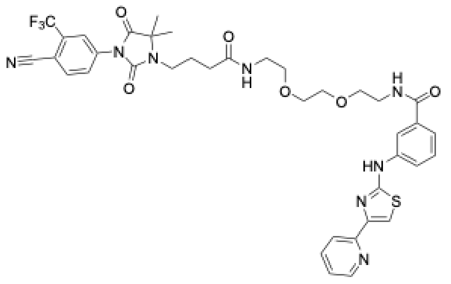

Was prepared according to **General Procedure A** starting from **CS-1-154**. (7.4 mg 92% yield)

**$^1$H NMR** (400 MHz, DMSO-*D*$_6$) δ 10.42 (s, 1H), 8.53 (dd, *J* = 4.9, 1.6 Hz, 1H), 8.43 (s, 1H), 8.23 (d, *J* = 8.2 Hz, 1H), 8.19 – 8.09 (m, 2H), 7.98 (td, *J* = 5.3, 2.5 Hz, 2H), 7.92 – 7.81 (m, 3H), 7.52 (s, 1H), 7.38 (d, *J* = 7.0 Hz, 2H), 7.28 (d, *J* = 12.4 Hz, 1H), 3.38 (dt, *J* = 16.2, 5.6 Hz, 4H), 3.27 (dt, *J* = 21.8, 7.5 Hz, 2H), 3.15 (q, *J* = 5.8 Hz, 2H), 2.11 (t, *J* = 7.5 Hz, 2H), 1.84 – 1.74 (m, 2H), 1.39 (s, 6H).

**$^{13}$C NMR** (101 MHz, DMSO-*D*$_6$) δ 175.26, 172.38, 167.17, 163.75, 157.3, 153.06, 152.57, 150.80, 149.91, 141.69, 137.85, 137.31, 136.53, 135.94, 131.47, 130.45, 129.54, 124.52, 123.31, 120.98, 120.10, 119.92, 116.62, 115.74, 107.68, 104.62, 70.04 (2C), 69.58, 69.38, 62.18, 39.04, 33.01, 25.56, 23.01(2C). (two carbon overlap with solvent peak)

**HRMS** for $C_{38}H_{39}F_3N_8O_6S$ $[M+H]^+$ calc.: 793.2738 Da; found: 793.2729 Da

*N*-(6-(4-(3-(4-cyano-3-(trifluoromethyl)phenyl)-5,5-dimethyl-2,4-dioxoimidazolidin-1-yl)butanamido)hexyl)-3-((4-(pyridin-2-yl)thiazol-2-yl)amino)benzamide (**CS-1-162**):

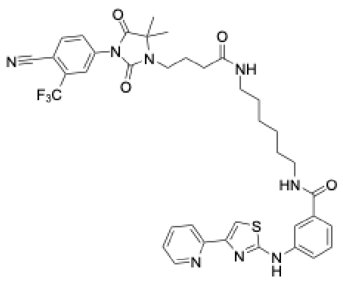

Was prepared according to **General Procedure A** starting from **CS-1-154**. (8.3 mg 73% yield**)**

**$^{1}$H NMR** (400 MHz, DMSO-*D*$_6$) δ 10.41 (s, 1H), 8.55 (dd, *J* = 4.9, 1.7 Hz, 1H), 8.38 (t, *J* = 5.6 Hz, 1H), 8.27 (dd, *J* = 8.5, 4.6 Hz, 1H), 8.19 – 8.11 (m, 2H), 8.05 – 7.96 (m, 2H), 7.95 – 7.80 (m, 2H), 7.75 (t, *J* = 5.6 Hz, 1H), 7.53 (s, 1H), 7.38 (d, *J* = 7.4 Hz, 2H), 7.28 (dd, *J* = 7.5, 5.0 Hz, 1H), 3.29 – 3.18 (m, 2H), 3.00 (q, *J* = 6.5 Hz, 2H), 2.11 (t, *J* = 7.6 Hz, 2H), 1.80 (q, *J* = 7.8 Hz, 3H), 1.54 – 1.23 (m, 14H).
**$^{13}$C NMR** (101 MHz, DMSO-*D*$_6$) δ 175.21, 171.83, 166.88, 163.75, 153.06, 152.62, 150.87, 149.95, 143.87, 141.67, 137.73, 137.40, 136.56, 136.42, 130.47, 129.43, 124.58, 124.14, 123.21, 120.90, 120.13, 119.67, 116.64, 115.75, 107.64, 107.18, 62.16, 38.92, 33.12, 29.67, 26.70 (2C), 25.70, 23.06 (2C). (two carbon overlap with solvent peak)
**HRMS** for $C_{38}H_{39}F_3N_8O_4S$ $[M+H]^+$ calc.: 761.2840 Da; found: 761.2834 Da

1-((*S*)-*sec*-butyl)-*N*-((4,6-dimethyl-2-oxo-1,2-dihydropyridin-3-yl)methyl)-6-(6-(4-(3-(2-(2-(2-((((13*S*,*E*)-3-hydroxy-13-methyl-6,7,8,9,11,12,13,14,15,16-decahydro-17*H*-cyclo-penta[*a*]phenanthren-17-ylidene)amino)oxy)acetamido)ethoxy)ethoxy)propanoyl) piperazin-1-yl)pyridin-3-yl)-3-methyl-1*H*-indole-4-carboxamide (**CS-1-123**):

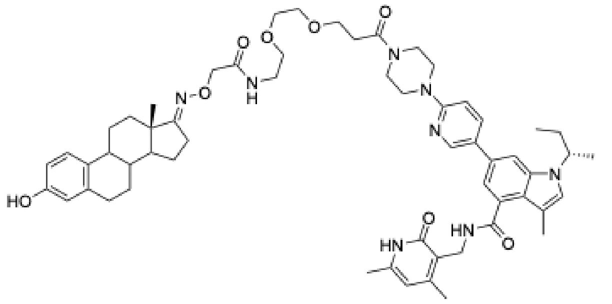

**$^{1}$H NMR** (400 MHz, DMSO-*D*$_6$) δ 11.43 (s, 1H), 8.96 (s, 1H), 8.46 (d, *J* = 2.5 Hz, 1H), 8.10 (d, *J* = 6.7 Hz, 1H), 7.94 (d, *J* = 8.7 Hz, 1H), 7.70 (s, 1H), 7.37 (t, *J* = 6.0 Hz, 1H), 7.22 (s, 1H), 7.14 (s, 1H), 6.99 (d, *J* = 8.5 Hz, 1H), 6.92 (d, *J* = 9.0 Hz, 1H), 6.46 (dd, *J* = 8.3, 2.7 Hz, 1H), 6.40 (d, *J* = 2.6 Hz, 1H), 5.83 (s, 1H), 4.56 (q, *J* = 6.8 Hz, 1H), 4.31 (d, *J* = 5.1 Hz, 2H), 4.27 (s, 2H), 3.51 (d, *J* = 19.7 Hz, 10H), 3.17 – 3.01 (m, 3H), 2.75 – 2.60 (m, 2H), 2.55 – 2.49 (m, 2H), 2.32 (d, *J* = 7.4 Hz, 2H), 2.29 (s, 3H), 2.20 (s, 3H), 2.12 (s, 4H), 2.07 (d, *J* = 3.9 Hz, 4H), 1.81 (q, *J* = 14.6 Hz, 6H), 1.53 – 1.22 (m, 10H), 0.84 (s, 4H), 0.69 (t, *J* = 7.3 Hz, 3H). (3 PEG-$CH_2$ and two piperazine-CH2 signals overlap with the $H_2O$ signal)
**HRMS** for $C_{58}H_{75}N_8O_8$ $[M+H]^+$ calc.: 1011.5702 Da; found: 1011.5676 Da
**Retention time Gradient A**: 2.484 min

4-(5-((furan-2-ylmethyl)amino)-[1,2,4]triazolo[4,3-*c*]pyrimidin-8-yl)-*N*-(1-((((13*S*,*E*)-3-hydroxy-13-methyl-6,7,8,9,11,12,13,14,15,16-decahydro-17*H*-cyclopenta[*a*] phenanthren-17-ylidene)amino)oxy)-2-oxo-6,9,12-trioxa-3-azatetradecan-14-yl)-*N*-methylbenzamide (**CS-1-132**):

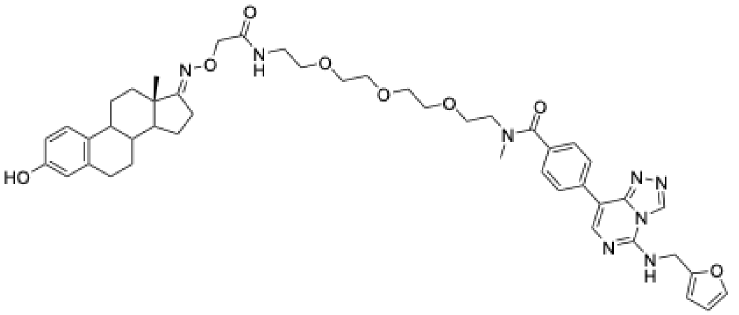

**$^1$H NMR** (500 MHz, DMSO) δ 9.48 (s, 1H), 8.97 (s, 1H), 8.27 (s, 1H), 8.18 (s, 2H), 8.11 (s, 1H), 7.63 (d, *J* = 1.7 Hz, 1H), 7.48 (d, *J* = 8.0 Hz, 2H), 7.38 (s, 1H), 7.01 (d, *J* = 8.5 Hz, 1H), 6.50 (dd, *J* = 8.4, 2.7 Hz, 2H), 6.44 (dt, *J* = 7.4, 3.0 Hz, 3H), 4.78 (d, *J* = 3.4 Hz, 2H), 4.32 (s, 2H), 3.63 (d, *J* = 15.9 Hz, 2H), 3.47 (d, *J* = 46.7 Hz, 10H), 3.26 (t, *J* = 5.9 Hz, 2H), 2.99 (s, 3H), 2.70 (dq, *J* = 16.0, 7.8 Hz, 2H), 2.32 – 2.21 (m, 1H), 2.11 (s, 1H), 1.93 – 1.72 (m, 4H), 1.59 – 1.43 (m, 1H), 1.45 – 1.20 (m, 6H), 0.85 (s, 3H). (one PEG-CH2 overlaps with $H_2O$ signal)

**HRMS** for $C_{46}H_{57}N_8O_8$ $[M+H]^+$ calc.: 849.4294 Da; found: 849.4291 Da

**Retention time Gradient A**: 2.541 min

*tert*-butyl (6-(4-((4-(pyridin-2-yl)thiazol-2-yl)amino)phenoxy)hexyl)carbamate (**CS-1-71**):

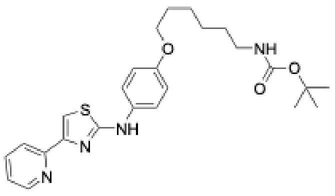

**CS-1-69**, *tert*-butyl (6-bromohexyl)carbamate (0.1 mmol each) and $K_2CO_3$ (0.6 mmol) were dissolved/suspended in MeCN (15 ml) and heated to reflux. Once TLC indicated full consumption of the starting material, the reaction was allowed to cool to r.t., precipitates were filtered off, the solvent was removed under reduced pressure and the crude was purified via flash chromatography (Hex/EtOAc, 0-80%) to obtain **CS-1-71** in 81% yield (38 mg). Before subsequent reactions, the compound was deprotected according to **General Procedure B** and used without further purification**.**

**$^1$H NMR** (400 MHz, CHLOROFORM-*D*) δ 8.49 (dd, *J* = 5.0, 1.6 Hz, 1H), 7.96 (s, 1H), 7.88 (d, *J* = 7.8 Hz, 1H), 7.60 (td, *J* = 7.7, 1.8 Hz, 1H), 7.30 – 7.22 (m, 3H), 7.08 (dd, *J* = 7.6, 4.8 Hz, 1H), 6.78 (d, *J* = 8.8 Hz, 2H), 3.83 (t, *J* = 6.4 Hz, 3H), 3.04 (d, *J* = 6.7 Hz, 2H), 1.67 (p, *J* = 6.5 Hz, 2H), 1.36 (s, 18H), 1.29 (d, *J* = 6.4 Hz, 3H).
**$^{13}$C NMR** (101 MHz, $CDCl_3$) δ 166.97, 156.13, 155.76, 152.75, 151.09, 149.38, 136.90, 133.80, 122.44, 121.79 (2C), 120.99, 115.37 (2C), 105.79, 79.14, 68.25, 40.60, 30.98, 29.26, 28.52 (3C), 26.62, 25.83.
**HRMS** for $C_{20}H_{24}N_4OS$ $[M+H]^+$ calc.: 469.2268 Da; found: 469.2266 Da
**HRMS** for $C_{20}H_{24}N_4OS$ $[M+H]^+$ calc.: 369.1744 Da; found: 369.1733 Da (after Boc deprotection)

*tert*-butyl (2-(2-(2-(2-(4-((4-(pyridin-2-yl)thiazol-2-yl)amino)phenoxy)ethoxy)ethoxy) ethoxy)ethyl)carbamate (**CS-1-76**):

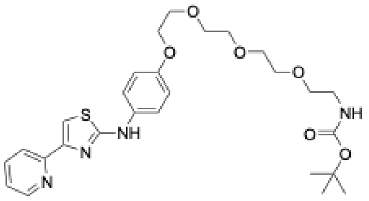

**CS-1-69**, *tert*-butyl (2-(2-(2-(2-bromoethoxy)ethoxy)ethoxy)ethyl) carbamate (0.1 mmol each) and $K_2CO_3$ (0.6 mmol) were dissolved/suspended in MeCN (15 ml) and heated to reflux. Once TLC indicated full consumption of the starting material, the reaction was allowed to cool to r.t., precipitates were filtered off, the solvent was removed under reduced pressure and the crude was purified via flash chromatography (Hex/EtOAc, 30-100%) to obtain **CS-1-76** in 77% yield (42 mg). Before subsequent reactions, the compound was deprotected according to **General Procedure B** and used without further purification.

**$^1$H NMR** (400 MHz, $CDCl_3$) δ 8.58 (d, *J* = 4.8 Hz, 1H), 7.95 (d, *J* = 7.8 Hz, 1H), 7.71 (t, *J* = 7.7 Hz, 1H), 7.54 (s, 1H), 7.34 – 7.32 (m, 3H), 7.21 – 7.14 (m, 1H), 6.91 (d, *J* = 8.9 Hz, 2H), 4.15 – 4.08 (m, 2H), 3.85 (t, *J* = 4.9 Hz, 2H), 3.76 – 3.50 (m, 10H), 3.30 (d, *J* = 5.3 Hz, 2H), 1.43 (s, 9H).
**$^{13}$C NMR** (101 MHz, $CDCl_3$) δ 166.67, 156.10, 155.52, 152.75, 151.21, 149.44, 136.91, 133.98, 122.48, 121.71, 120.96, 115.63, 105.91, 79.28, 70.93, 70.74, 70.67, 70.34, 70.30, 69.87, 67.90, 40.46, 28.52.
**HRMS** for $C_{27}H_{36}N_4O_6SNa^+$ $[M+Na]^+$ calc.: 567.2248 Da; found: 567.2236 Da

*N*-(6-aminohexyl)-3-((4-(pyridin-2-yl)thiazol-2-yl)amino)benzamide (**CS-1-99**):

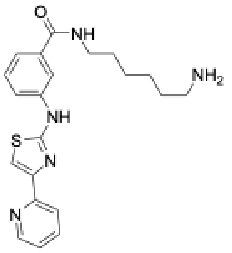

Was prepared according to **General Procedure A** starting from 3-((4-(pyridin-2-yl)thiazol-2-yl)amino)benzoic acid and *tert*-butyl (6-aminohexyl)carbamate. After purification of the Boc-protected intermediate, deprotection according to **General Procedure B** afforded **CS-1-99** as a yellow solid that was used without further purification. (115 mg, 89%)

**$^1$H NMR** (500 MHz, DMSO-*D*$_6$) δ 10.60 (s, 1H), 8.62 – 8.57 (m, 1H), 8.48 (t, *J* = 5.7 Hz, 1H), 8.43 (s, 1H), 8.24 – 8.18 (m, 1H), 8.03 (d, *J* = 7.8 Hz, 1H), 7.96 (dt, *J* = 7.0, 2.3 Hz, 1H), 7.89 (td, *J* = 7.7, 1.8 Hz, 1H), 7.58 (s, 1H), 7.48 – 7.38 (m, 2H), 7.33 (ddd, *J* = 7.6, 4.7, 1.3 Hz, 1H), 3.27 (q, *J* = 6.6 Hz, 2H), 2.75 (t, *J* = 7.5 Hz, 2H), 1.53 (tt, *J* = 11.0, 4.8 Hz, 4H), 1.42 – 1.30 (m, 4H).
**$^{13}$C NMR** (126 MHz, DMSO-*D*$_6$) δ 166.85, 163.72, 152.58, 150.79, 149.90, 141.67, 137.68, 136.30, 129.38, 123.16, 120.83, 120.04, 116.60, 107.58, 39.13, 29.44, 27.76, 26.46, 26.07. (one carbon overlaps with solvent peak)
**HRMS** for $C_{21}H_{26}N_5OS$ [M+H]$^+$ calc.: 396.1853 Da; found: 396.1844 Da

*N*-(2-(2-(2-aminoethoxy)ethoxy)ethyl)-3-((4-(pyridin-2-yl)thiazol-2-yl)amino)benzamide (**CS-1-138**):

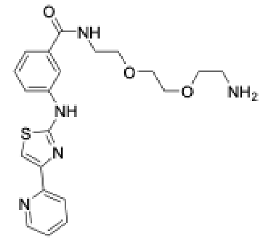

Was prepared according to **General Procedure A** starting from 3-((4-(pyridin-2-yl)thiazol-2-yl)amino)benzoic acid and *tert*-butyl (6-aminohexyl)carbamate. After purification of the Boc-protected intermediate, deprotection according to **General Procedure B** afforded **CS-1-138** as a yellow solid that was used without further purification. (145 mg, 86%)

**$^1$H NMR** (400 MHz, DMSO-*D*$_6$) δ 11.06 (s, 1H), 8.77 (d, *J* = 5.2 Hz, 1H), 8.66 (t, *J* = 5.7 Hz, 1H), 8.46 (s, 1H), 8.33 (s, 1H), 8.13 (d, *J* = 7.8 Hz, 2H), 8.08 (s, 1H), 7.82 (t, *J* = 5.4 Hz, 1H), 7.46 (dt, *J* = 15.3, 7.7 Hz, 2H), 3.56 (s, 6H), 3.44 (q, *J* = 5.6 Hz, 2H), 3.10 – 2.97 (m, 2H), 2.91 (q, *J* = 5.6 Hz, 2H).
**$^{13}$C NMR** (101 MHz, DMSO-*D*$_6$) δ 166.99, 164.51, 146.79, 145.21, 143.81, 141.20, 135.97, 129.62, 125.18, 120.83, 120.44, 116.80, 114.52, 106.23, 70.20, 69.95, 69.41, 67.13, 45.84, 38.99.
**HRMS** for $C_{21}H_{25}N_5O_3S$ [M+H]$^+$ calc.: 428.1751 Da; found: 428.1741 Da

*N*-(3-aminopropyl)-3-((4-(pyridin-2-yl)thiazol-2-yl)amino)benzamide (**CS-1-157**):

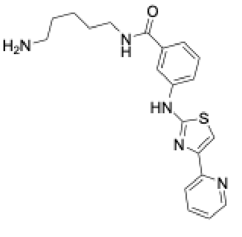

Was prepared according to **General Procedure A** starting from 3-((4-(pyridin-2-yl)thiazol-2-yl)amino)benzoic acid and *tert*-butyl (5-pentyl)carbamate. After purification of the Boc-protected intermediate, deprotection according to **General Procedure B** afforded **CS-1-159** as a yellow solid that was used without further purification. (168 mg, 88%) A small portion was purified via semipreparative HPLC for analytical purposes.

**$^1$H NMR** (500 MHz, DMSO-*D*$_6$) δ 10.57 (s, 1H), 8.63 – 8.58 (m, 1H), 8.47 (t, *J* = 5.6 Hz, 1H), 8.35 (s, 1H), 8.24 – 8.17 (m, 1H), 8.04 (d, *J* = 7.8 Hz, 1H), 7.96 (dt, *J* = 6.8, 2.4 Hz, 1H), 7.90 (td, *J* = 7.7, 1.8 Hz, 1H), 7.59

(s, 1H), 7.49 – 7.40 (m, 2H), 7.34 (ddd, $J$ = 7.5, 4.7, 1.2 Hz, 1H), 3.28 (q, $J$ = 6.6 Hz, 2H), 2.78 (s, 2H), 1.58 (dp, $J$ = 10.7, 7.4 Hz, 4H), 1.38 (qd, $J$ = 8.7, 6.1 Hz, 2H).
**HRMS** for $C_{20}H_{23}N_5OS$ $[M+H]^+$ calc.: 382.1696 Da; found: 382.1690 Da
**Retention time Gradient A**: 1.623 min

piperazin-1-yl(3-((4-(pyridin-2-yl)thiazol-2-yl)amino)phenyl)methanone (**CS-1-158**):

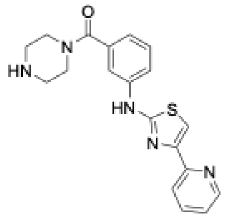


Was prepared according to **General Procedure A** starting from 3-((4-(pyridin-2-yl)thiazol-2-yl)amino)benzoic acid and *tert*-butyl piperazine-1-carboxylate. After purification of the Boc-protected intermediate, deprotection according to **General Procedure B** afforded **CS-1-158** as a yellow solid that was used without further purification. (114 mg, 62%) A small portion was purified via semipreparative HPLC for analytical purposes.

**$^1$H NMR** (500 MHz, DMSO-$D_6$) δ 10.52 (s, 1H), 8.92 (s, 2H), 8.62 (dd, $J$ = 4.8, 1.6 Hz, 1H), 8.00 (d, $J$ = 7.8 Hz, 1H), 7.95 (td, $J$ = 7.7, 1.8 Hz, 1H), 7.89 (dd, $J$ = 8.2, 2.3 Hz, 1H), 7.79 (d, $J$ = 2.0 Hz, 1H), 7.63 (s, 1H), 7.47 (t, $J$ = 7.9 Hz, 1H), 7.41 – 7.35 (m, 1H), 7.08 (d, $J$ = 7.5 Hz, 1H), 3.19 (s, 4H). Proton signal for the piperazine $CH_2$-groups closer to the amide overlaps with $H_2O$ signal.
**HRMS** for $C_{19}H_{19}N_5OS$ $[M+H]^+$ calc.: 366.1383 Da; found: 366.1376 Da
**Retention time Gradient A**: 1.539 min

*N*-(12-aminododecyl)-3-((4-(pyridin-2-yl)thiazol-2-yl)amino)benzamide (**CS-1-159**):

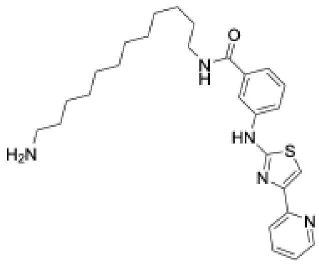


Was prepared according to **General Procedure A** starting from 3-((4-(pyridin-2-yl)thiazol-2-yl)amino)benzoic acid and *tert*-butyl (12-dodecyl)carbamate. After purification of the Boc-protected intermediate, deprotection according to **General Procedure B** afforded **CS-1-159** as a yellow solid that was used without further purification. (191 mg, 83%)

**$^1$H NMR** (400 MHz, TFA/$CDCl_3$/DMSO-$D_6$) δ 10.47 (s, 1H), 8.62 (d, $J$ = 5.1 Hz, 1H), 8.31 (t, $J$ = 5.6 Hz, 1H), 8.20 (p, $J$ = 2.7 Hz, 1H), 8.11 – 8.04 (m, 2H), 7.94 (dt, $J$ = 7.9, 1.8 Hz, 1H), 7.75 (s, 1H), 7.52 – 7.42 (m, 1H), 7.41 – 7.31 (m, 2H), 3.23 (q, $J$ = 6.7 Hz, 2H), 2.71 (td, $J$ = 7.7, 5.5 Hz, 2H), 1.49 (q, $J$ = 7.4 Hz, 4H), 1.20 (d, $J$ = 6.5 Hz, 16H).
**HRMS** for $C_{27}H_{37}N_5OS$ $[M+H]^+$ calc.: 480.2792 Da; found: 480.2783 Da
**Retention time Gradient A**: 2.031 min

*N*-(3-aminopropyl)-3-((4-(pyridin-2-yl)thiazol-2-yl)amino)benzamide (**CS-1-160**):

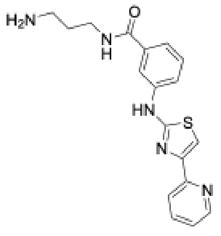

Was prepared according to **General Procedure A** starting from 3-((4-(pyridin-2-yl)thiazol-2-yl)amino)benzoic acid and *tert*-butyl (3-aminopropyl)carbamate. After purification of the Boc-protected intermediate, was deprotection according to **General Procedure B** afforded **CS-1-160** as a yellow solid that was used without further purification. (127 mg, 89%) A small portion was purified via semipreparative HPLC for analytical purposes.

**$^1$H NMR** (400 MHz, DMSO-$D_6$) δ 10.49 (s, 1H), 8.61 (t, *J* = 5.8 Hz, 1H), 8.59 – 8.52 (m, 1H), 8.23 (s, 1H), 8.19 (d, *J* = 2.4 Hz, 1H), 7.99 (d, *J* = 7.9 Hz, 1H), 7.89 (ddt, *J* = 17.4, 9.6, 5.3 Hz, 2H), 7.54 (s, 1H), 7.46 – 7.36 (m, 2H), 7.33 – 7.24 (m, 1H), 3.32 (q, *J* = 6.4 Hz, 2H), 2.82 (t, *J* = 7.4 Hz, 2H), 1.79 (p, *J* = 7.0 Hz, 2H).
**HRMS** for $C_{18}H_{19}N_5OS$ $[M+H]^+$ calc.: 354.1383 Da; found: 354.1377 Da
**Retention time Gradient A**: 1.579 min

(3-((4-(pyridin-2-yl)thiazol-2-yl)amino)phenyl)(3,9-diazaspiro[5.5]undecan-3-yl) methanone (**CS-1-161**):

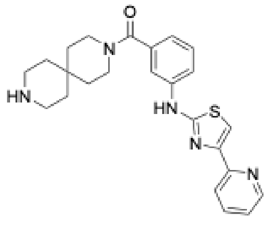

Was prepared according to **General Procedure A** starting from 3-((4-(pyridin-2-yl)thiazol-2-yl)amino)benzoic acid and *tert*-butyl (6-aminopropyl)carbamate. After purification of the Boc-protected intermediate, was deprotection according to **General Procedure B** afforded **CS-1-157** as a yellow solid that was used without further purification. (76 mg, 83%)

**$^1$H NMR** (400 MHz, DMSO-$D_6$) δ 10.54 (s, 1H), 8.64 – 8.47 (m, 1H), 8.35 (s, 1H), 7.92 (d, *J* = 7.8 Hz, 1H), 7.85 (td, *J* = 7.6, 1.8 Hz, 1H), 7.76 (d, *J* = 7.6 Hz, 2H), 7.54 (s, 1H), 7.38 (t, *J* = 7.8 Hz, 1H), 7.28 (ddd, *J* = 7.4, 4.7, 1.3 Hz, 1H), 6.92 (d, *J* = 7.5 Hz, 1H), 3.30 (s, 4H), 2.95 (s, 4H), 1.72 – 1.31 (m, 8H).
**HRMS** for $C_{24}H_{27}N_5OS$ $[M+H]^+$ calc.: 434.2009 Da; found: 434.2004 Da
**Retention time Gradient A**: 1.620 min

4-(3-(4-cyano-3-(trifluoromethyl)phenyl)-5,5-dimethyl-2,4-dioxoimidazolidin-1-yl)butanoic acid (**CS-1-154**):

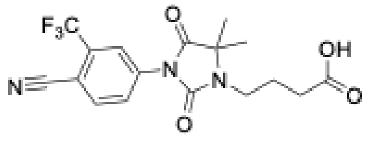

CS-1-54 was prepared according to a literature procedure reported by Gustafson et al.[9] After oxidation was complete, the reaction was diluted with 1 M HCl and extracted with $Et_2O$. Combined organic layers were dried over $MgSO_4$ filtered and evaporated. The compound obtained after lyophilization from water/MeCN as a white powder and used without further purification.

**$^1$H NMR** (400 MHz, METHANOL-$D_4$) δ 8.17 (d, *J* = 2.1 Hz, 1H), 8.11 – 7.98 (m, 2H), 3.43 (dd, *J* = 8.6, 6.4 Hz, 2H), 2.40 (t, *J* = 7.2 Hz, 2H), 2.06 – 1.91 (m, 2H), 1.56 – 1.47 (m, 6H).
**HRMS** for $C_{17}H_{17}F_3N_3O_4$ $[M+H]^+$ calc.: 384.1166 Da; found: 384.1159 Da

4-(3-(4-cyano-3-(trifluoromethyl)phenyl)-5,5-dimethyl-4-oxo-2-thioxoimidazolidin-1-yl) butanoic acid (**CS-1-170**)

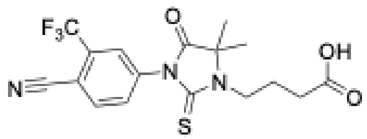

**CS-1-170** was prepared analogously to **CS-1-154** and obtained as an amorphous solid.

**$^1$H NMR** (400 MHz, METHANOL-$D_4$) δ 8.14 – 7.98 (m, 2H), 7.87 (dd, $J$ = 8.3, 2.2 Hz, 1H), 3.85 – 3.71 (m, 2H), 2.40 (t, $J$ = 7.2 Hz, 2H), 2.20 – 2.02 (m, 1H), 1.56 (s, 6H).
**HRMS** for $C_{17}H_{17}F_3N_3O_3S$ $[M+H]^+$ calc.: 400.0937 Da; found: 400.0927 Da

**CS-1-69:**

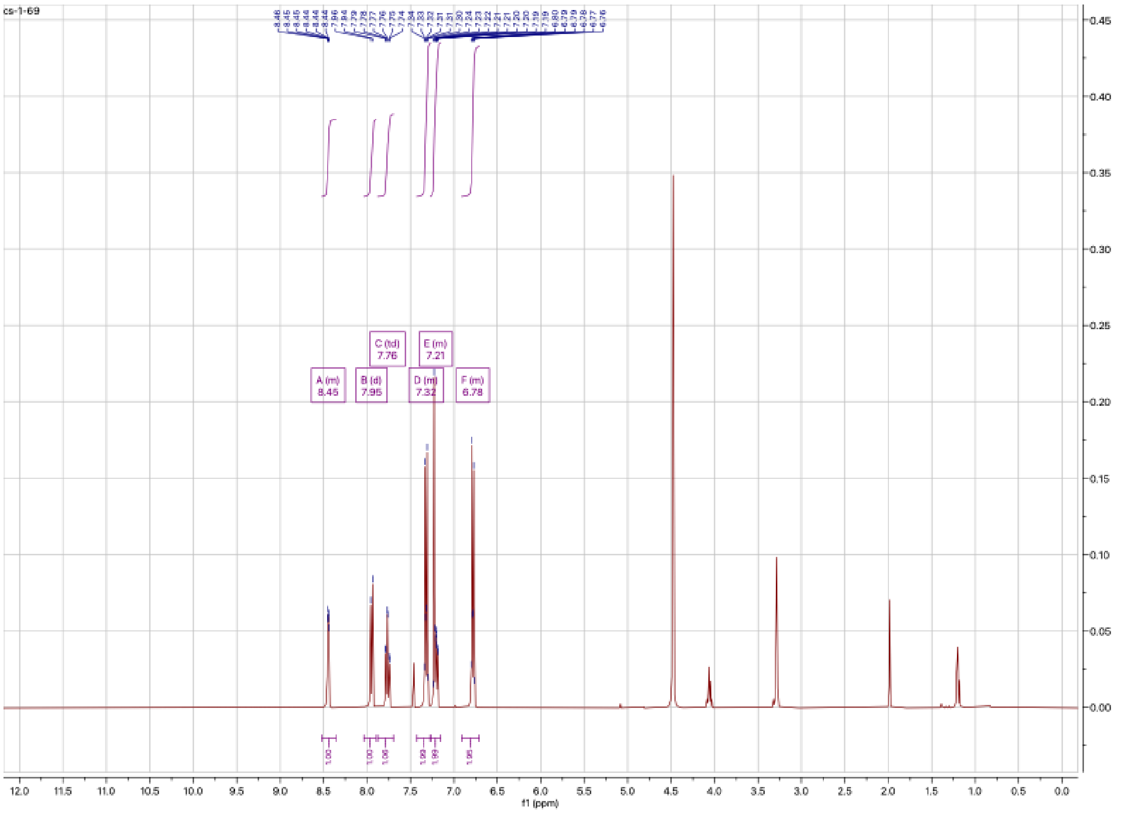
cs-1-69
A (m) 8.45
B (d) 7.95
C (td) 7.76
D (m) 7.32
E (m) 7.21
F (m) 6.78
f1 (ppm)

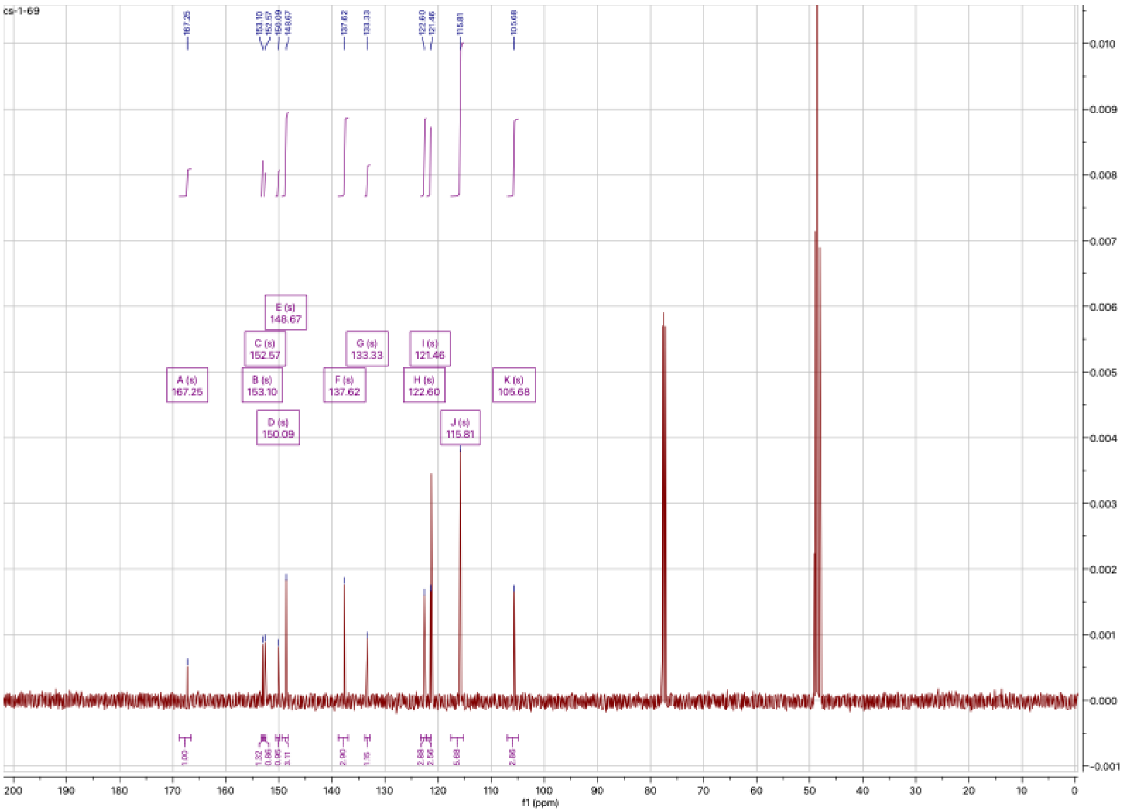
cs-1-69
A (s) 167.25
B (s) 153.10
C (s) 152.57
D (s) 150.09
E (s) 148.67
F (s) 137.62
G (s) 133.33
H (s) 122.60
I (s) 121.46
J (s) 115.81
K (s) 105.68
f1 (ppm)

**CS-1-71:**

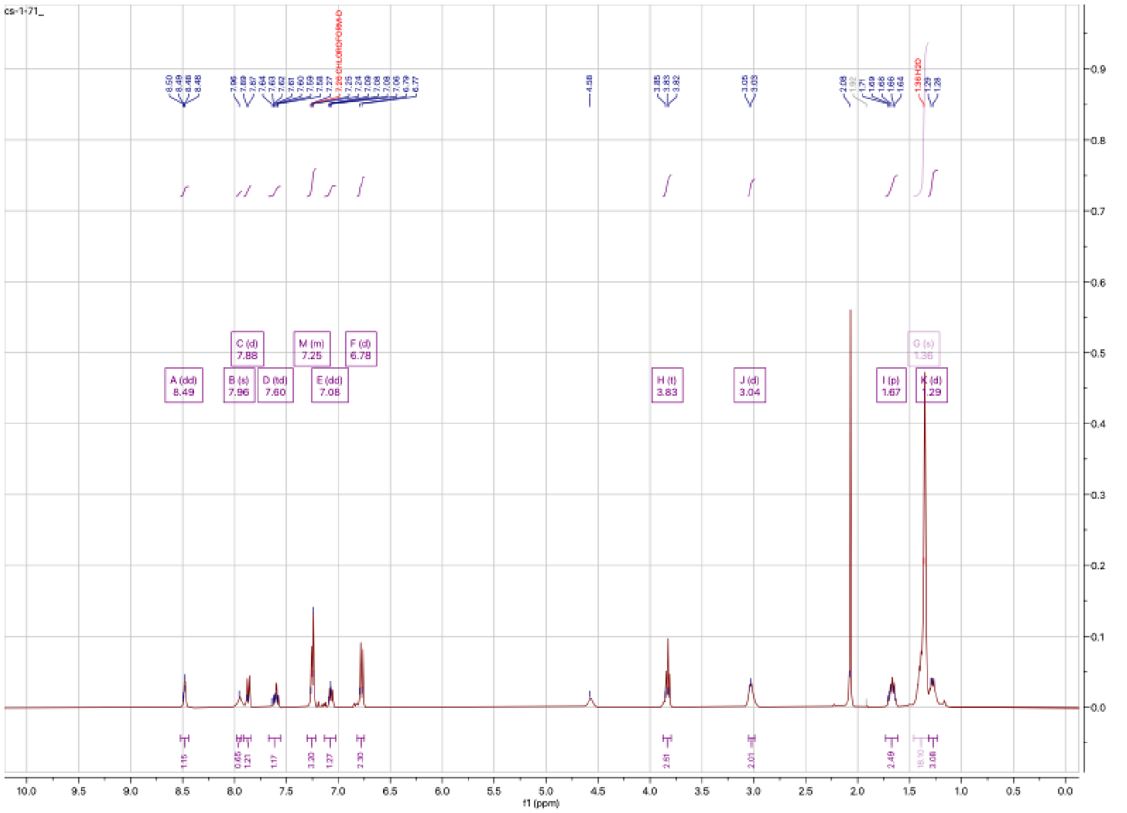

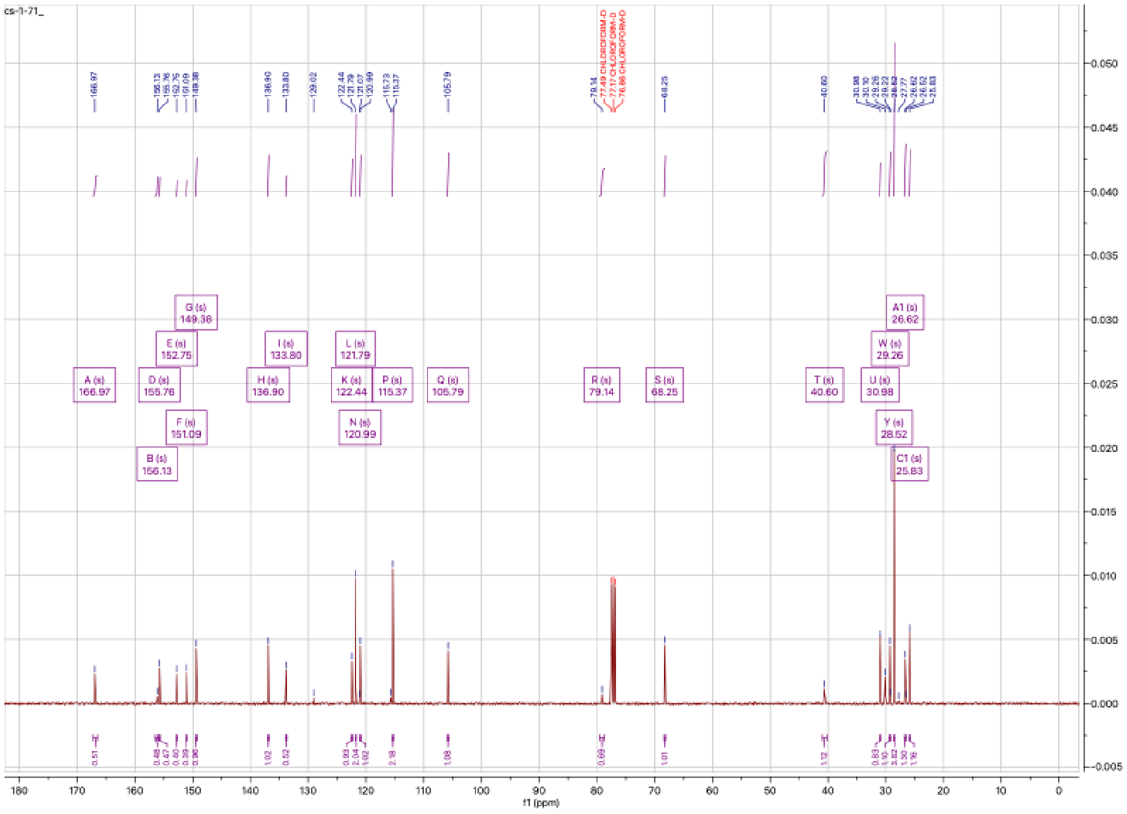

# CS-1-76:

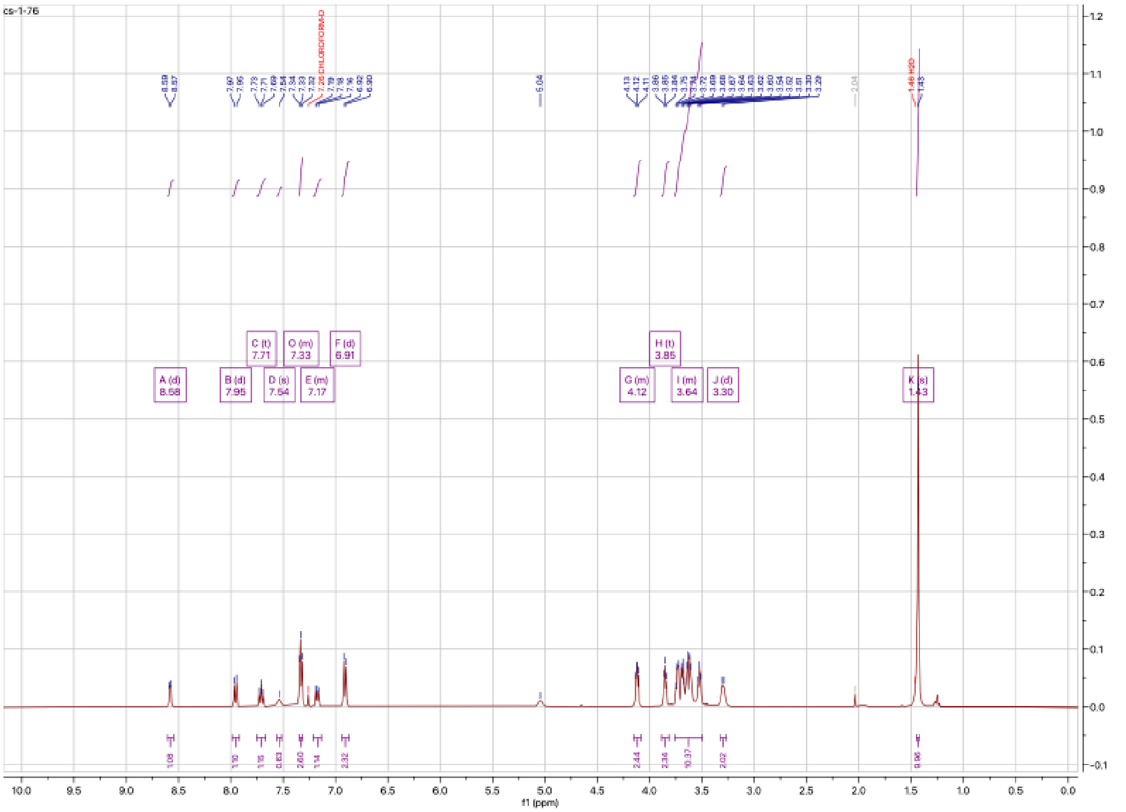

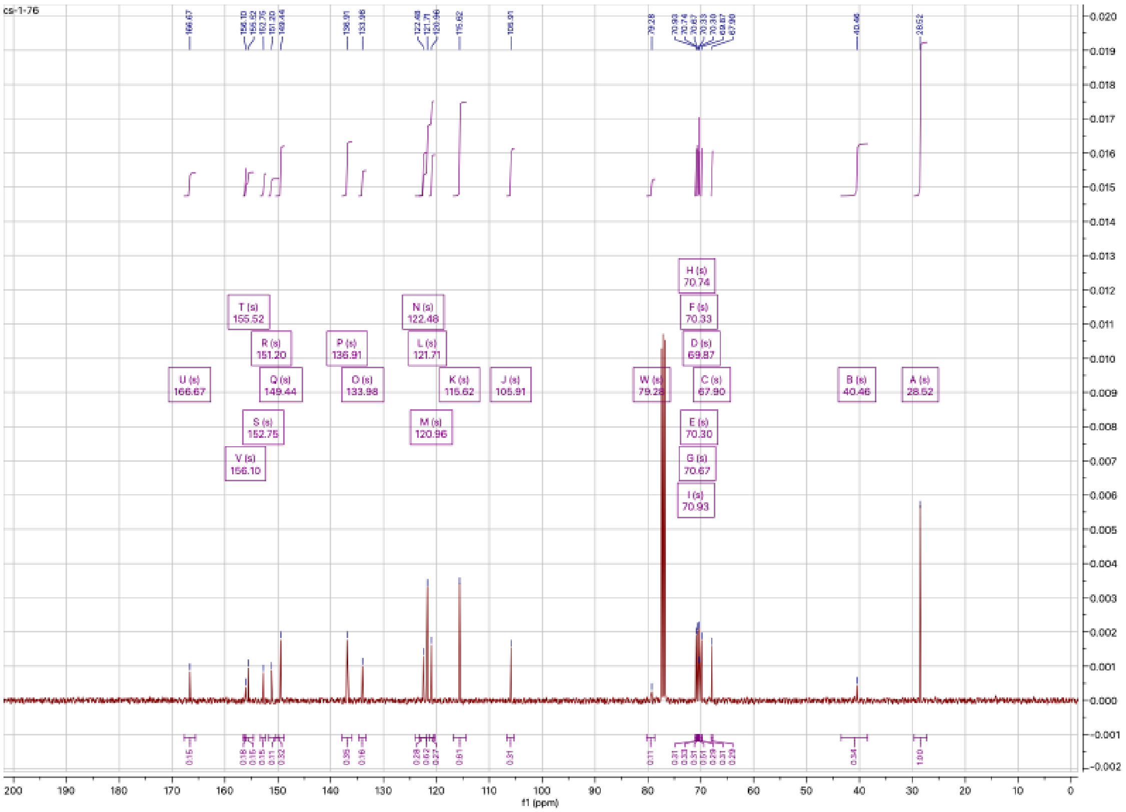

# CS-1-85:

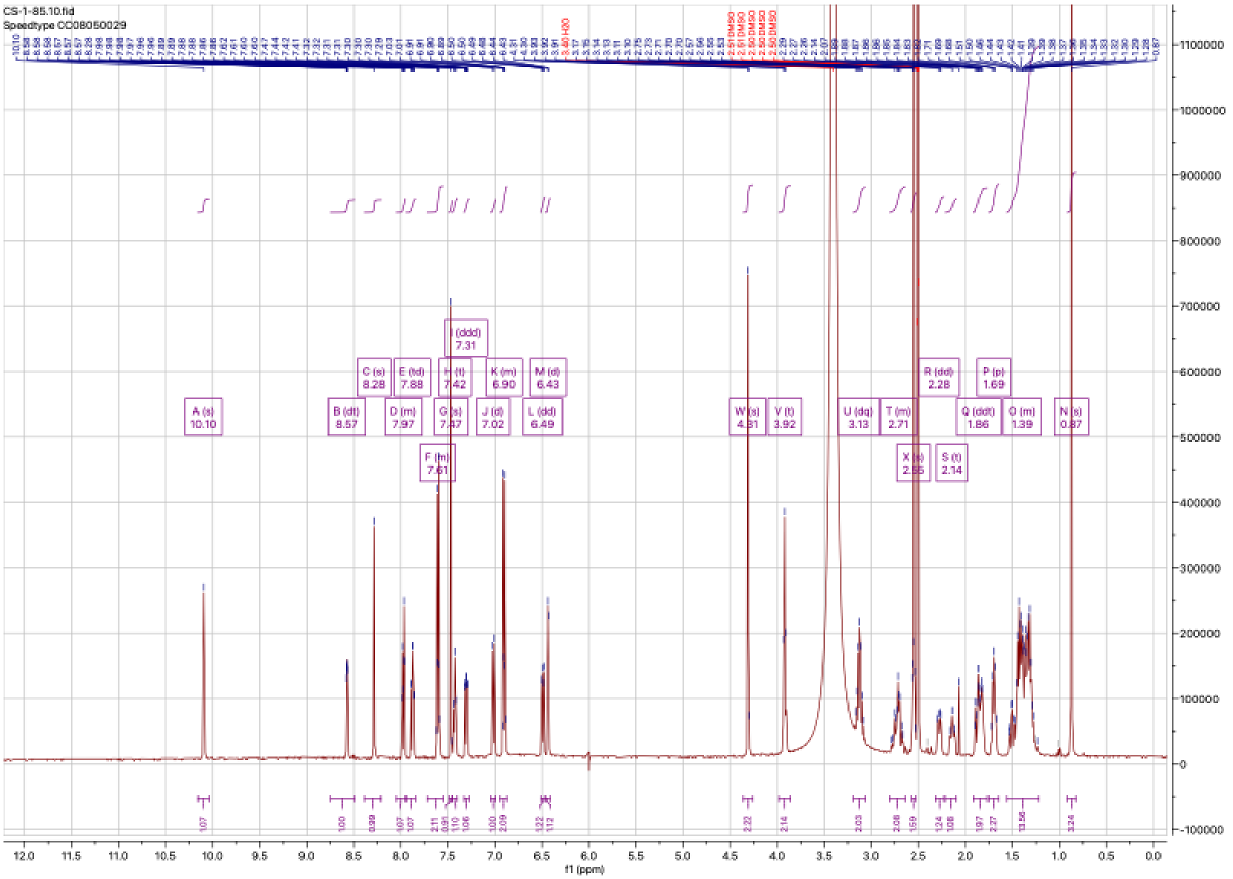
CS-1-85.10.fid
Speedtype CC08050029
A (s) 10.10
B (dt) 8.57
C (s) 8.28
D (m) 7.97
E (td) 7.88
F (m) 7.61
G (s) 7.47
H (t) 7.42
I (ddd) 7.31
J (d) 7.02
K (m) 6.90
L (dd) 6.49
M (d) 6.43
W (s) 4.31
V (t) 3.92
U (dq) 3.13
T (m) 2.71
X (s) 2.55
R (dd) 2.28
S (t) 2.14
Q (ddt) 1.86
P (p) 1.69
O (m) 1.39
N (s) 0.87
f1 (ppm)

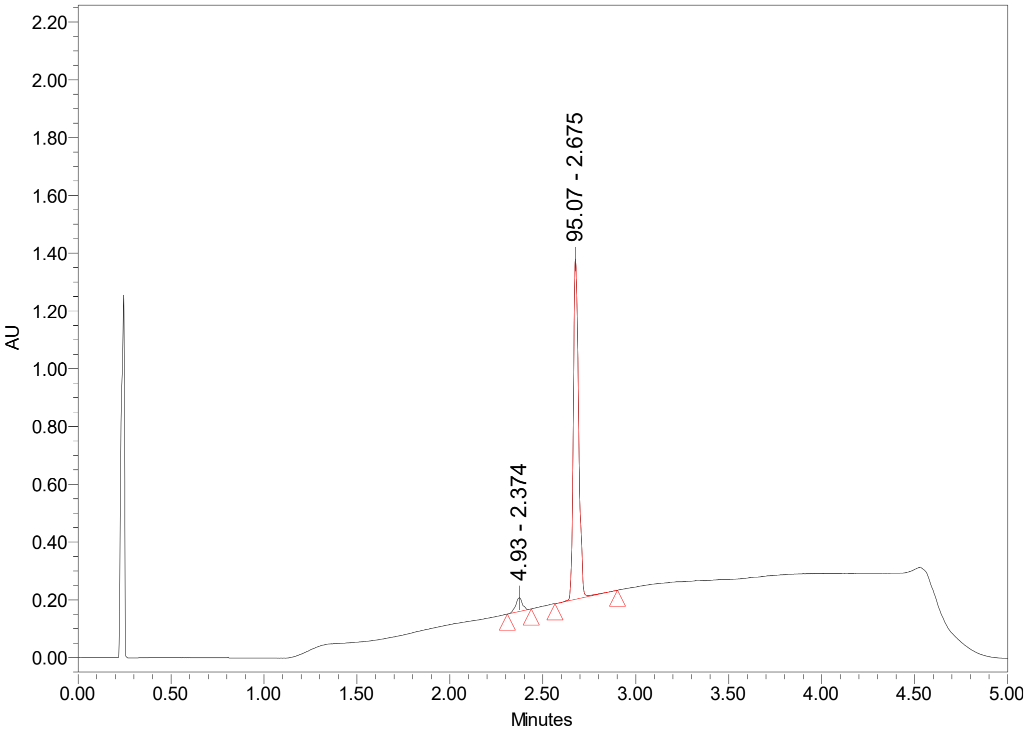
AU
Minutes
4.93 - 2.374
95.07 - 2.675

## CS-1-86:

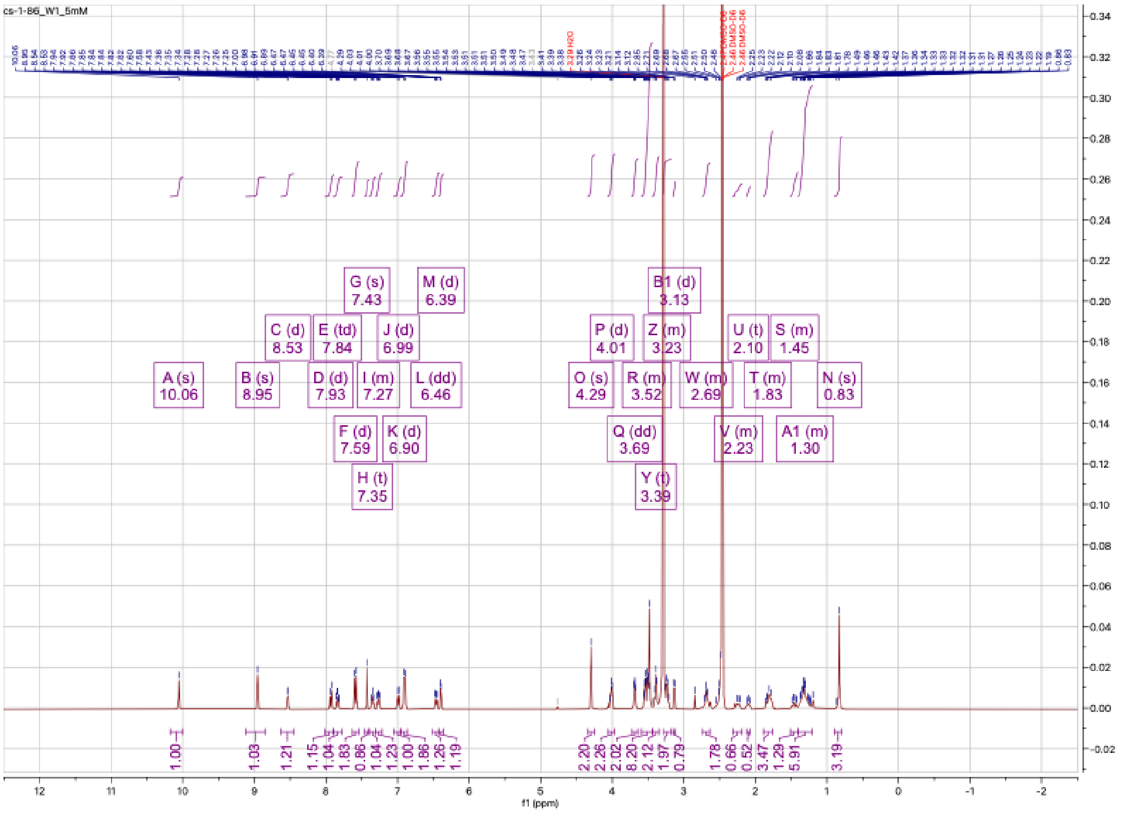

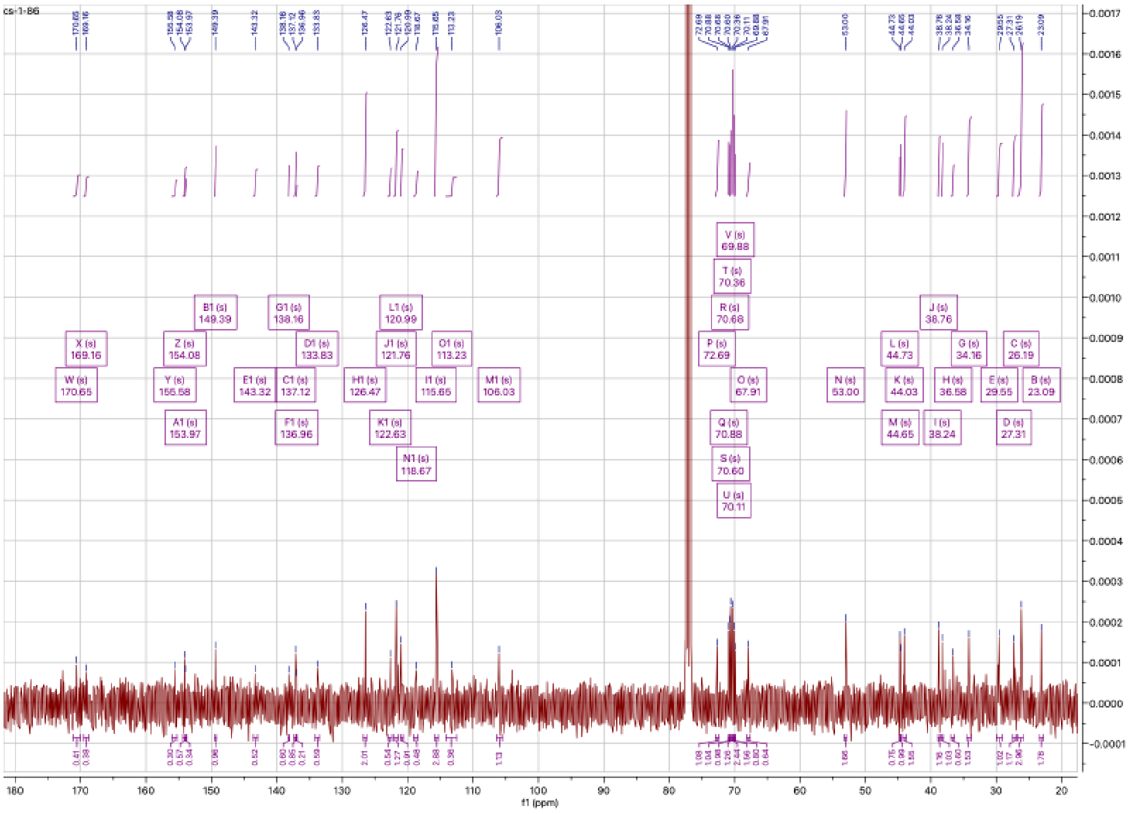

**CS-1-99:**

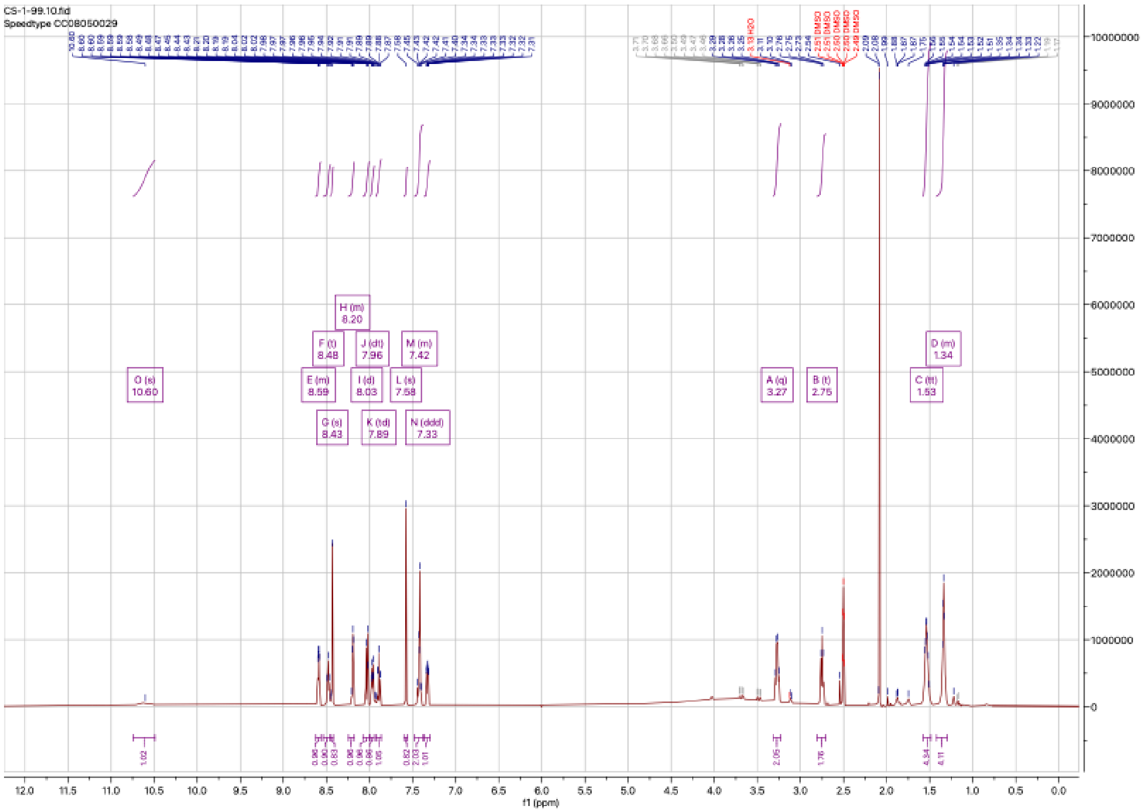

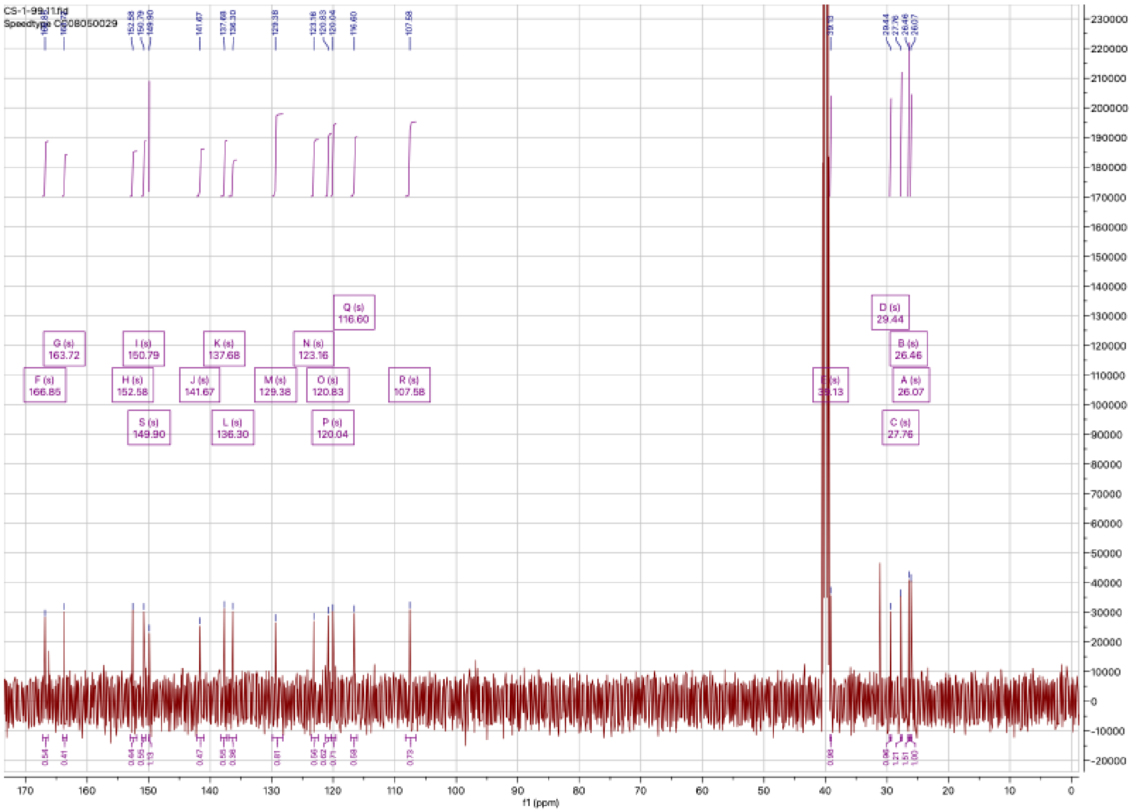

**CS-1-102**:

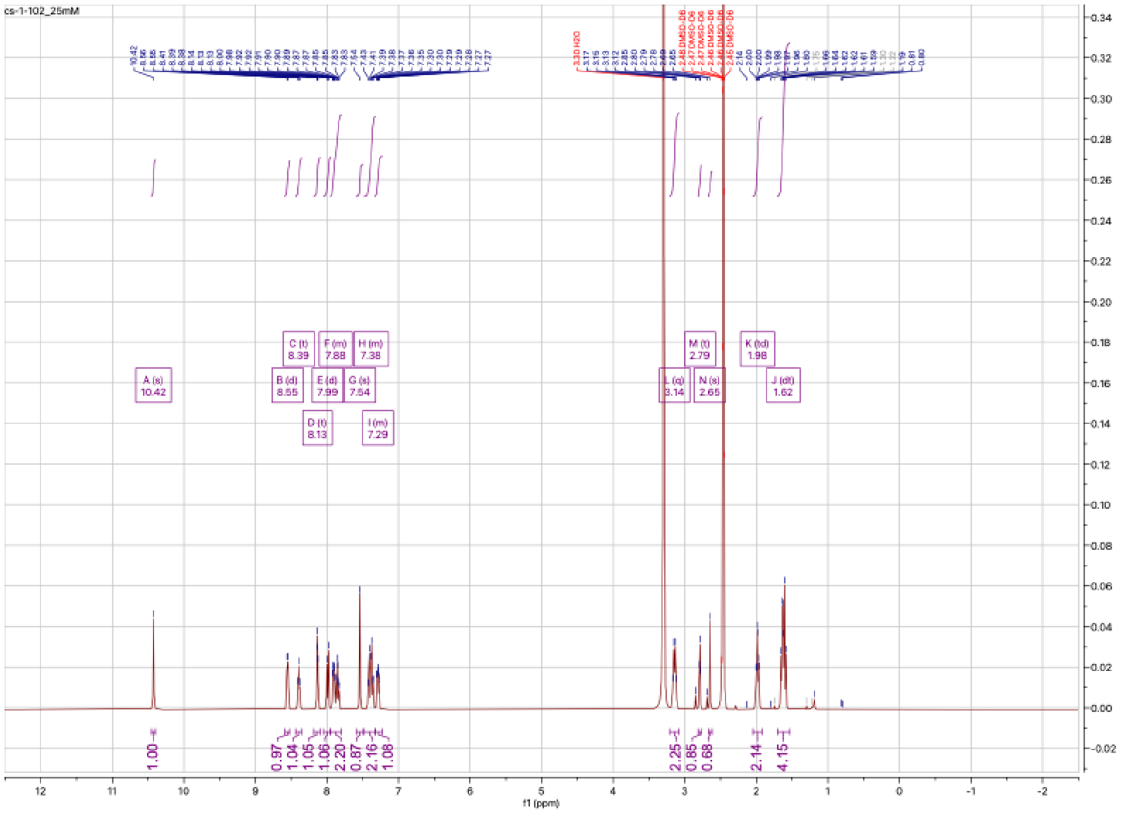

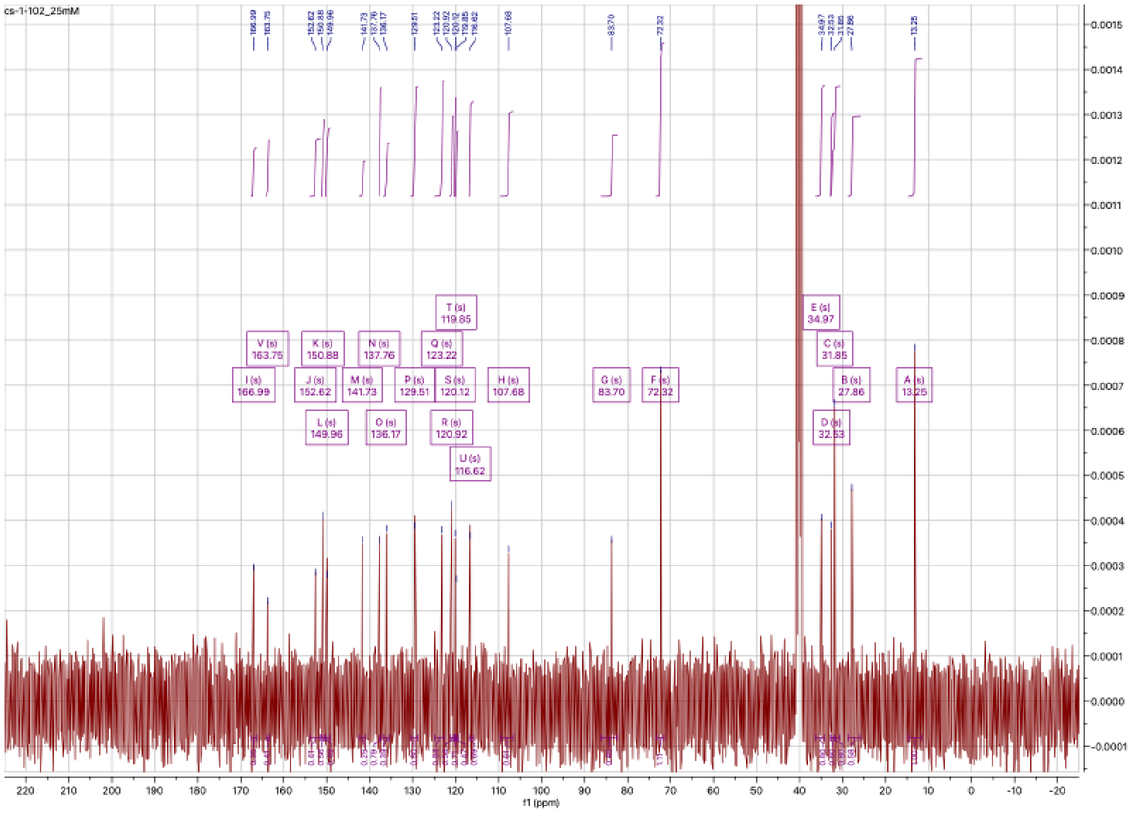

**CS-1-103**:

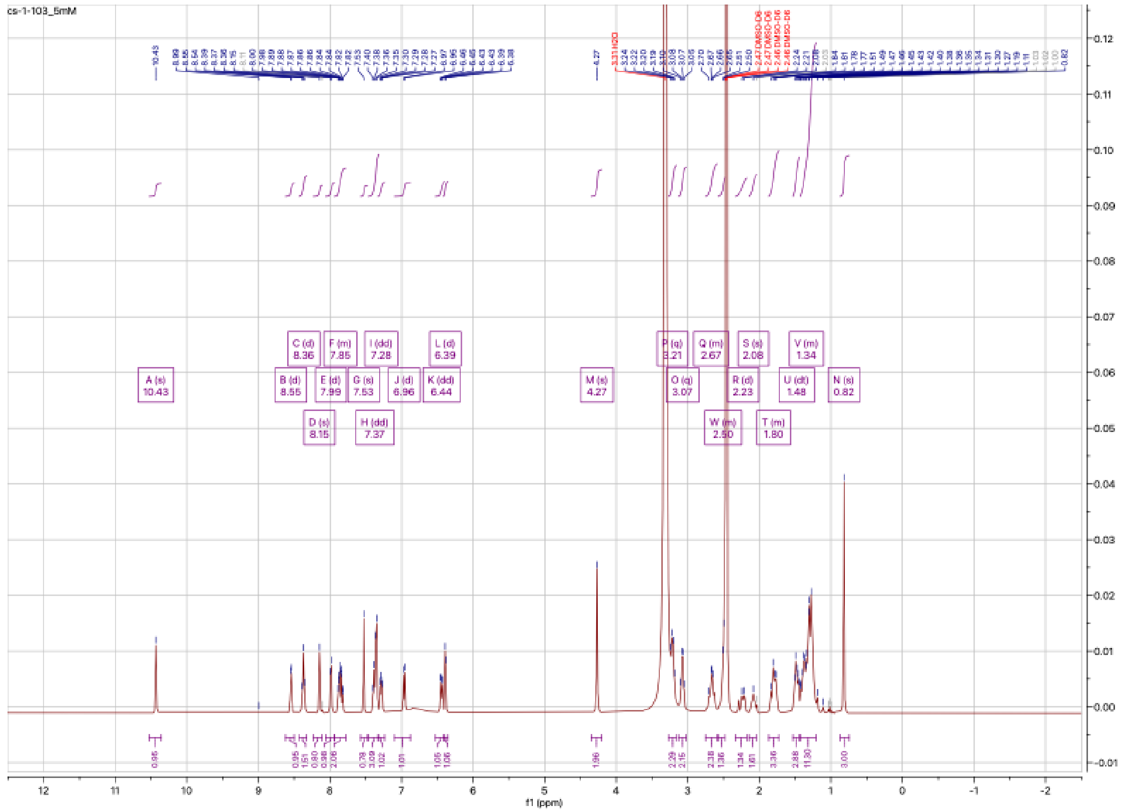

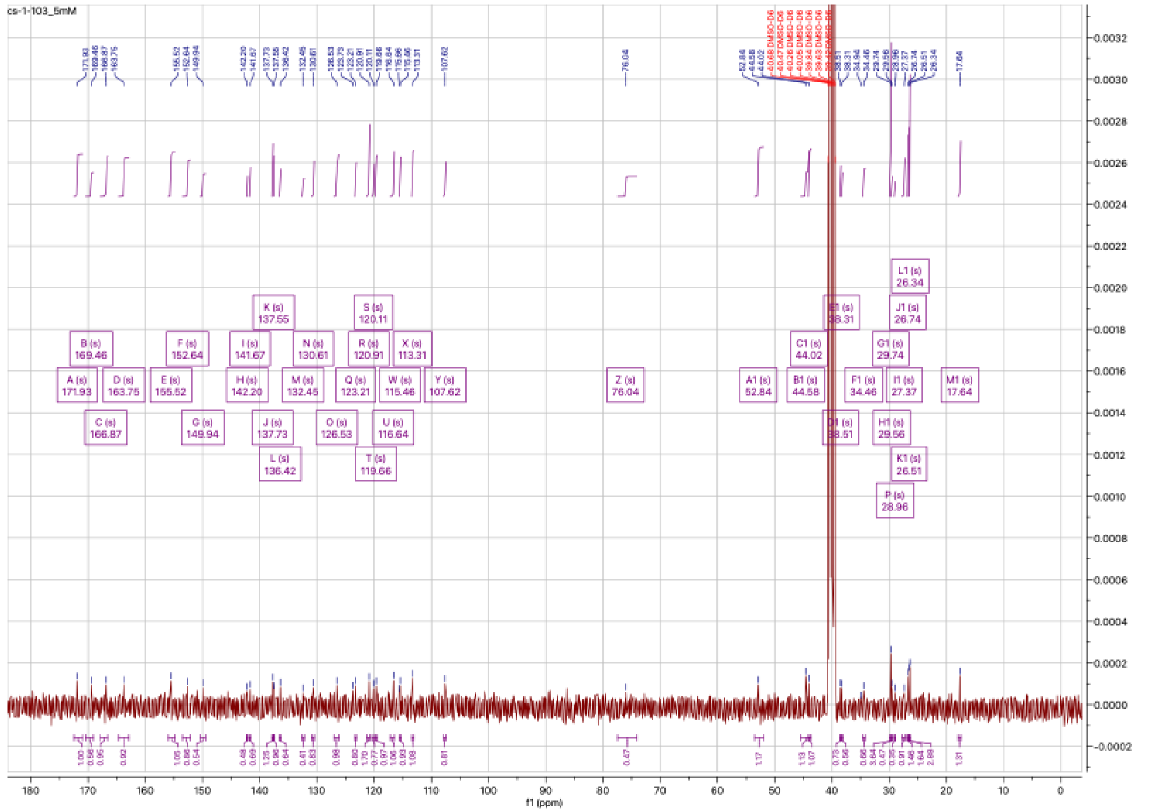

**CS-1-123:**

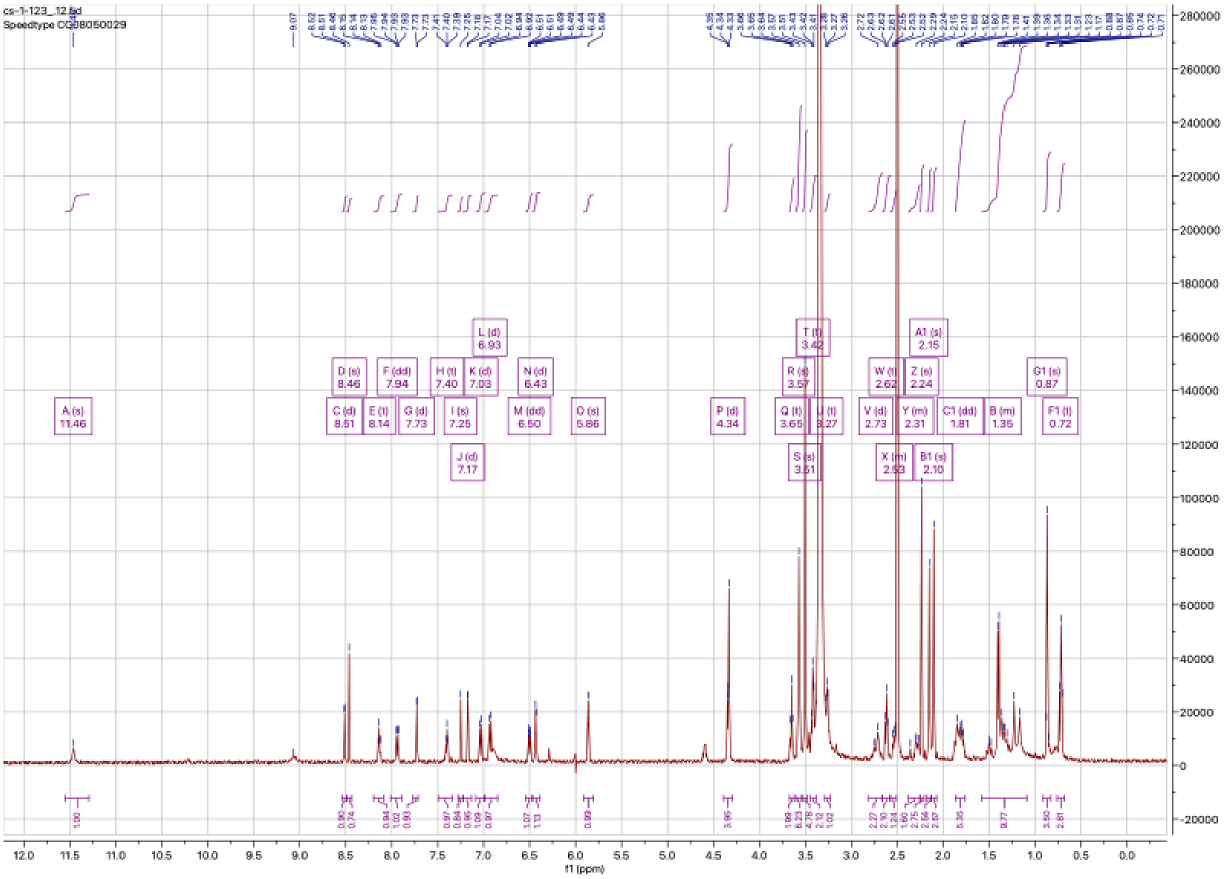

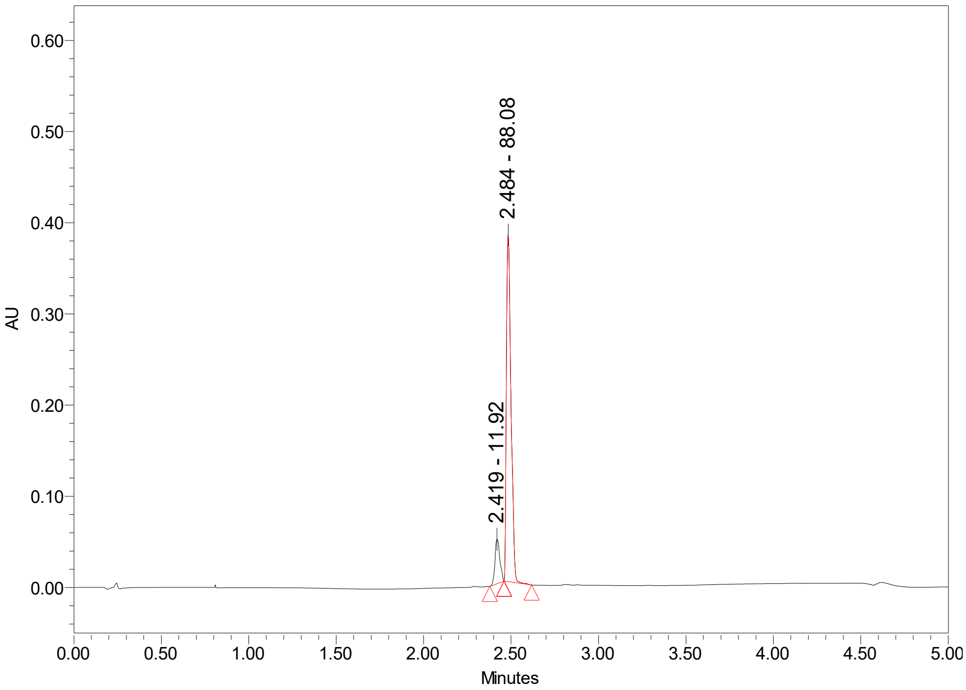

**CS-1-132:**

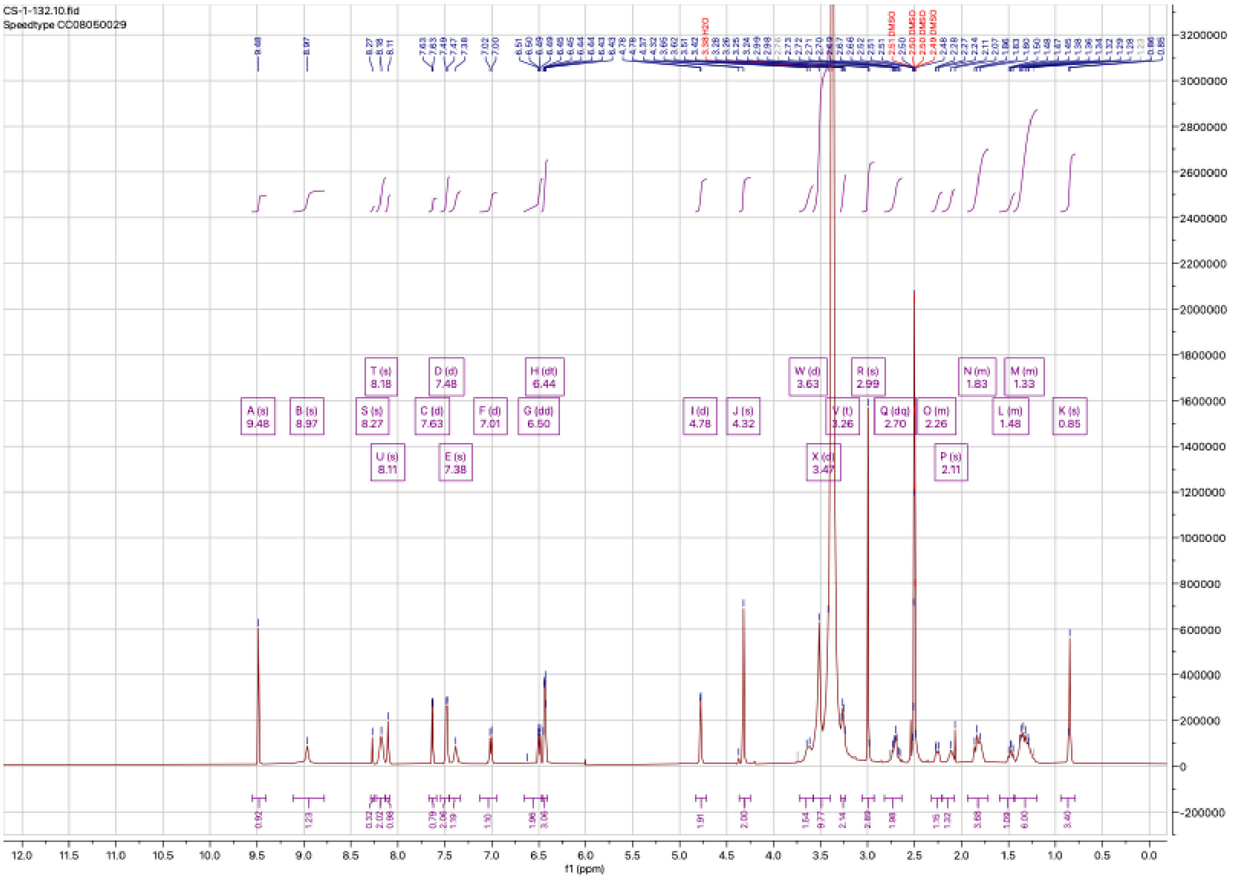


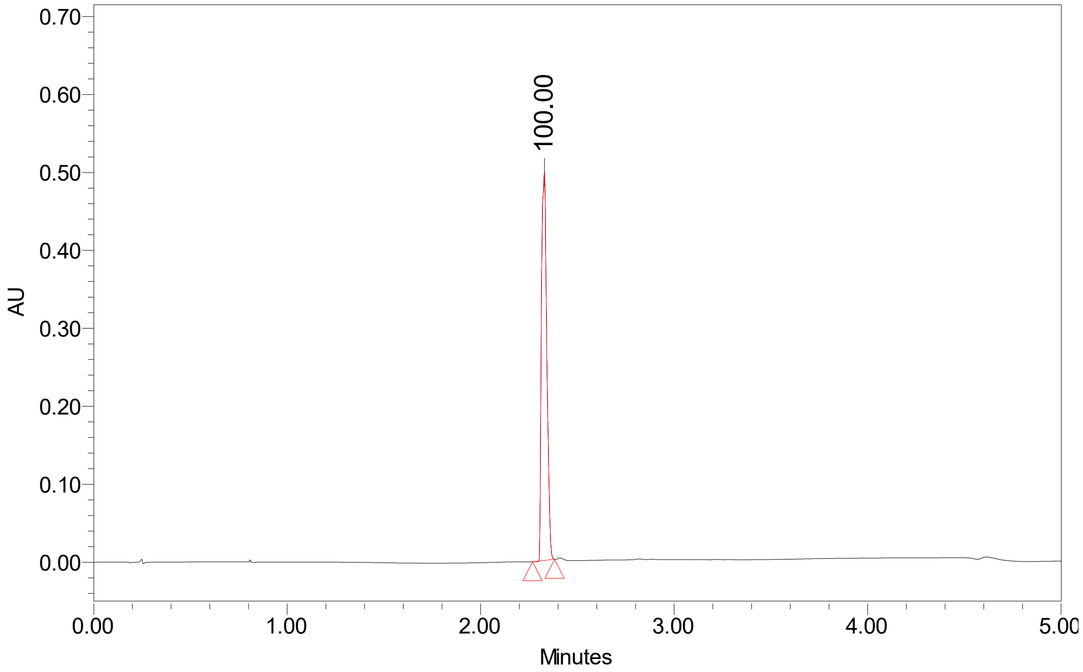

**CS-1-157:**

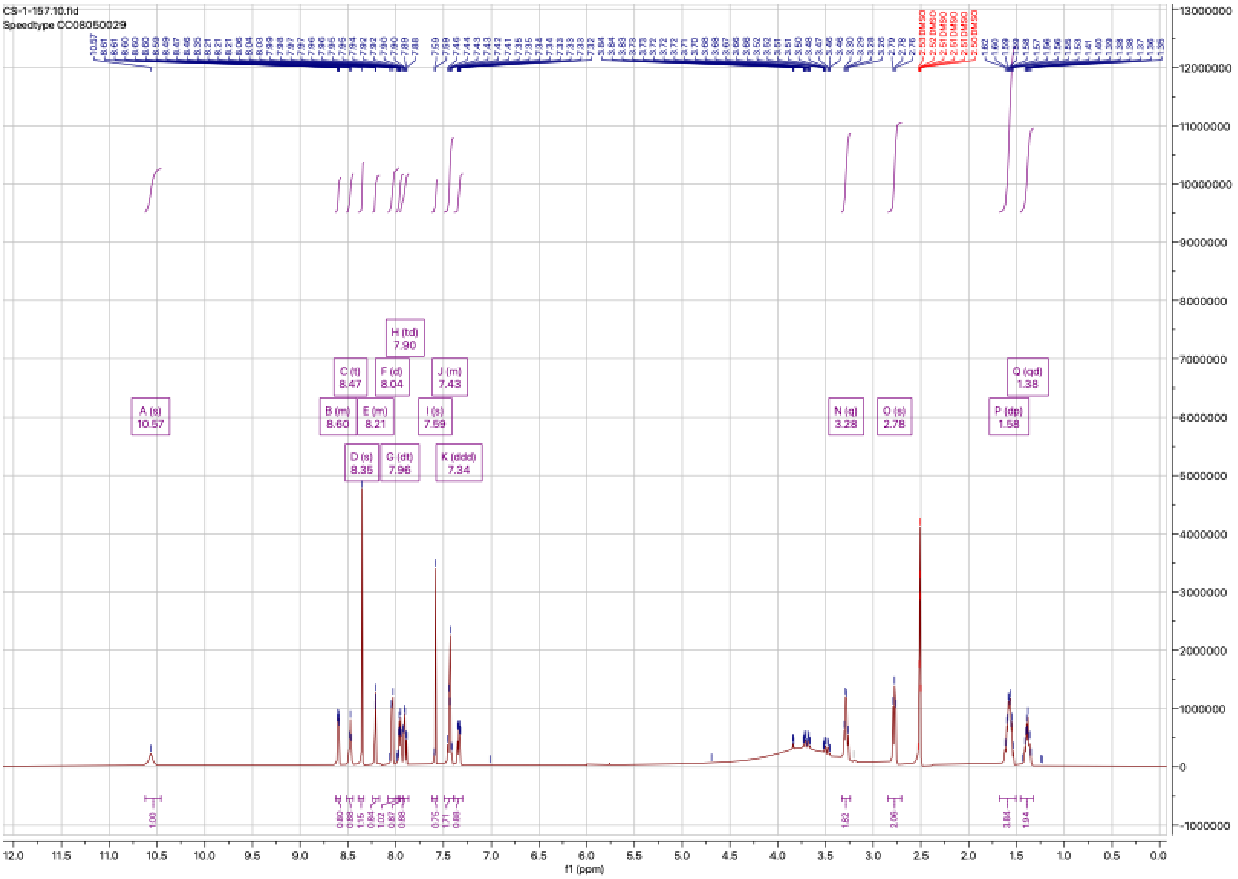

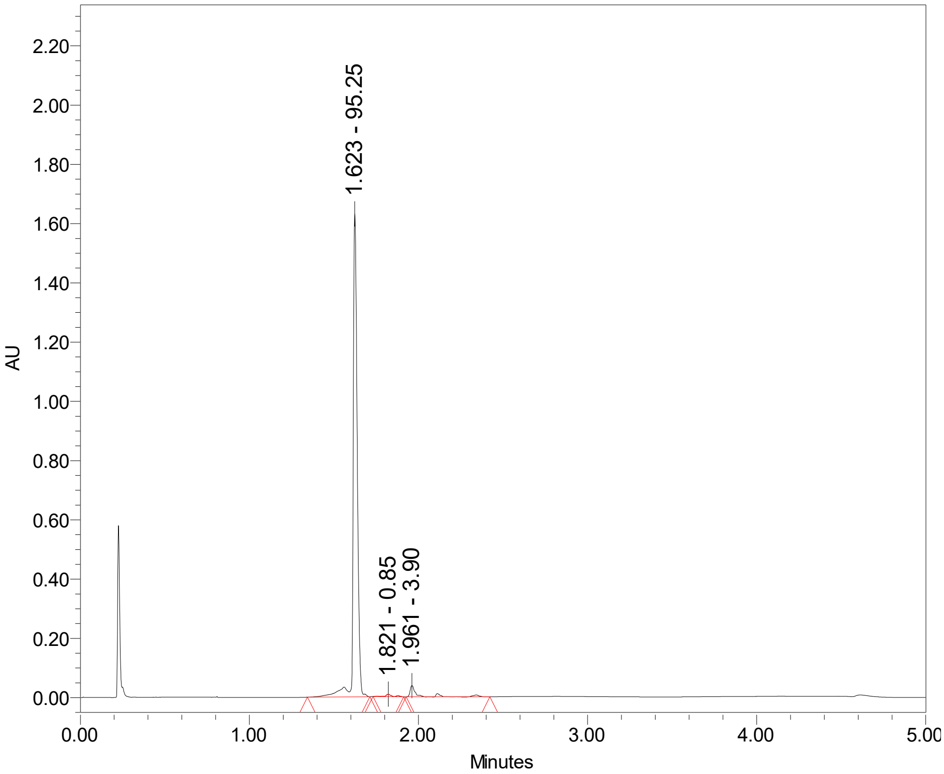

AU
2.20
2.00
1.80
1.60
1.40
1.20
1.00
0.80
0.60
0.40
0.20
0.00
1.623 - 95.25
1.821 - 0.85
1.961 - 3.90
0.00
1.00
2.00
3.00
4.00
5.00
Minutes

**CS-1-158:**

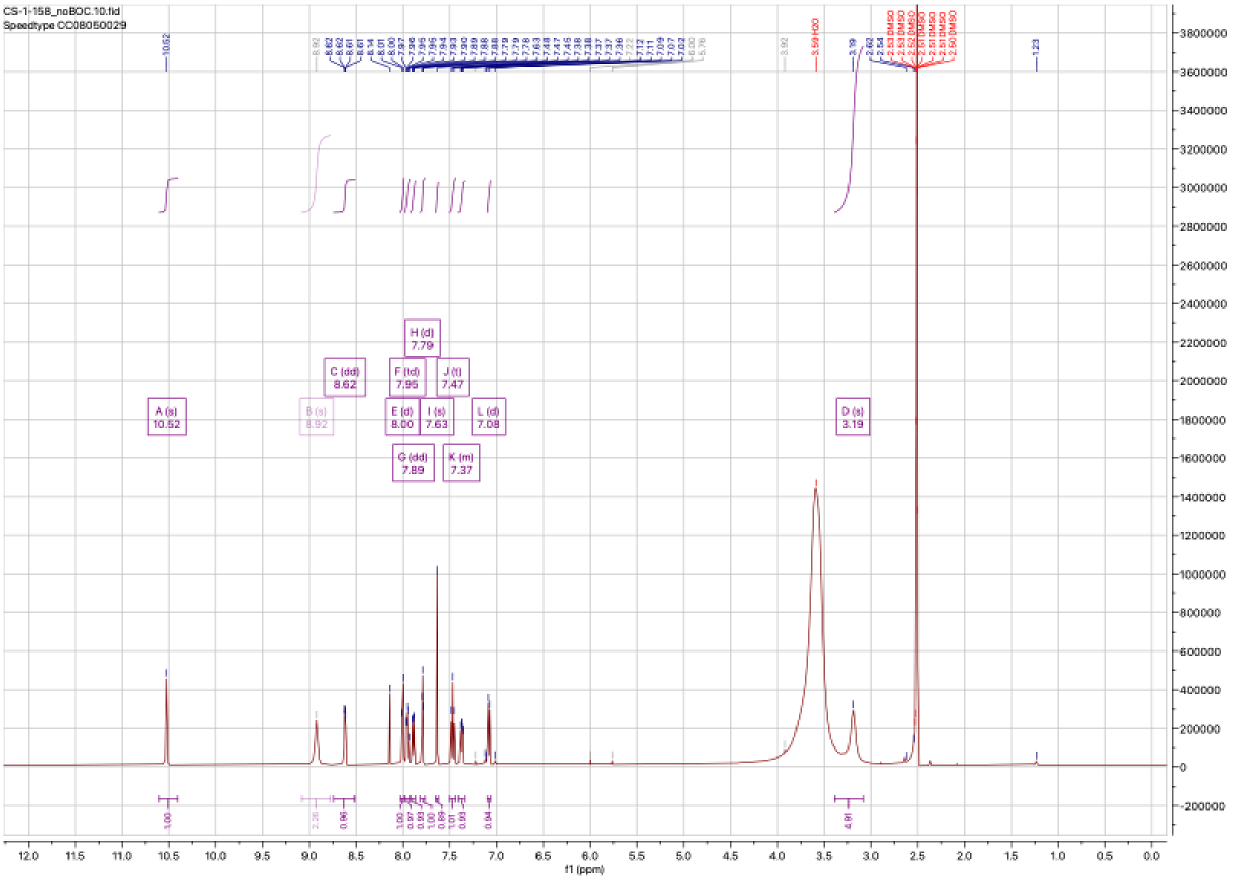

CS-1-158_noBOC.10.fid
Speedtype CC08050029
A (s)
10.52
B (s)
8.92
C (dd)
8.62
E (d)
8.00
F (td)
7.95
G (dd)
7.89
H (d)
7.79
I (s)
7.63
J (t)
7.47
K (m)
7.37
L (d)
7.08
D (s)
3.19
f1 (ppm)


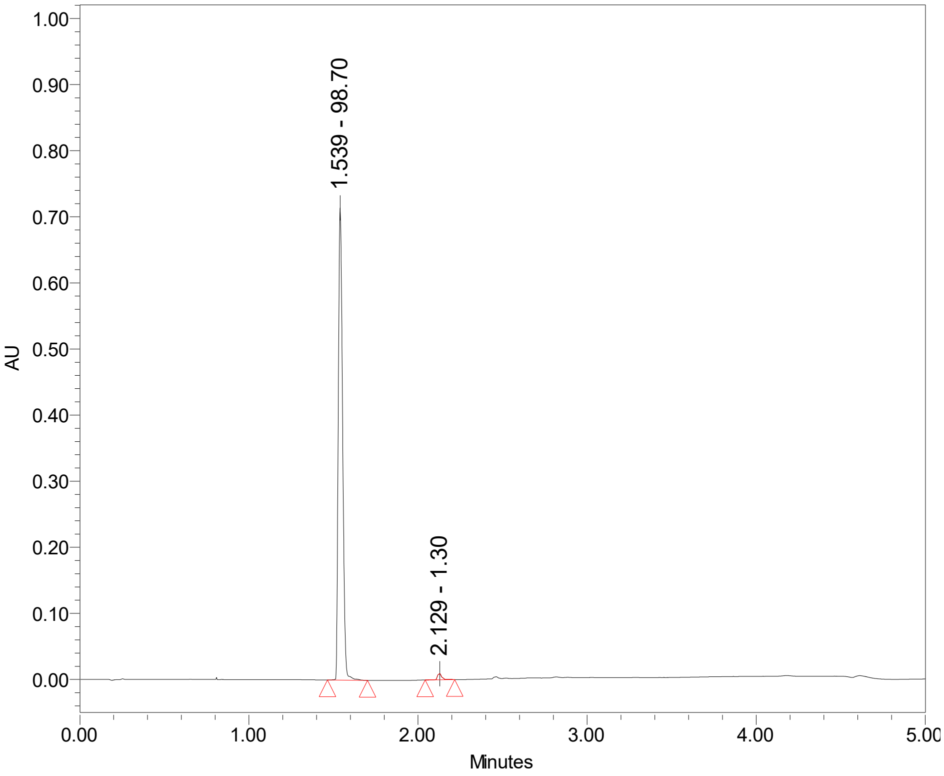

1.539 - 98.70
2.129 - 1.30
AU
Minutes

**CS-1-159:**

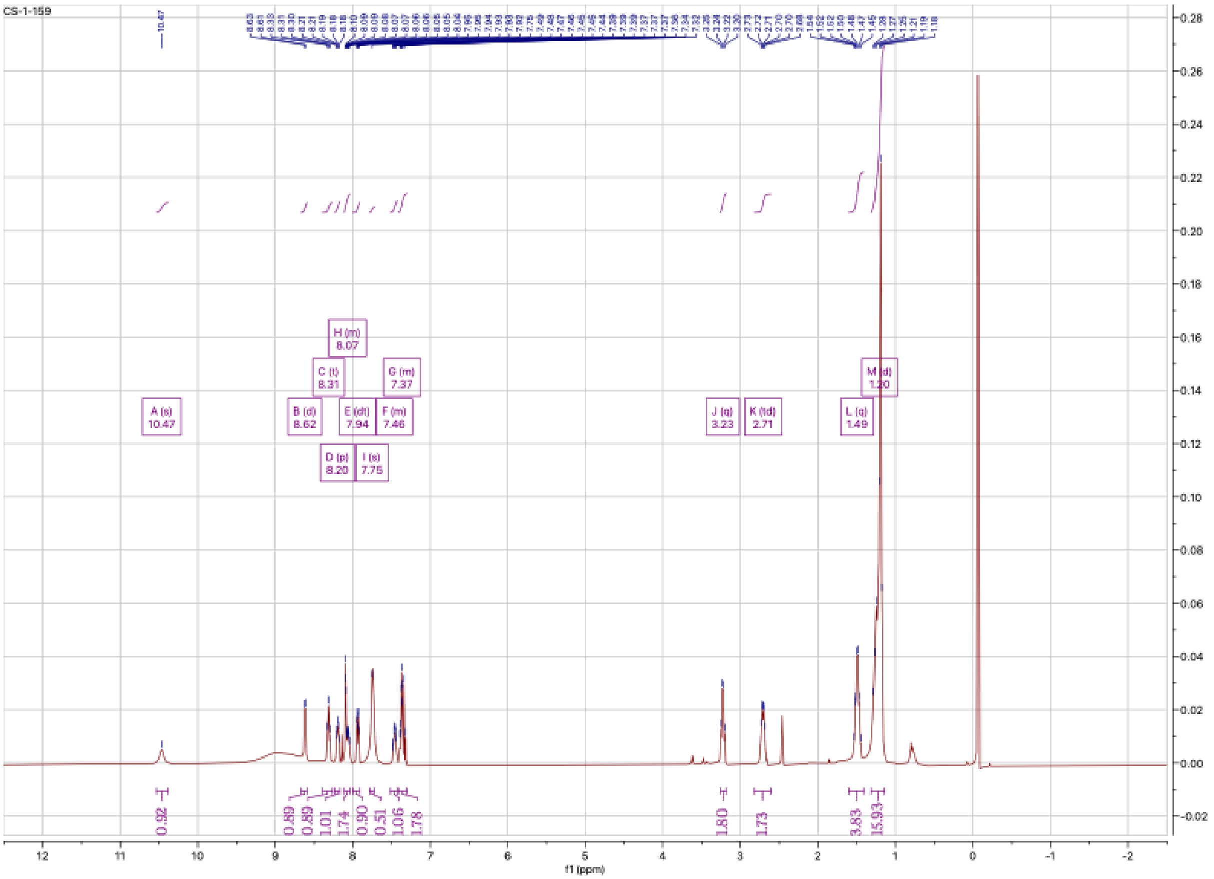

CS-1-159
A (s)
10.47
B (d)
8.62
C (t)
8.31
D (p)
8.20
E (dt)
7.94
F (m)
7.46
G (m)
7.37
H (m)
8.07
I (s)
7.75
J (q)
3.23
K (td)
2.71
L (q)
1.49
M (d)
1.20
f1 (ppm)


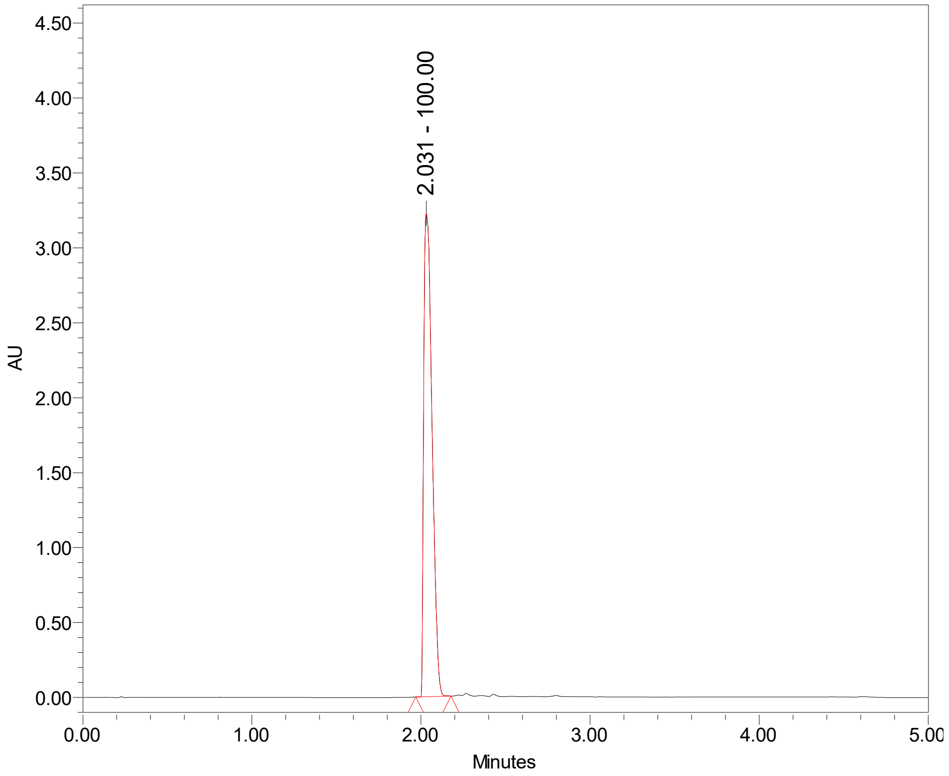

2.031 - 100.00
AU
Minutes

**CS-1-160:**

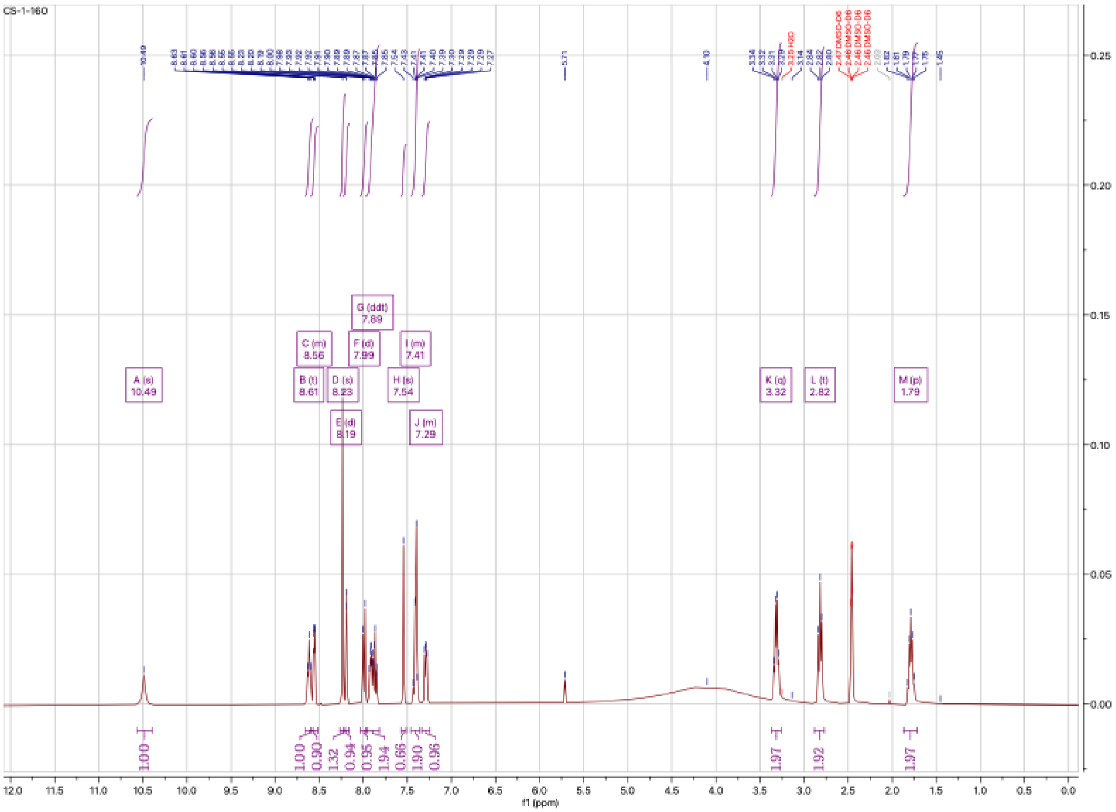
CS-1-160
A (s) 10.49
B (t) 8.61
C (m) 8.56
D (s) 8.23
E (d) 8.19
F (d) 7.99
G (ddt) 7.89
H (s) 7.54
I (m) 7.41
J (m) 7.29
K (q) 3.32
L (t) 2.82
M (p) 1.79
1.00
1.00
0.90
1.32
0.94
0.95
1.94
0.66
1.90
0.96
1.97
1.92
1.97
f1 (ppm)

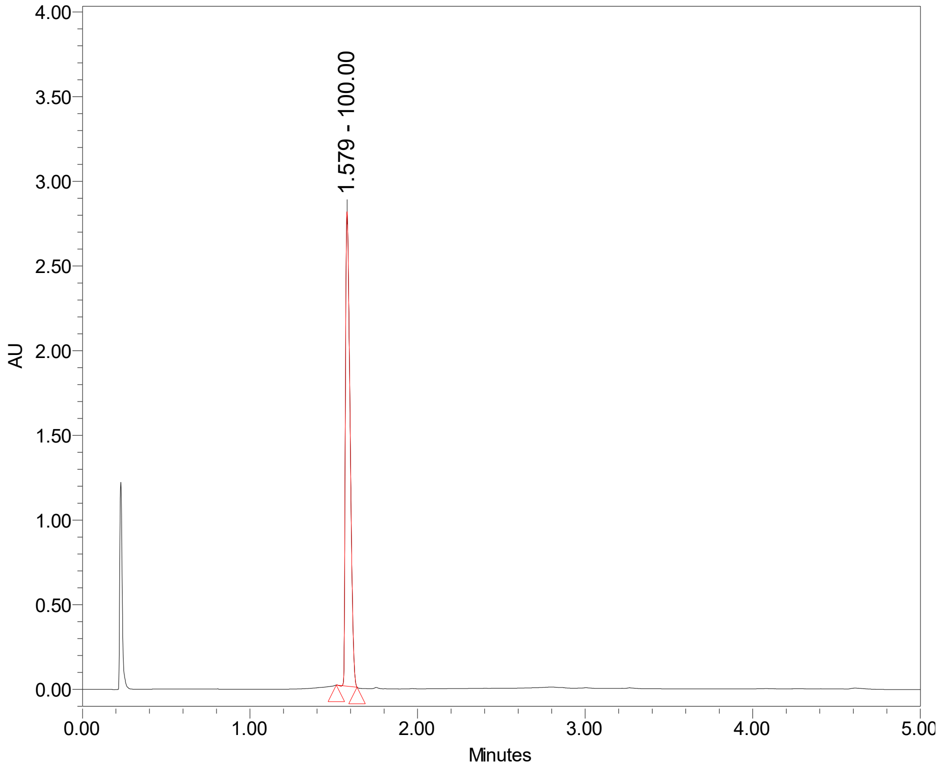
1.579 - 100.00
AU
Minutes

# CS-1-161:

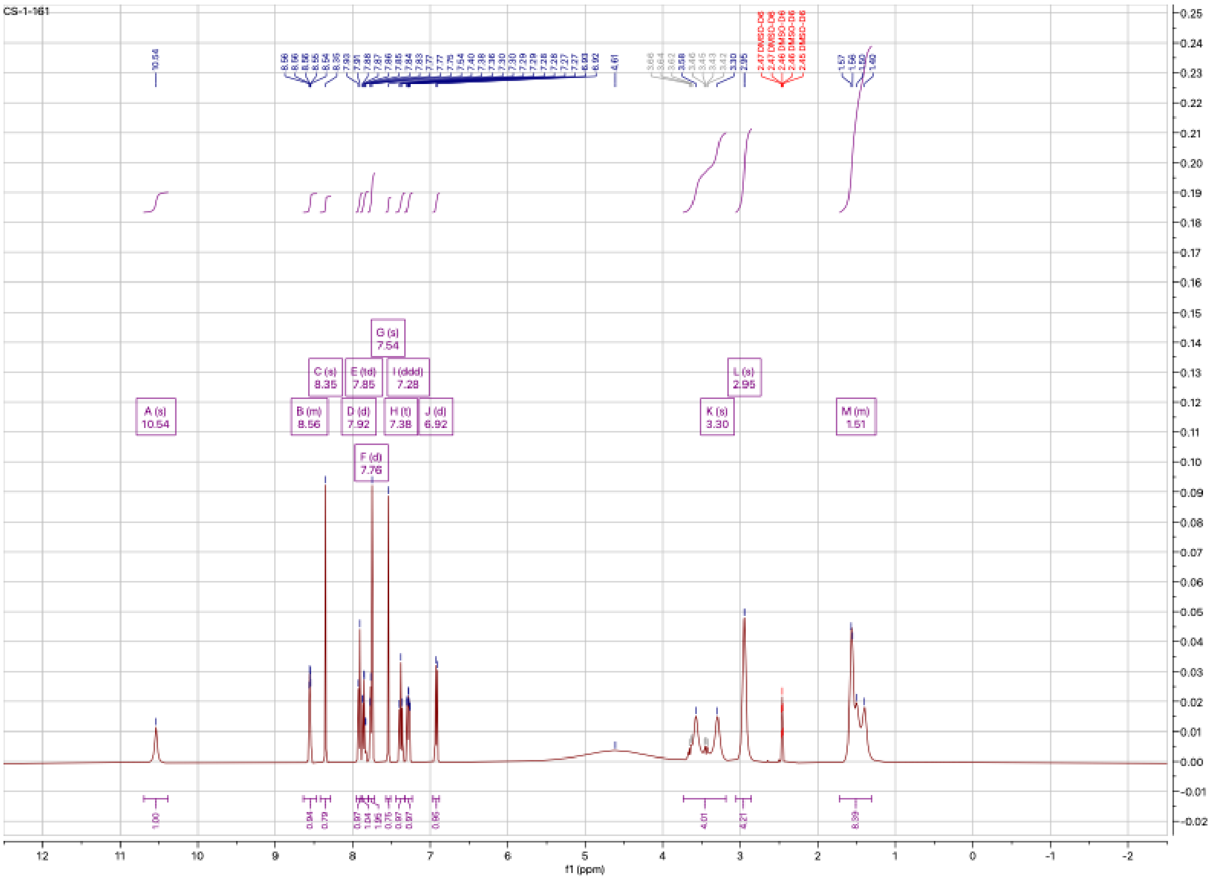

CS-1-161
A (s)
10.54
B (m)
8.56
C (s)
8.35
D (d)
7.92
E (td)
7.85
F (d)
7.76
G (s)
7.54
H (t)
7.38
I (ddd)
7.28
J (d)
6.92
K (s)
3.30
L (s)
2.95
M (m)
1.51
f1 (ppm)

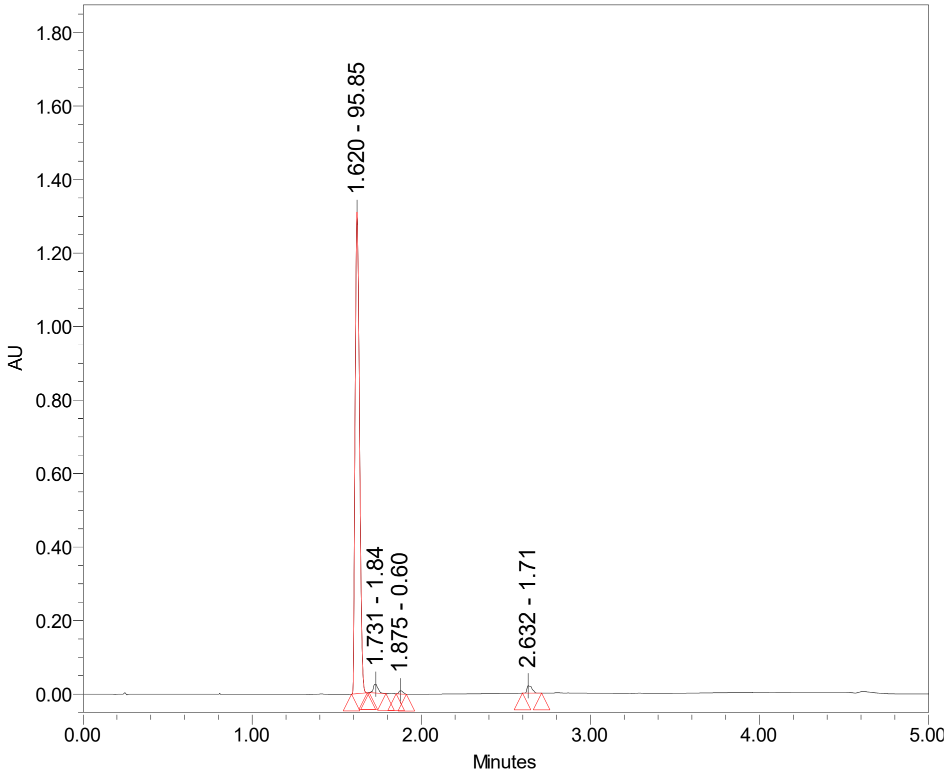

1.620 - 95.85
1.731 - 1.84
1.875 - 0.60
2.632 - 1.71
AU
1.80
1.60
1.40
1.20
1.00
0.80
0.60
0.40
0.20
0.00
0.00
1.00
2.00
3.00
4.00
5.00
Minutes

**CS-1-162**

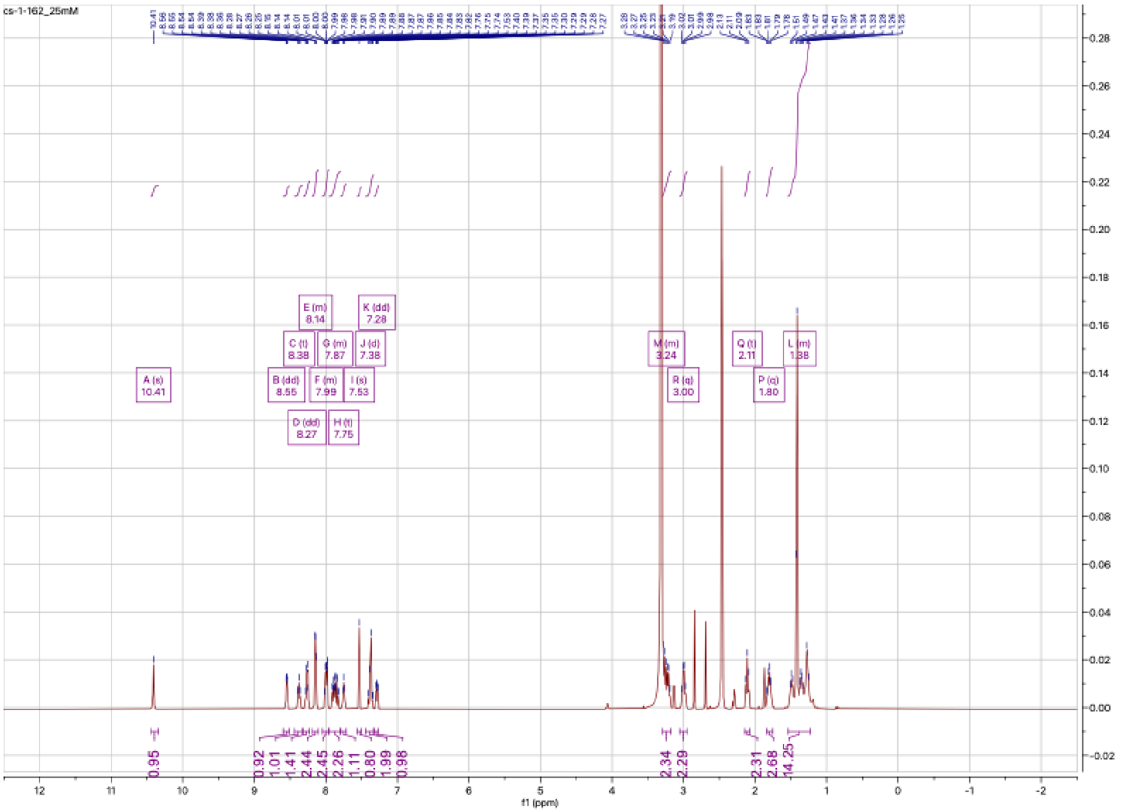
cs-1-162_25mM
A (s) 10.41
B (dd) 8.55
C (t) 8.38
D (dd) 8.27
E (m) 8.14
F (m) 7.99
G (m) 7.87
H (t) 7.75
I (s) 7.53
J (d) 7.38
K (dd) 7.28
M (m) 3.24
R (q) 3.00
Q (t) 2.11
P (q) 1.80
L (m) 1.38
f1 (ppm)

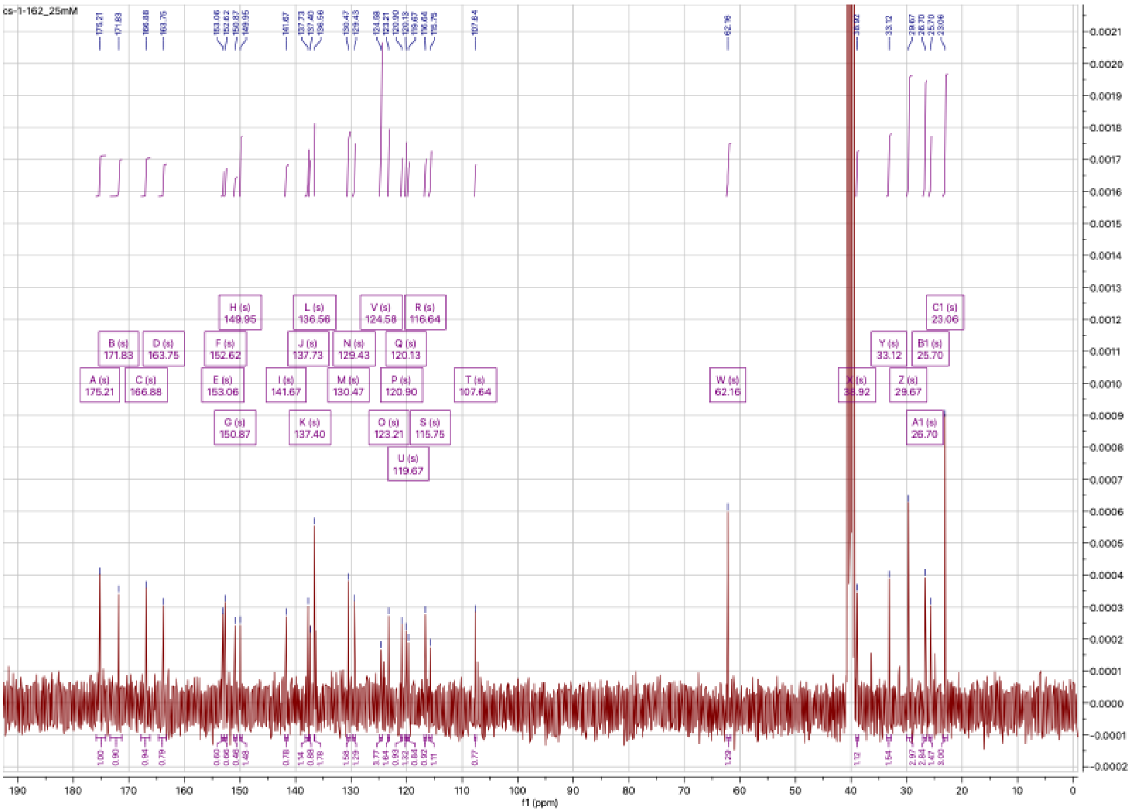
cs-1-162_25mM
A (s) 175.21
B (s) 171.83
C (s) 166.88
D (s) 163.75
E (s) 153.06
F (s) 152.62
G (s) 150.87
H (s) 149.95
I (s) 141.67
J (s) 137.73
K (s) 137.40
L (s) 136.56
M (s) 130.47
N (s) 129.43
O (s) 123.21
V (s) 124.58
P (s) 120.90
Q (s) 120.13
U (s) 119.67
R (s) 116.64
S (s) 115.75
T (s) 107.64
W (s) 62.16
X (s) 38.92
Y (s) 33.12
Z (s) 29.67
A1 (s) 26.70
B1 (s) 25.70
C1 (s) 23.06
f1 (ppm)

## CS-1-163:

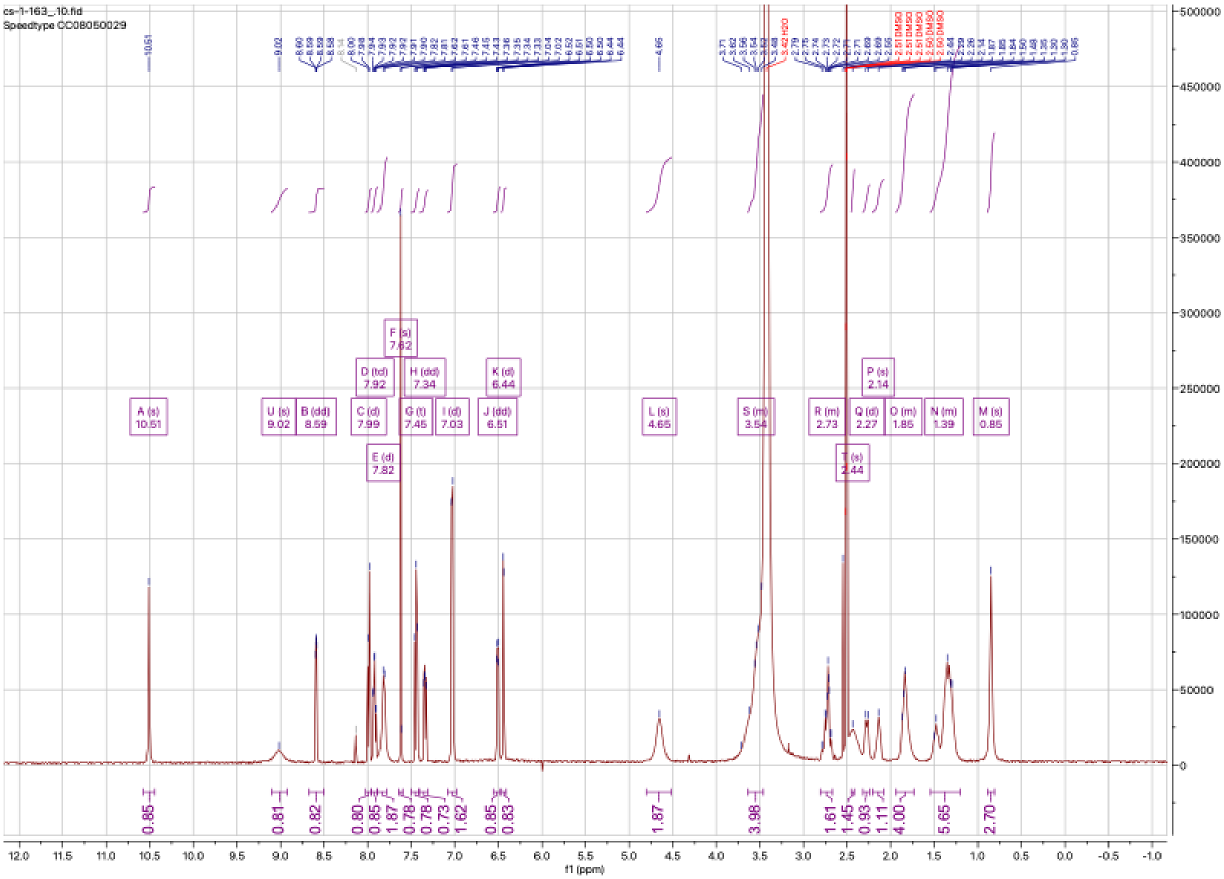
cs-1-163_.10.fid
Speedtype CC08050029
A (s) 10.51
U (s) 9.02
B (dd) 8.59
C (d) 7.99
D (td) 7.92
E (d) 7.82
F (s) 7.62
G (t) 7.45
H (dd) 7.34
I (d) 7.03
J (dd) 6.51
K (d) 6.44
L (s) 4.65
S (m) 3.54
R (m) 2.73
T (s) 2.44
Q (d) 2.27
P (s) 2.14
O (m) 1.85
N (m) 1.39
M (s) 0.85
f1 (ppm)

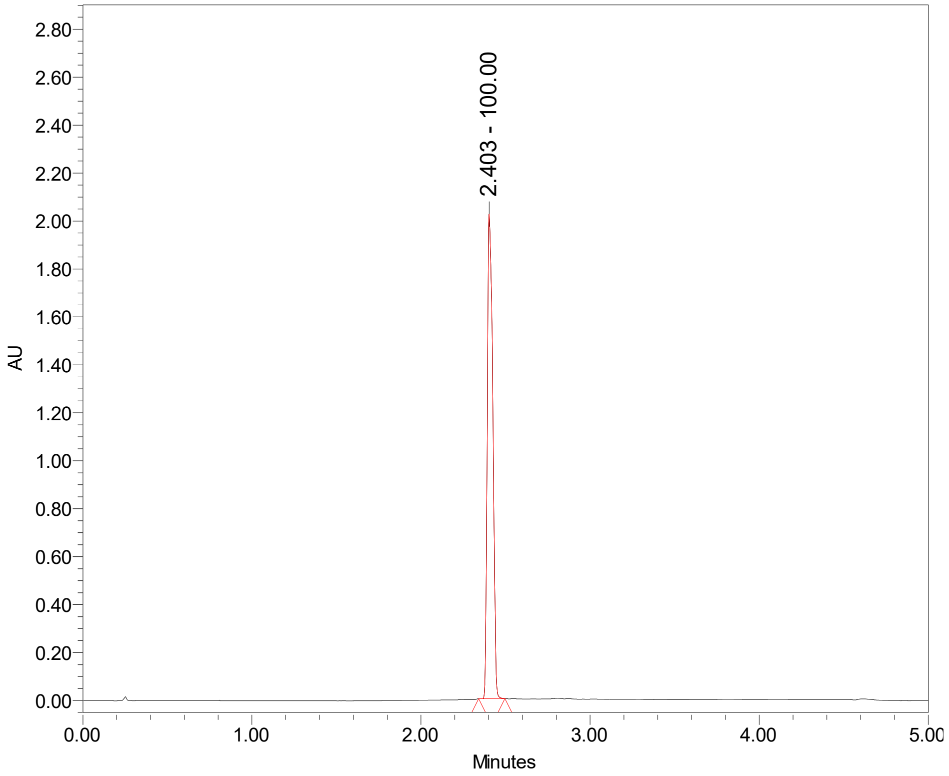
2.403 - 100.00
AU
Minutes

## CS-1-164:

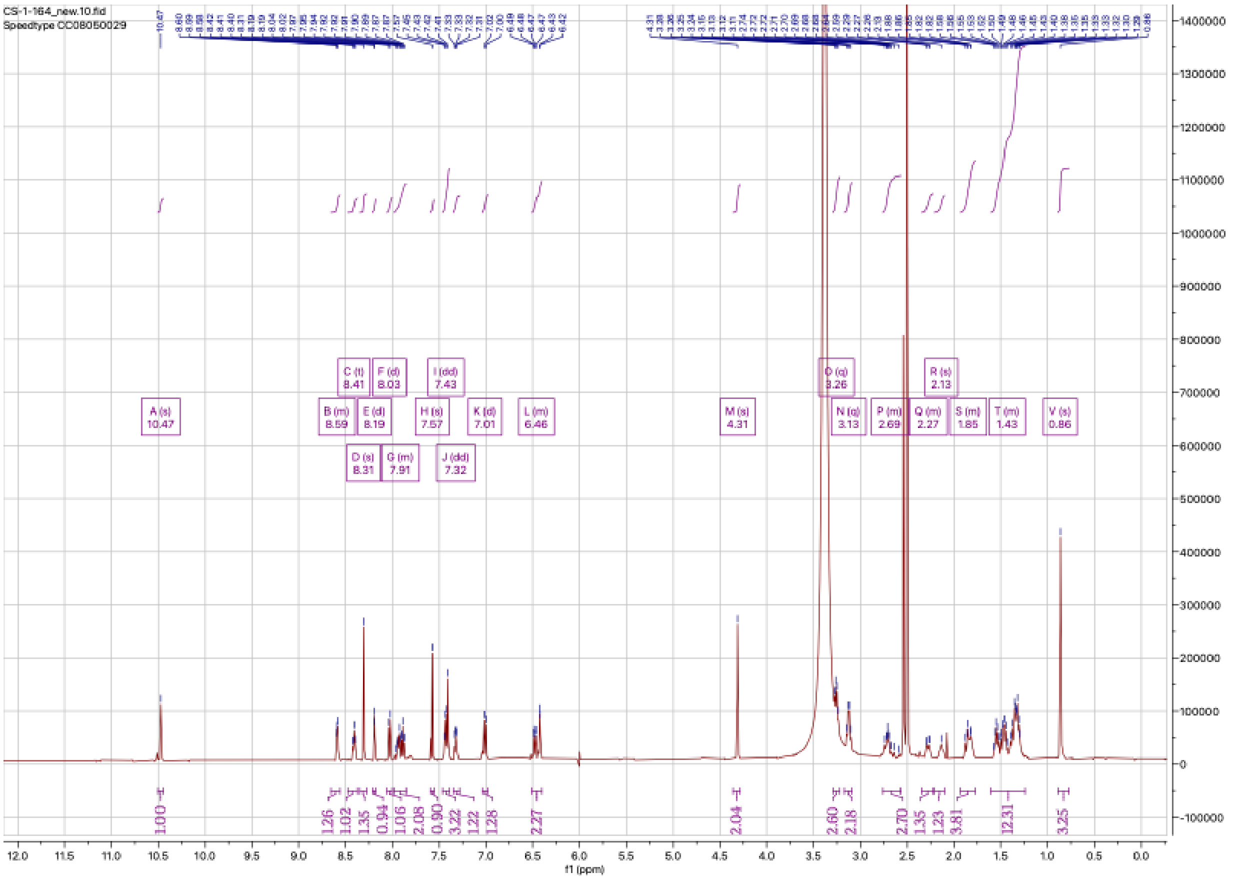
CS-1-164_new.10.fid
Speedtype CC08050029
f1 (ppm)

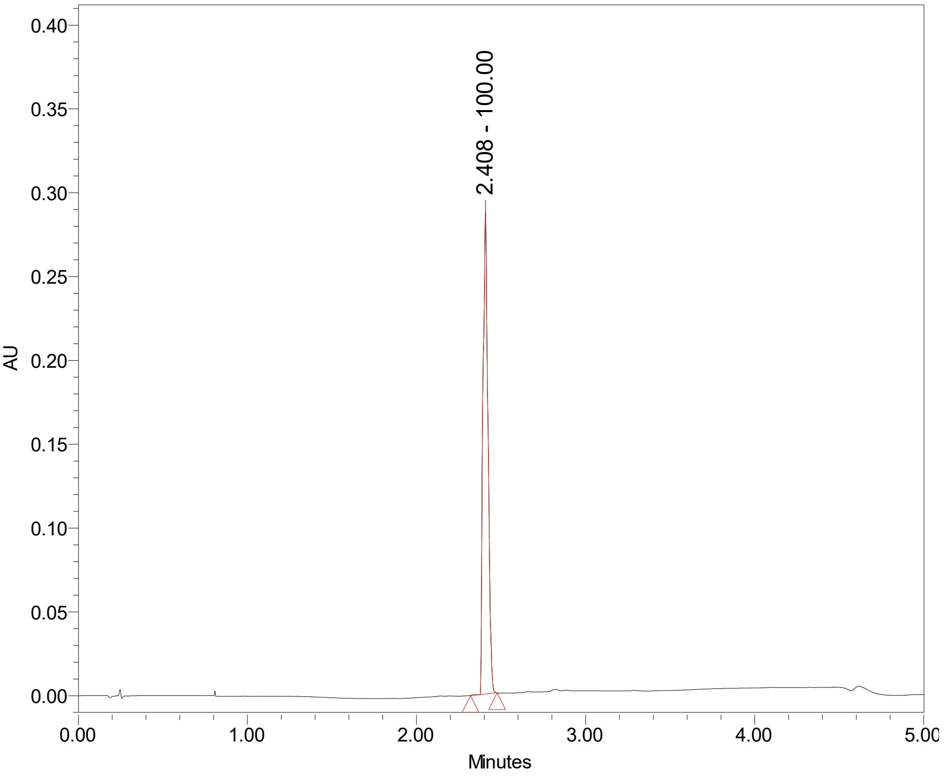
2.408 - 100.00
AU
Minutes

**CS-1-165:**

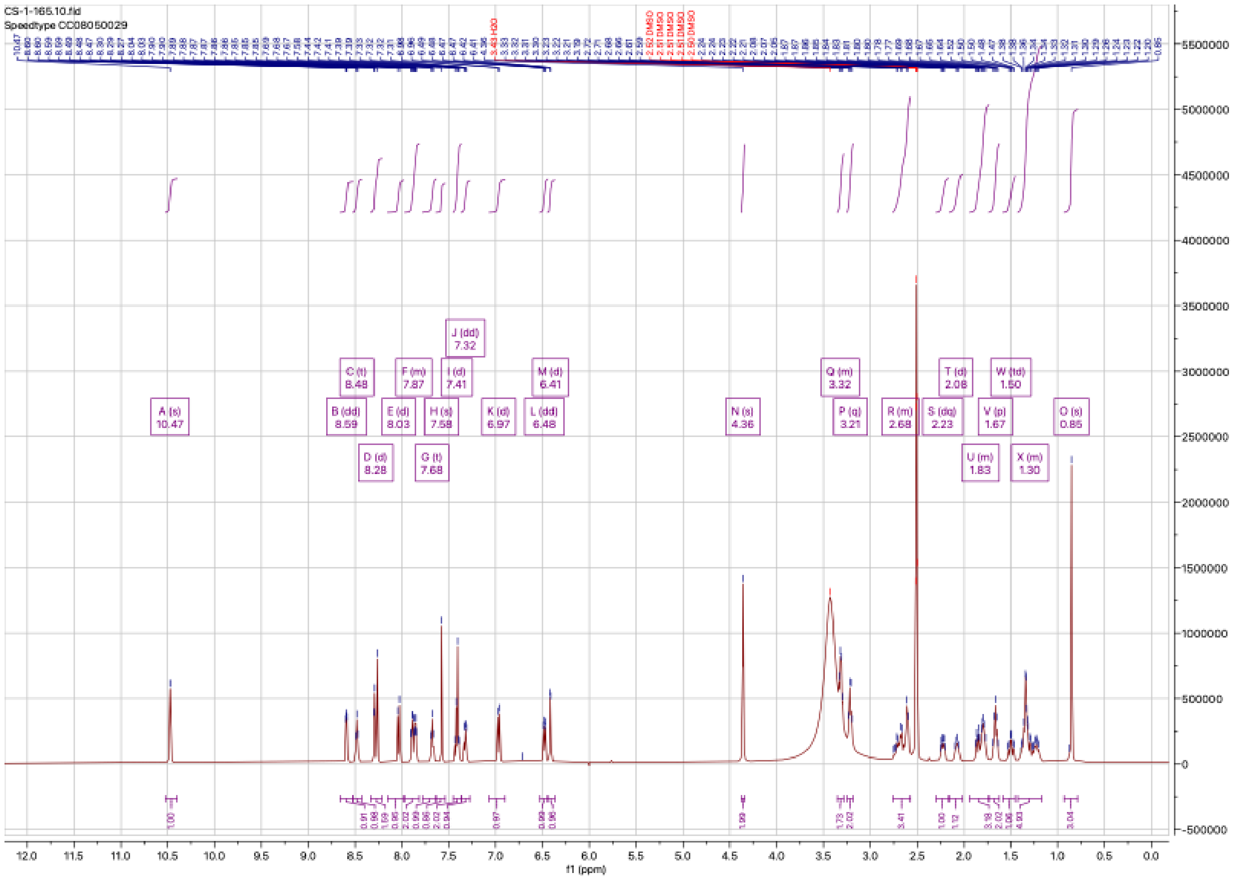

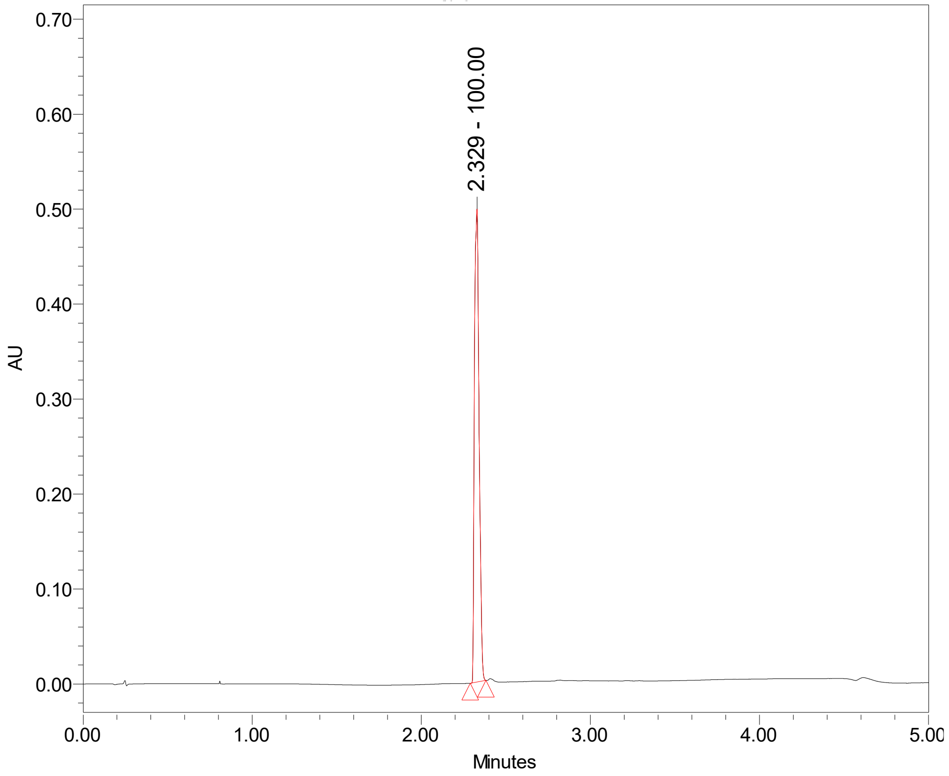

**CS-1-166:**

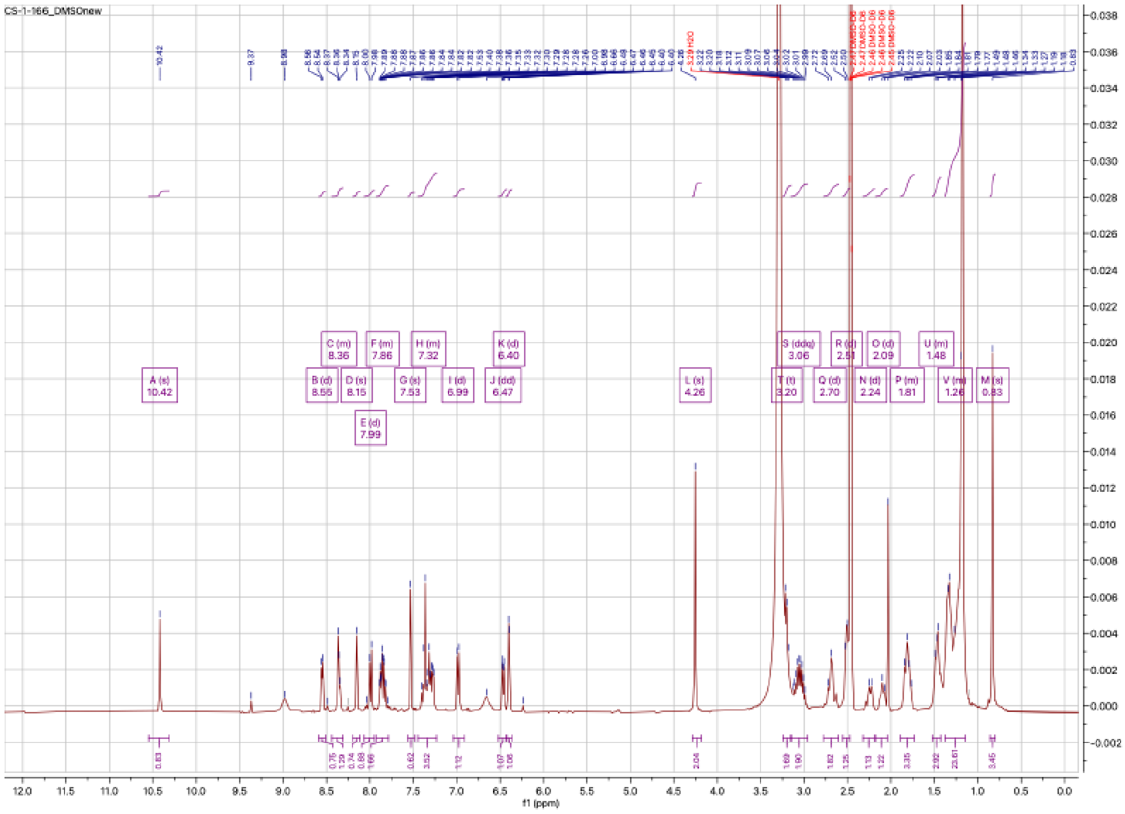
CS-1-166_DMSOnew
A (s) 10.42
B (d) 8.55
C (m) 8.36
D (s) 8.15
E (d) 7.99
F (m) 7.86
G (s) 7.53
H (m) 7.32
I (d) 6.99
J (dd) 6.47
K (d) 6.40
L (s) 4.26
T (t) 3.20
S (ddq) 3.06
Q (d) 2.70
R (d) 2.51
N (d) 2.24
O (d) 2.09
P (m) 1.81
U (m) 1.48
V (m) 1.26
M (s) 0.83
f1 (ppm)

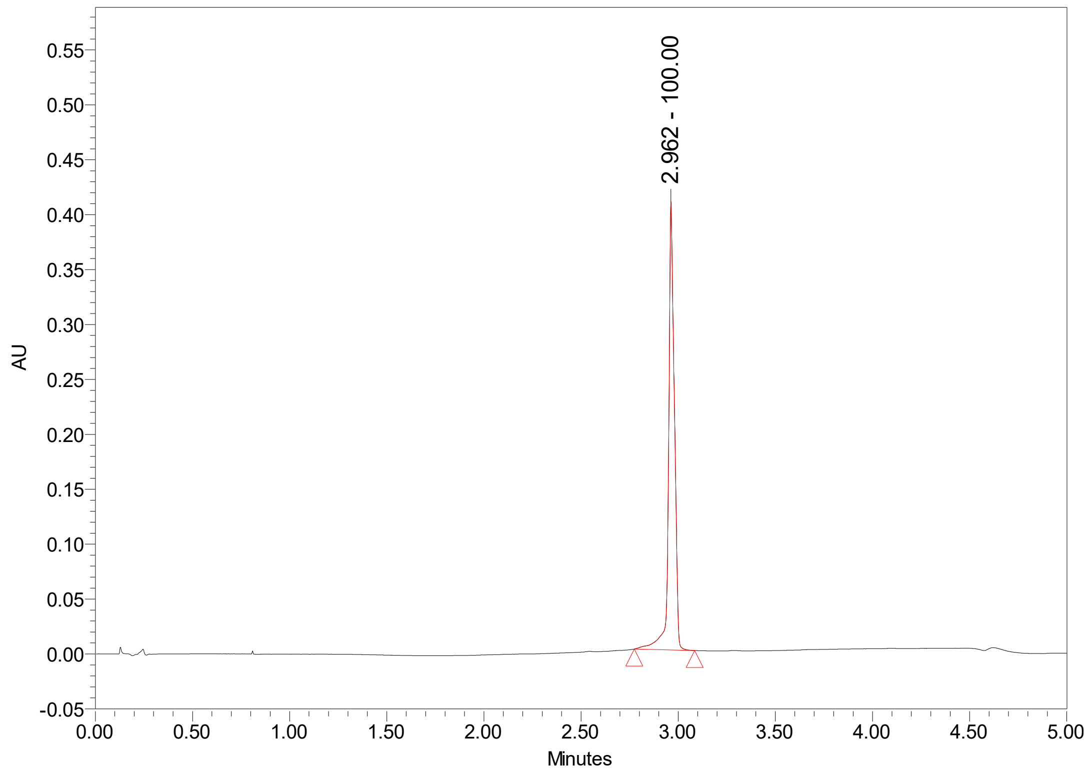
2.962 - 100.00
AU
Minutes

# CS-1-167:

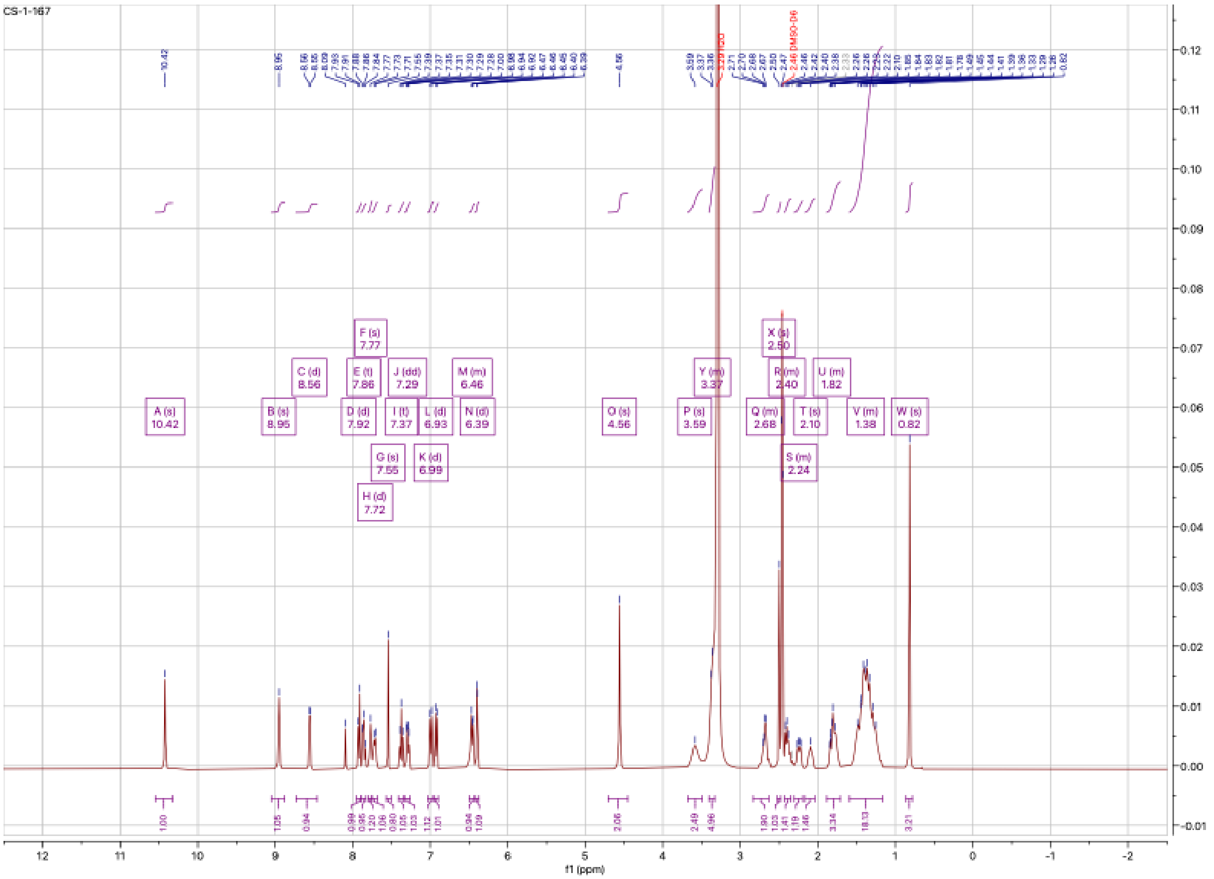

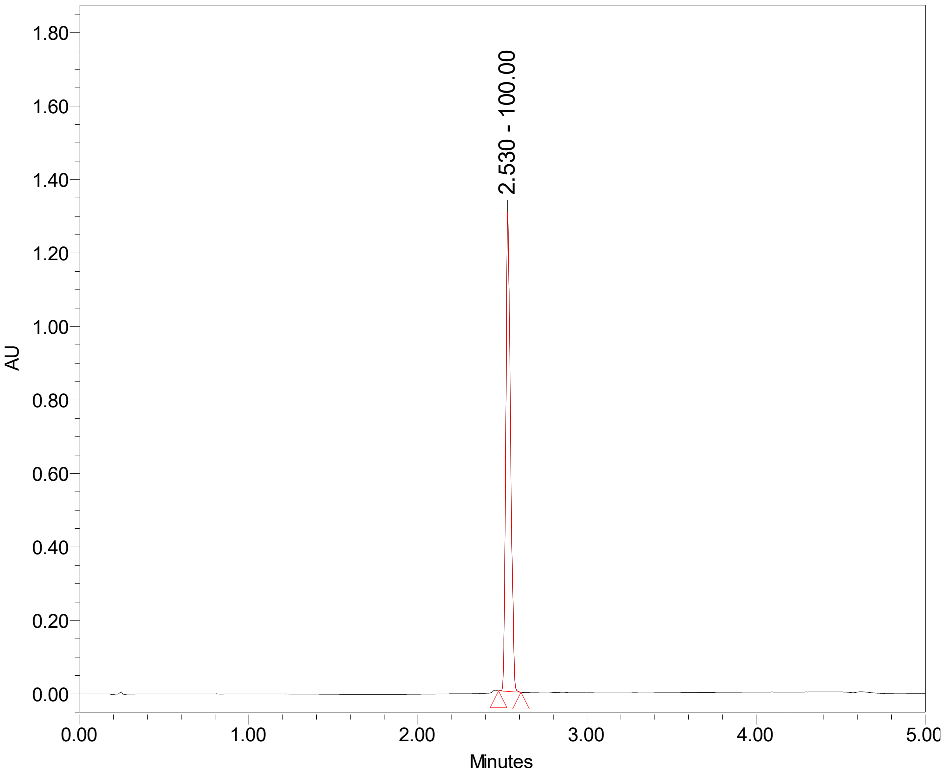

2.530 - 100.00
AU
Minutes

**CS-1-168**

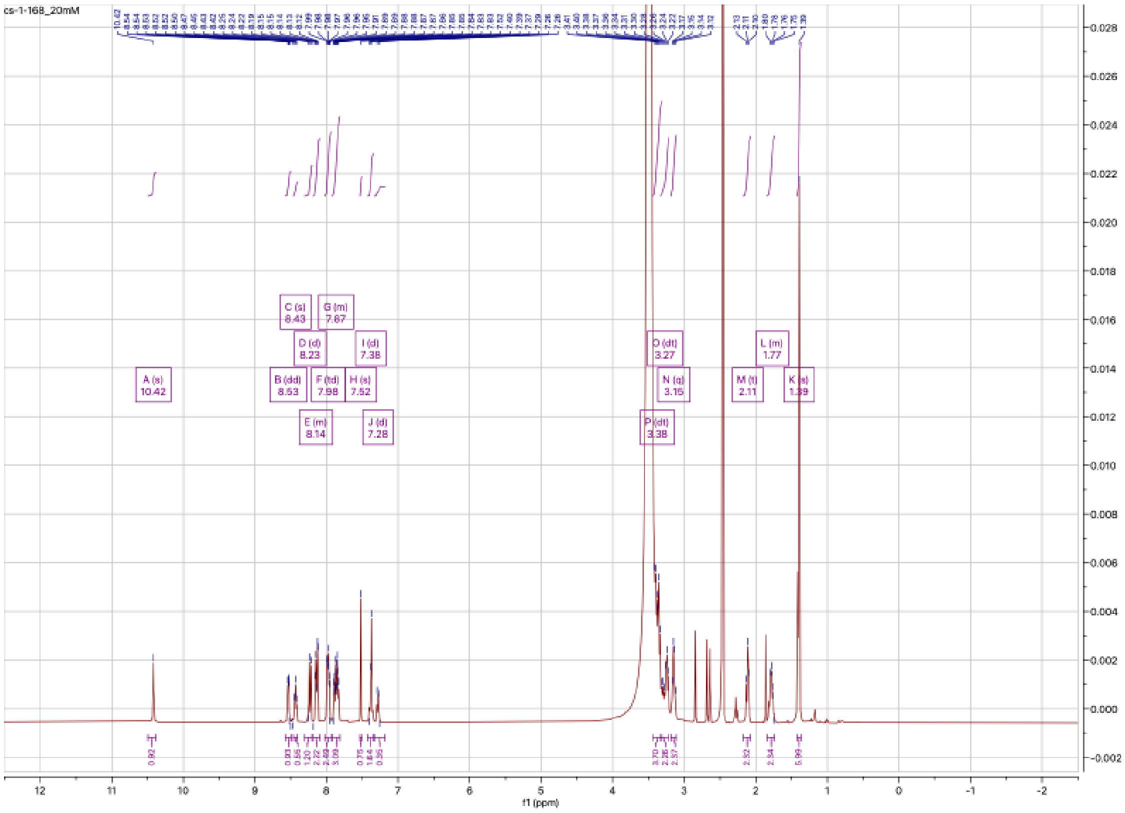

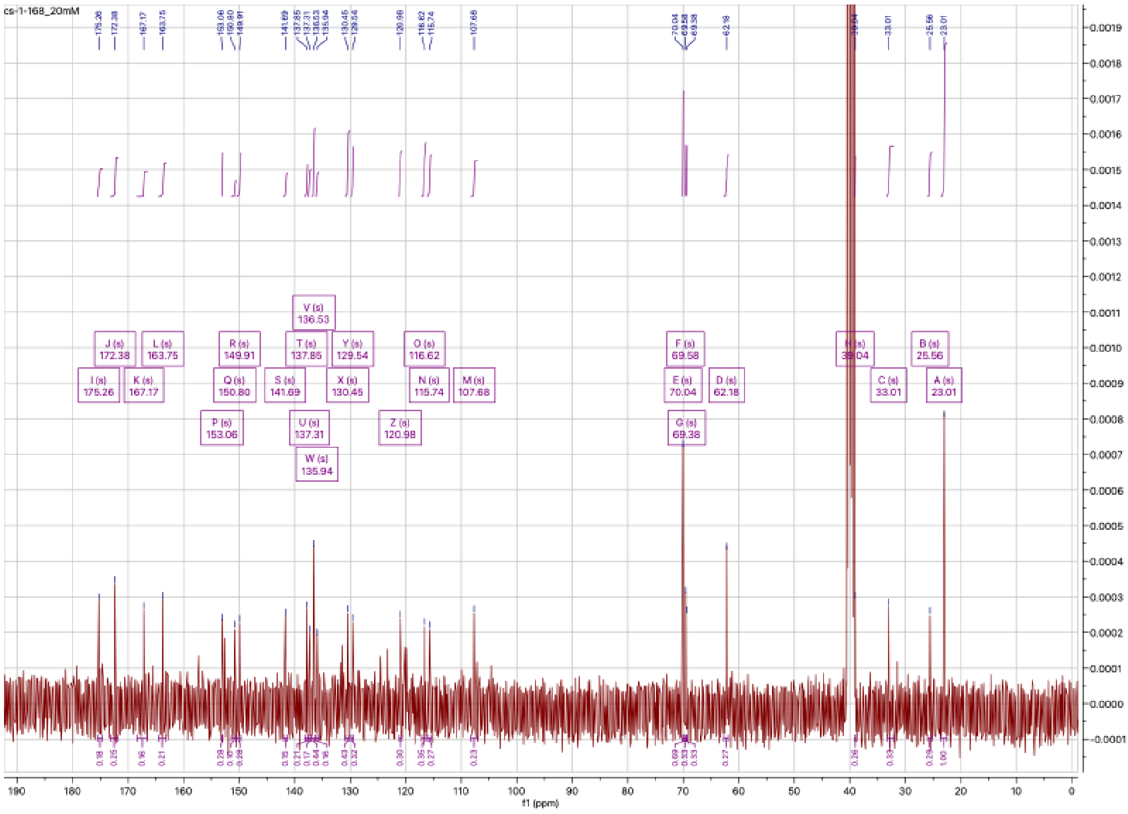

## CS-1-169

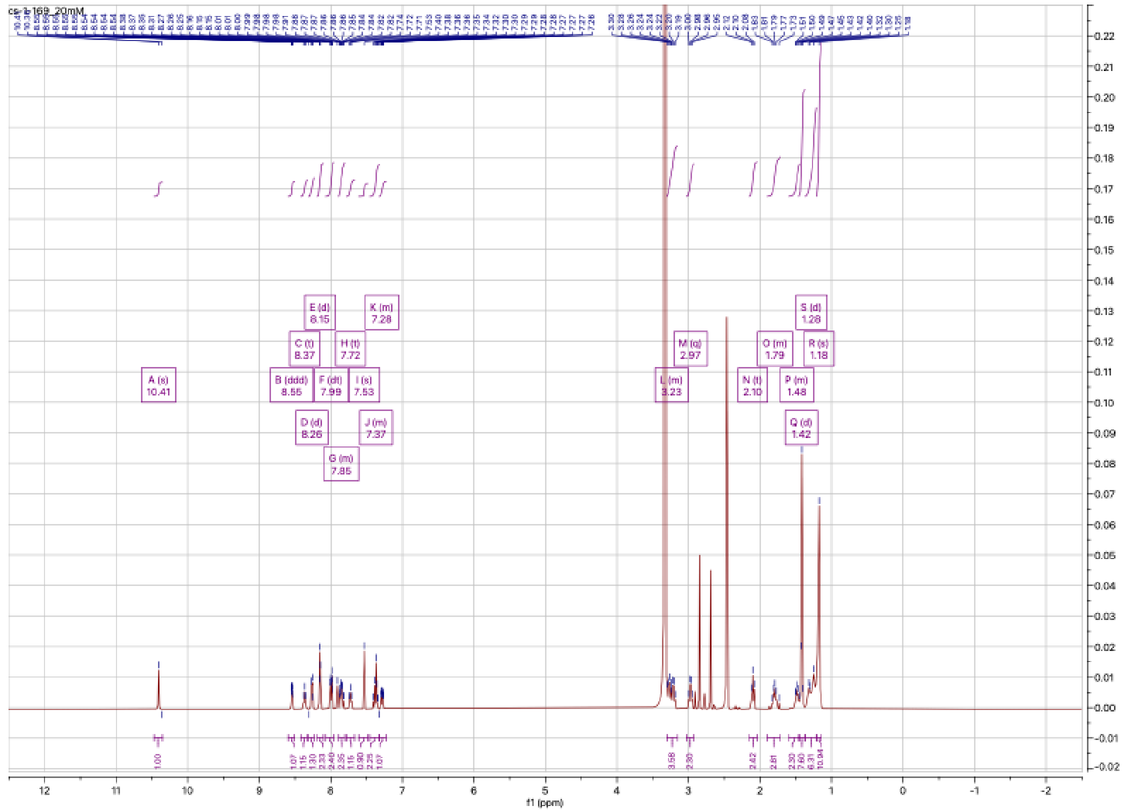

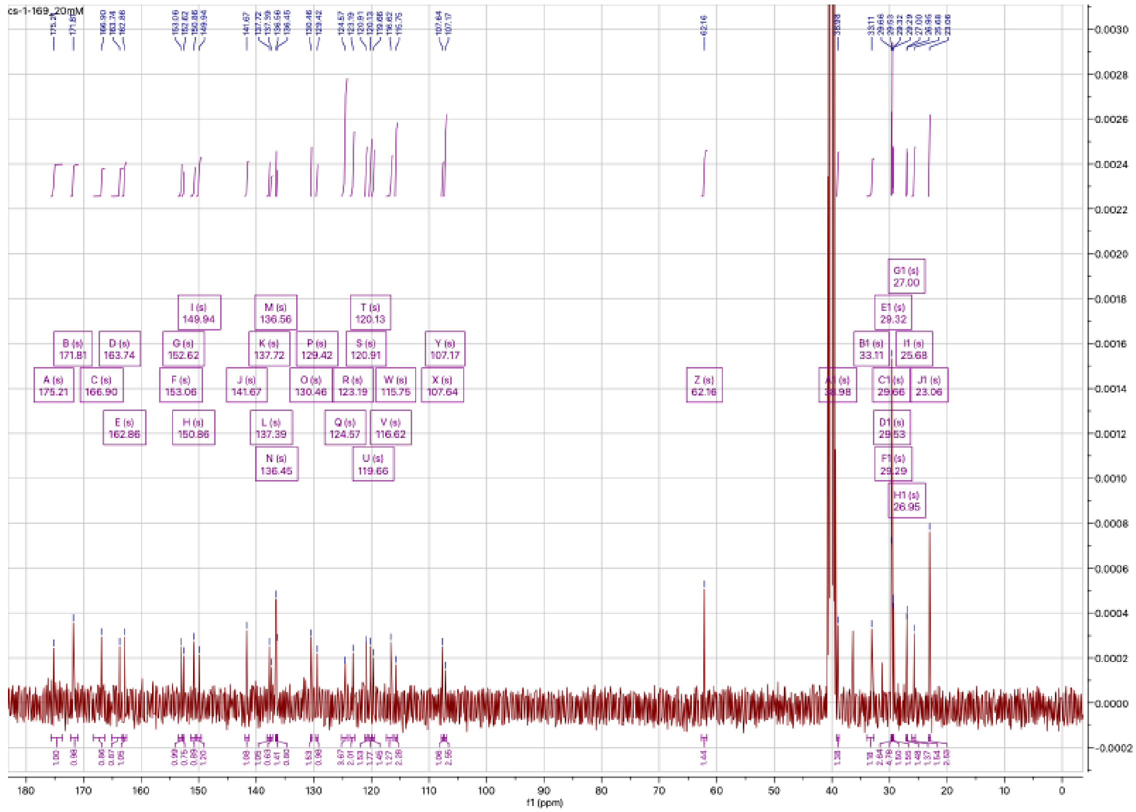

**CS-1-175**:

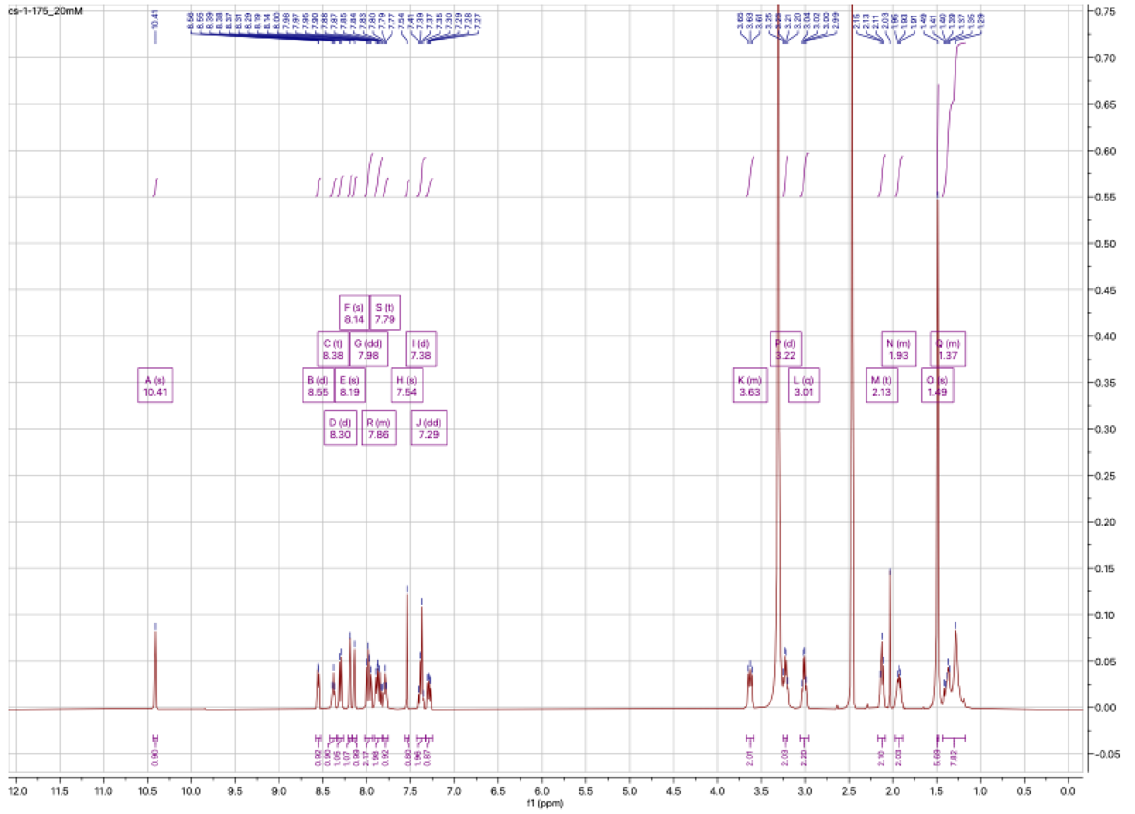

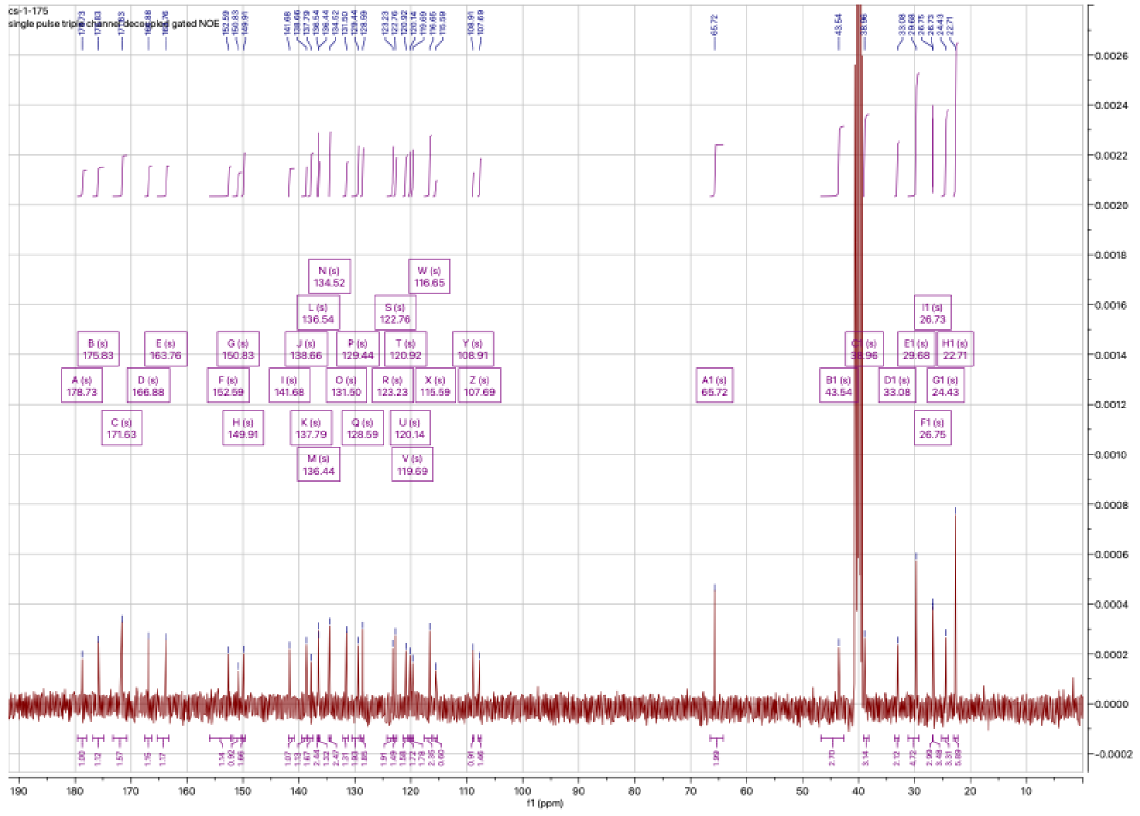

# CS-1-176

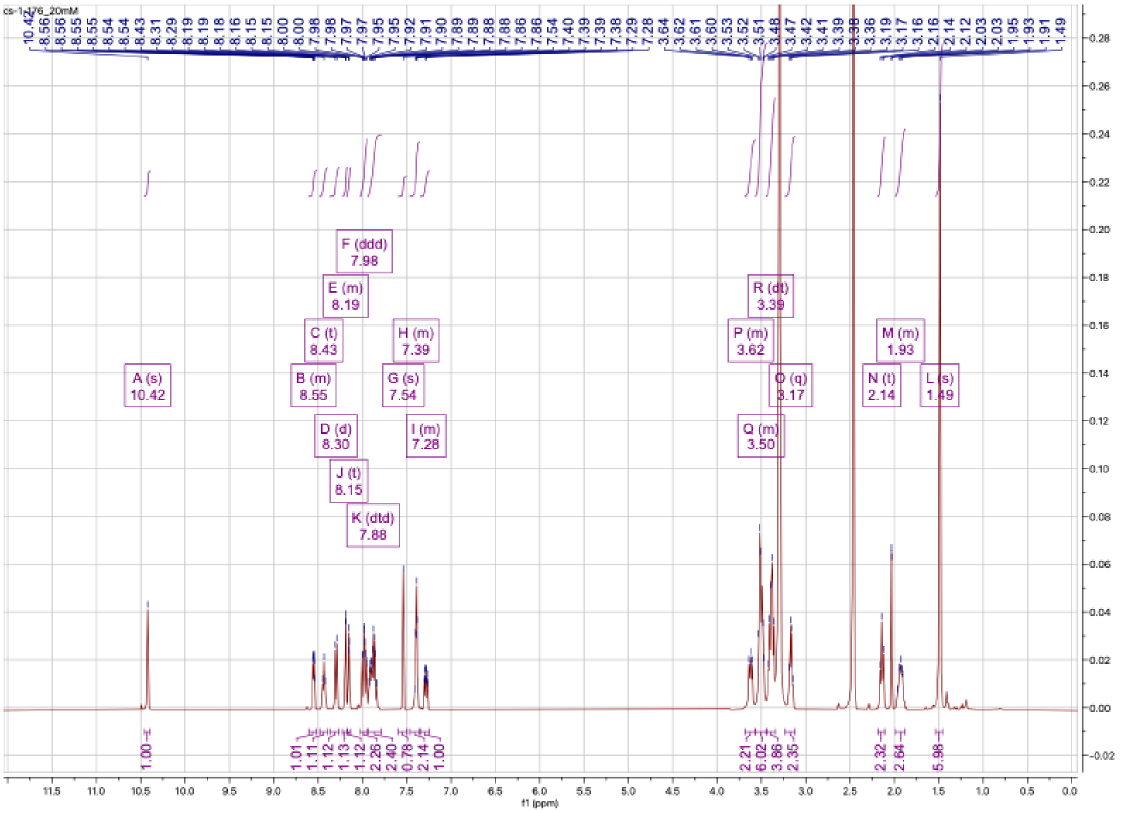

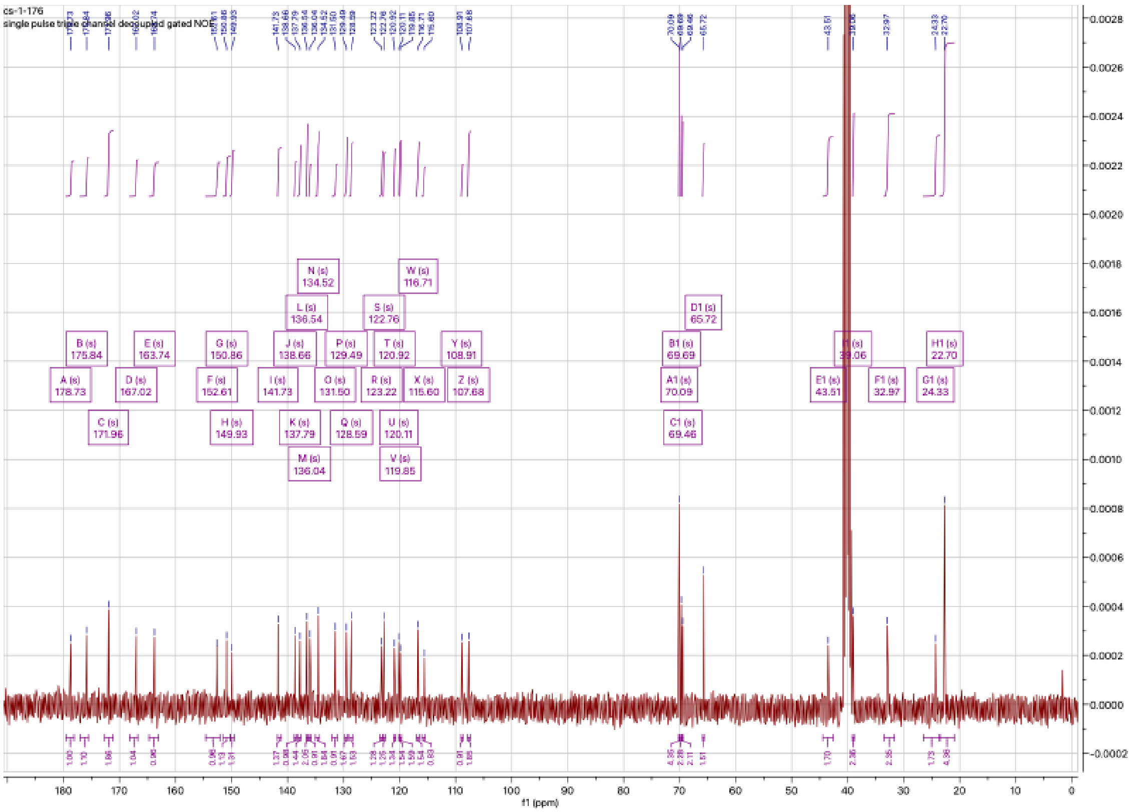
